# From Language-Cognition Coupling to AI Foundations: An Evolutionary Perspective on 13,415 SNVs in 33 Language/Cognition Genes

**DOI:** 10.64898/2026.08.19.745865

**Authors:** Zhizhou Zhang, Yongdong Xu

## Abstract

This study aims to quantify the genetic similarity of different species (from fish to humans) to the human reference genome (pp6, Homo sapiens.GRCh38) based on the allele presence/absence patterns of 13,415 SNV loci from 33 language/cognition-related genes, identify key breakpoints during evolution, and evaluate the enrichment of language and cognition genes at these breakpoints. We designed a similarity calculation method relying on binary features (four columns for A/T/C/G), adopted five difference/distance measures (Sørensen, Rogers, Nei, Reynolds, and Hellinger), and converted them into similarity values (1/(1+distance)). For each method, samples were independently ranked, the first derivative of similarity was computed, and the top 12 peaks were selected as candidate breakpoints. Results show that the similarity curves from the five methods are highly consistent (correlation coefficients >0.9), with major peaks concentrated at positions 355, 363, 381, 382, 390, 400, etc., where the corresponding samples are predominantly ancient hominins and primates. Furthermore, we defined 13 peak groups (starting positions 355–401). For each peak within a group, pairwise SNV differences between the peak apex sample and its immediate left neighbor were compared, and the intersection F_INTERSECTION (shared differential loci) was obtained. For each F_INTERSECTION, we calculated the proportions of language genes and cognition genes. In addition, we computed the differential sets between adjacent groups’ F_INTERSECTION to trace the gradual emergence of new loci. In F_INTERSECTION, language genes accounted for an average of 59.5%, and cognition genes for an average of 62.9%. The proportion of language genes reached a peak at position 383 (61.2%), while cognition genes peaked at position 386 (64.9%). High-frequency peak samples include c25, c27, and ja2, suggesting that language-cognition genes may have undergone independent intensification during Eurasian evolution. Differential analysis between adjacent F_INTERSECTION revealed a stepwise acquisition of new loci from position 355 to 401, with three bursts of newly added loci along the entire evolutionary axis.

These molecular patterns carry implications beyond evolutionary genetics. The absence of robust enrichment for either language or cognition genes at any breakpoint is compatible with the hypothesis that language and cognition are not separable functional modules, but rather constitute an integrated symbolic-cognitive system. This distinction matters for artificial intelligence research: if language, understood as an internal symbolic system, is inseparable from cognition, then treating large language models’ linguistic performance as a direct proxy for cognitive capacity may be theoretically problematic. The present study therefore provides not only a quantitative evolutionary framework based on similarity curves, but also a methodological caution for AI foundations—namely, that the competence–performance distinction must be carefully preserved when interpreting what large language models can and cannot do. The results highlight the potential importance of East Asian archaic hominins in the evolution of language-cognition genes, and suggest that future AI research may benefit from distinguishing between the mastery of deep symbolic regularities (closer to I-language) and the imitation of surface statistical patterns (closer to externalized behavior).

## 1 Introduction

There are essential differences between language genes and cognition genes at the functional level. Language genes (e.g., *FOXP2*, *CNTNAP2*, etc.) originally manifested as motor control genes during evolution; the proteins they encode participate in the formation of neuromuscular junctions and motor coordination, enabling the vocal organs to produce complex sound sequences. From an evolutionary perspective, the emergence of language ability depends on refined motor skills; therefore, the original functions of language genes are closer to those of motor genes. In contrast, cognition genes are responsible for higher-level integrative functions; they participate in neuronal synaptic plasticity, signal transduction, and cerebral cortex development, enabling organisms to process multimodal stimuli from the environment (sound, light, heat, electricity, etc.) and combine them with memory for judgment and decision-making. The realization of cognitive functions requires an efficient “signal-integration computing hardware,” namely the higher centres of the nervous system, such as the cerebral cortex and subcortical nuclei. Thus, the evolution of cognition genes is closely linked to the increasing complexity of the nervous system, whereas language genes are more associated with motor output circuits [1–4].

Although cognitive functions are typically associated with higher mammals, a large body of research has demonstrated that fish and other lower vertebrates already possess complex cognitive abilities, including spatial memory, social learning, and tool use. For instance, fish can establish cognitive maps, perform quantity discrimination, and engage in avoidance learning. These findings suggest that the neural basis of cognitive functions had already formed early in vertebrate evolution, and that the human cognitive framework may have been gradually modified from that of fish ancestors. Previous studies have proposed that many genes related to human cognition (e.g., those involved in synaptic plasticity) already exist in fish and perform similar functions. Therefore, investigating the evolution of cognition genes from fish to humans can help reveal the core molecular networks or multi-gene SNV polymorphism patterns underlying higher cognitive functions [5–11].

Language function and cognitive function are not independent of each other but overlap at multiple levels. The higher-order aspects of language—such as grammatical comprehension, pragmatic inference, and rhetorical usage—essentially involve cognitive processes including working memory, attention, reasoning, and decision-making. These advanced language functions rely on the engagement of cognitive regions such as the prefrontal cortex; therefore, language genes automatically become part of the cognitive gene network at the higher level. For example, FOXP2 not only regulates vocal motor control but also influences cerebral cortex development and synaptic plasticity, thereby participating in cognitive functions. Similarly, cognition genes such as RELN and TBR1 play critical roles in early neural development, yet their mutations also lead to language disorders. Thus, during evolution, the boundary between language and cognition genes gradually blurs, forming an intertwined regulatory network that jointly shapes the unique communication and thinking abilities of humans [12–15].

The relationship between language and cognition is a foundational issue in contemporary cognitive science and artificial intelligence research. Chomsky’s internalist view of language provides the most influential theoretical framework for this issue: the core function of language is not communication, but thought [16]. The faculty of language generates an internal, individual, intensional computational system (I-language), whose typical use is nearly 100% internal, with externalized communication being merely an “ancillary process” of this internal system. This view is supported by neuroscientific evidence: pseudo-language systems that obey principles of universal grammar can normally activate language brain regions, whereas systems that violate structural dependency and use linear order are treated by the brain as “puzzles” rather than “language.” Anderson, from the perspective of comparative linguistics, argues that although animal communication systems are complex, they lack the symbolic representation and recursive combinatorial properties of human language [17]; “animals can communicate, but they cannot speak.” Fitch further points out that opposing cultural explanations to genetic explanations is conceptually invalid, and that the dual nature of language as both an evolved biological capacity and a cultural entity requires interdisciplinary integration [18].

However, the above theoretical framework faces an empirical testing challenge: if language (as an internal symbolic system) and cognition are indeed functionally integrated, do they also exhibit distinguishable patterns at the molecular evolutionary level? Mozzi et al.’s study showed that the positive selection signal of FOXP2 in the human lineage was already present in archaic hominins (Neanderthals and Denisovans), whereas non-coding changes in genes such as KIAA0319 and CNTNAP2 reached high frequency only after divergence from modern humans [19]. Tiwary further reported [20] that cognitive and language genes were jointly shaped by positive selection and relaxed purifying selection in the human lineage, exhibiting a co-evolutionary trend. However, these studies mostly focused on a small number of candidate genes and lacked systematic quantification of the differences in evolutionary patterns between “language genes” and “cognition genes” at the cross-species, multi-locus level. Based on 13,415 SNV loci from 33 language/cognition-related genes, the present study compares the contribution patterns of the two gene categories at key nodes of the inter-species similarity curve, providing observational evidence at the molecular evolutionary level for the above theoretical controversy.

For the above consideration and background, a key scientific question emerges: along the similarity curve of polymorphism loci in language-cognition genes, do the occurrence frequencies of language genes and cognition genes differ significantly across all steep-slope positions corresponding to polymorphic pattern evolution? And if so, does this difference exhibit directionality and stage-specificity? To address this question, it is necessary to establish a systematic method that can simultaneously quantify multiple species, multiple SNV sites, and the enrichment levels of the two gene categories. The basic rationale of this study is as follows: first, using binary coding of allele presence/absence, all 13145 SNV sites are converted into binary feature vectors, and the similarity of each sample to the reference genome (pp6) is determined based on five classical distance measures; second, the similarity curve is differentiated to identify all steep-slope (peak) positions, which correspond to evolutionary nodes (similarity transitions/candidate clustering boundaries) where multi-SNV patterns undergo rapid changes; third, for each peak, the differential loci between the peak apex sample and its immediate left neighbor are compared, and through a group-intersection strategy (see PeakRank7 in the text), a set of shared differential loci, F_INTERSECTION, is extracted across methods; finally, the proportions of language-gene and cognition-gene loci within F_INTERSECTION are calculated, and the emergence patterns of new loci between adjacent peaks are traced. Through this pipeline, we can not only quantify the enrichment of the two gene categories at breakpoints but also reveal their dynamic changes over time (along the similarity axis), thereby providing new molecular evidence for the co-evolution of language and cognitive abilities.

## 2 Methods

### 2.1 Data sources and preprocessing

All computations were performed using R code. The input data comprised meta-information for 413 cross-taxon samples (meta_data.xlsx) and their 13,415 SNV genotypes (snv_data.xlsx) from 33 language-cognition genes (Supplementary tables, Table 1). The meta_data.xlsx contained seven columns: sample (sample ID), taxon, country, region, age(BP) (years before present), sample_details, and accession (public database source and accession number for whole-genome sequences). The samples covered 11 taxa, including: Amphibian (6 samples), Artiodactyla (40), Birds (50), Cetacea (34), Fish (92), Human (34; 29 ancient hominins + 5 modern humans), Laurasiatherians (38), Others (23), Primates (33), Reptiles (34), and Rodents (29). Among them were representative ancient hominin samples (with complete chronological information) and living-fossil samples (e.g., coelacanth, lungfish, tuatara, etc.), spanning from fish to humans, thus providing rich material for cross-species evolutionary studies of cognition genes. Except for living-fossil species (e.g., lungfish, coelacanth, hagfish, horseshoe crab, *Nautilus pompilius*, pig-nosed turtle, platypus, etc.) and human samples, all other samples were representative species, with no cases of multiple samples from the same species (which would cause redundancy in similarity). The SNV loci were obtained as follows: using a custom-built search software written in Python/C++ [10], we searched the whole-genome sequence files (fasta or fastq formats) of each sample for the 13,415 SNV loci sequences of the 33 human language-cognition genes (SNV base plus equal-length flanking bases on both sides, total length 27 bp). The flanking 26-bp sequences were required to be 100% matched, thus yielding the allelic base at each SNV locus in the sample’s whole-genome sequencing data. The base at each SNV locus was one of the following 16 types: A, T, G, C, AT, AG, AC, TG, TC, GC, ATG, ATC, AGC, TGC, ATGC, or 0 [11].

**Table 1.** Language and Cognition Genes[21–34].

| gene_name | Language | Cognition | Part of known functions |
| --- | --- | --- | --- |
| ARHGAP11B | FALSE | TRUE | Hominin-specific development and evolutionary expansion of the brain neocortex |
| ASPM | TRUE | TRUE | Associated with microcephaly |
| ATP2C2 | TRUE | FALSE | Memory |
| CHRM2 | TRUE | TRUE | A nervous system gene associated with depression disorder |
| CMIP | TRUE | FALSE | Reading, memory |
| CNTNAP2 | TRUE | TRUE | Early language development |
| DAB1 | FALSE | TRUE | Regulation of neuron differentiation |
| DCC | FALSE | TRUE | Associated with impaired intellectual development |
| DCDC2 | TRUE | FALSE | Reading, dyslexia |
| EP300 | FALSE | TRUE | Associated with rare neurological diseases and impairment of intellectual development |
| FADS2 | FALSE | TRUE | Fatty acid desaturase 2 expressed at high levels in the brain |
| FLNC | TRUE | FALSE | Reading, language |
| FOXP1 | TRUE | FALSE | Expressive language |
| FOXP2 | TRUE | TRUE | Speech |
| GNB5 | FALSE | TRUE | Associated with language delay and cognitive impairment |
| GRID2 | FALSE | TRUE | Predominant excitatory neurotransmitter receptors in the mammalian brain |
| HAR1A | FALSE | TRUE | A gene whose expression levels associated with memory and cognitive abilities |
| IGF2R | FALSE | TRUE | Insulin-like growth factor gene associated with behavior/neurological phenotype |
| KIAA0319 | TRUE | FALSE | Reading, dyslexia |
| KMT2D | FALSE | TRUE | Lysine Methyltransferase 2D, associated with intellectual disability and eye diseases |
| MCPH1 | TRUE | TRUE | Associated with microcephaly, primary, autosomal recessive and lymphatic malformation |
| NBPF8 | FALSE | TRUE | A gene associated with macrocephaly, autism, schizophrenia, cognitive disability |
| NFXL1 | TRUE | FALSE | Speech |
| NOTCH2NLA | TRUE | TRUE | Human-specific NOTCH2NL involved in cerebral cortex development |
| NRXN1 | FALSE | TRUE | Neurexin 1 required for efficient neurotransmission and formation of synaptic contacts |
| ROBO1 | TRUE | FALSE | Phonological buffer |
| ROBO2 | TRUE | FALSE | Expressive vocabulary |
| SNAP25 | TRUE | TRUE | Synaptosomal-associated protein 25 |
| SRGAP2 | TRUE | TRUE | Vocal learning, vital for cortical neuron development |
| SRGAP2C | TRUE | FALSE | Human-specific synaptic regulator |
| THSD7B | FALSE | TRUE | Associated with eye diseases/neuronal diseases |
| TM4SF20 | TRUE | FALSE | Language delay, communication disorder |
| TPK1 | TRUE | FALSE | Syntactic and lexical ability |

### 2.2. Data Preprocessing

Each SNP site genotype (e.g., ‘AT’, ‘G’) is decomposed into four binary variables indicating the presence (1) or absence (0) of each of the four nucleotides A, T, C, G. This yields a binary feature vector of length 4 * (number of SNPs) for each sample. Missing data (genotype ‘0’) are treated as NA and excluded from pairwise comparisons. The reference sample is pp6 (modern human standard genome Homo_sapiens.GRCh38).

### 2.3 Distance Measures

For two individuals x and y, let:

- a = number of features where both x and y have 1
- b = number of features where x=1, y=0
- c = number of features where x=0, y=1
- d = number of features where both are 0
- n = a+b+c+d (valid features, excluding NAs)

The following distances are computed:

A. Sørensen (Dice) distance: D_S = 1 - (2a) / (2a + b + c) This is the complement of the Dice coefficient, suitable for binary presence/absence data. Range: [0,1]. 0 indicates identical, 1 indicates no shared features.
B. Rogers distance: D_R = sqrt((b + c) / (2n)) This is the square root of the simple matching dissimilarity normalized by 2n. It is equivalent to the Euclidean distance divided by sqrt(2n). Range: [0,1].
C. Nei distance: D_N = 1 - a / sqrt((a+b)*(a+c)) This is derived from Nei’s genetic distance, using the geometric mean of the total occurrences in each individual. It measures the dissimilarity in allele sharing proportions. Range: [0,1]. 0 if identical, 1 if no shared alleles.
D. Reynolds distance: D_Re = (b + c) / n This is the proportion of differing features, a simple and robust measure for binary data. It is equivalent to the Hamming distance divided by n. Range: [0,1].
E. Hellinger distance: D_H = sqrt(1 - a / sqrt((a+b)*(a+c))) This is the square root of the complement of the Bhattacharyya coefficient for binary vectors. It is a symmetric measure that satisfies the triangle inequality and is widely used in population genetics. Range: [0,1].

### 2.4 Similarity Transformation

For each distance D, we compute similarity as: S = 1 / (1 + D)

This monotonic transformation maps distances to the (0,1] range, with 1 being identical and values approaching 0 as distance increases.

### 2.5 Sample Sorting and Peak Detection

For each method, samples are sorted in ascending order of similarity to the reference (pp6). The first derivative of the similarity curve is approximated by the difference between consecutive points. Only derivatives at positions ≥ 300 are considered to focus on the latter portion of the curve (more derived samples). The top 12 peaks (by absolute derivative value) are retained as candidate evolutionary breakpoints.

### 2.6 Peak Window Analysis

For each peak at index i (start position), a window of samples from i-WINDOW_HALF_SIZE + 1 to i + WINDOW_HALF_SIZE + 1 is extracted. The peak sample is the one at i+1. For each SNP site, we compare the alleles of the peak sample against all other samples in the window. A site is considered ‘different’ if the peak sample has any allele not shared by the other sample (i.e., no intersection of allele sets). The top 600 sites with the highest difference scores are reported.

### 2.7 Left-Window Comparison (SNVgenebyLeft, PeakRank)

For each peak, we also compare the peak sample against a fixed number of samples immediately to its left (positions i-LEFT_WINDOW_SIZE to i-1). Sites where the peak sample has alleles not present in any left sample are recorded. This analysis is used to identify mutations that are unique to the peak sample relative to its immediate predecessors.

### 2.8 Peak Grouping

Peaks from all methods that share the same peak sample are grouped together to identify samples that consistently appear as breakpoints across methods.

### 2.9 PeakRank7: Pairwise Difference Intersection Analysis

To systematically explore the genomic changes at the most prominent breakpoints, we defined 13 groups of peaks based on their start indices (ranging from 355 to 401). Each group contains 3–5 peaks from different methods that occur at the same or very close start positions. For each peak within a group, we compare the peak sample (at position start_idx+1) with its immediate left neighbor (at position start_idx) across all 13,415 SNV sites. Sites where the two samples differ (i.e., their allele sets are not identical) are recorded as a difference set for that peak. Thus, for a group with k peaks, we obtain k difference sets (F1, F2, …, Fk). We then take the intersection of these k sets to obtain F_INTERSECTION (the set of sites that are consistently different between each peak and its left neighbor across all peaks in the group). For each group, we compute the proportion of sites in F_INTERSECTION that belong to language genes and cognition genes (based on our gene annotation). Additionally, we list the top 6 SNV sites in F_INTERSECTION (ranked by the number of peaks in which they appear, but since all appear in all peaks, we simply list them in alphabetical order). These proportions are then plotted as a function of start index to reveal trends in the enrichment of language/cognition genes across the evolutionary transition from position 355 to 401.

Additionally, to track the sequential emergence of new SNV sites along the similarity axis (from 355 to 401), we compute pairwise differences between consecutive F_INTERSECTION sets: i.e., F_INTERSECTION_2 F_INTERSECTION_1, F_INTERSECTION_3 F_INTERSECTION_2, …, F_INTERSECTION_13 F_INTERSECTION_12. These difference sets represent sites that appear uniquely in the later group compared to the immediately preceding group, thus highlighting newly gained genomic changes at each step.

## 3 Results and Analysis

### 3.1 Consistency assessment of similarity among methods

To evaluate the consistency of the five genetic distance methods in capturing interspecific genetic similarity patterns, we calculated the Pearson correlation coefficients between the similarity vectors derived from each method. The Pearson correlation coefficient measures the strength and direction of linear correlation between two variables, ranging from –1 to 1; the closer the absolute value is to 1, the stronger the consistency. This analysis was based on the similarity scores (sim_original matrix) for all samples under each method, encompassing 413 samples. The resulting correlation coefficient matrix is shown in Table 2. The average correlation coefficient was 0.982, with the lowest being 0.956. All inter-method correlations were highly positive (>0.9), indicating that although the distance metrics differ in definition, the relative genetic similarity patterns among species captured by them are highly consistent and the results are robust.

**Table 2.**
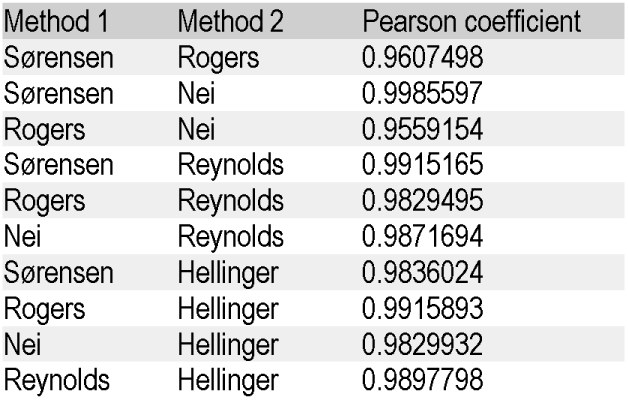
Pearson correlation coefficients of similarity vectors among the five methods.

| Method 1 | Method 2 | Pearson coefficient |
| --- | --- | --- |
| Sørensen | Rogers | 0.9607498 |
| Sørensen | Nei | 0.9985597 |
| Rogers | Nei | 0.9559154 |
| Sørensen | Reynolds | 0.9915165 |
| Rogers | Reynolds | 0.9829495 |
| Nei | Reynolds | 0.9871694 |
| Sørensen | Hellinger | 0.9836024 |
| Rogers | Hellinger | 0.9915893 |
| Nei | Hellinger | 0.9829932 |
| Reynolds | Hellinger | 0.9897798 |

### 3.2 Comparative analysis of similarity and derivative curves

In Figure 1, multiple steep rising segments (i.e., steep slopes) are visible, corresponding to regions of rapid similarity jumps—these are exactly the peak positions we detected. For instance, at positions around 355, 363, and 381 (Figure 2), the curve rises almost vertically, suggesting that the genomes of samples at these positions have undergone a large emergence of new alleles. The essential point of this figure is that the presence of steep slopes indicates similarity transitions/candidate clustering boundaries, which in turn imply a sudden acceleration in evolutionary rate, whereas plateaus represent relatively stable genetic phases. By identifying steep-slope positions, we can locate evolutionary breakpoints of similarity transitions/candidate clustering boundaries and further analyse the enrichment of specific SNVs of language and cognition genes at these breakpoints. It should be noted that changes in sample composition after similarity ranking cannot automatically be equated with “evolutionary rate.” A distinction should be made between “large ranking gradient” and “fast evolutionary rate over time.” Although steep slopes may partly arise from sampling bias (e.g., lack of genomic data for representative intermediate species), we consider them still of substantial quantitative value. First, the slope of a steep segment reflects the change in similarity per unit ranking step; this value is not affected by absolute species richness, so it can still serve as a relative indicator of evolutionary rate. Second, even if sampling is insufficient, the appearance of steep slopes suggests that significant genetic clustering already exists among the available samples, and these clusters often coincide with known evolutionary events (e.g., speciation or migration). The small plateaus between steep slopes represent intervals of relatively stable similarity, possibly corresponding to prolonged periods of niche conservatism or genetic drift equilibrium. Notably, these small plateaus have been confirmed not to result from redundant sampling of the same species.

**Figure 1.**
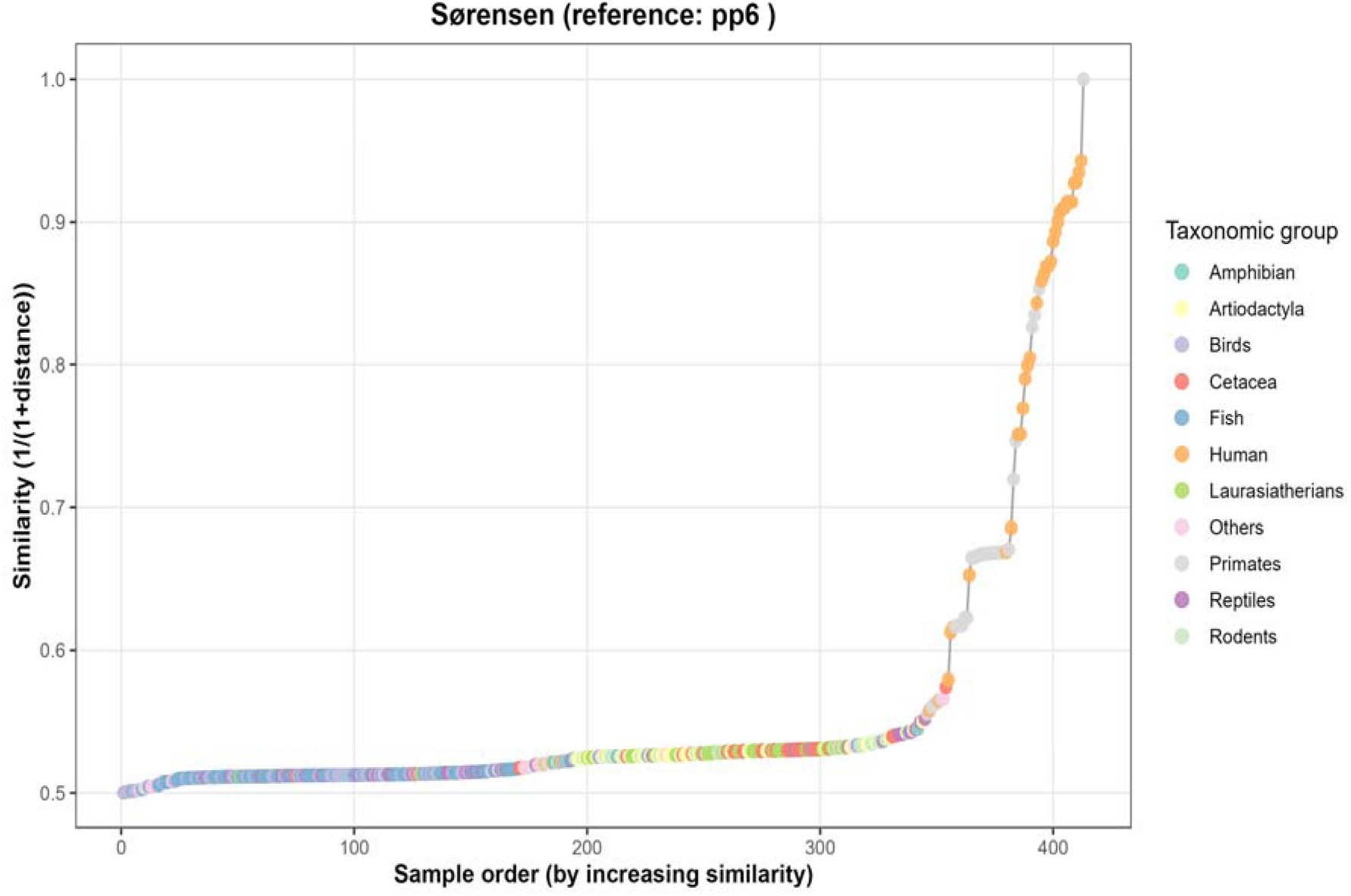
Representative similarity curve (Sørensen method). The horizontal axis represents the sample rank order (dimensionless integer), and the vertical axis represents the similarity value (range 0–1, dimensionless), defined as *S* = 1/(1 +*D*), where *D* is the genetic distance. Each point denotes a sample, with colours indicating its taxonomic group. The curve shows an overall upward trend, indicating that as ranking proceeds, the genetic similarity of samples to the reference genome (pp6) gradually increases. The similarity curves for the other methods are provided in the supplementary material (Figure S1).

**Figure 2.**
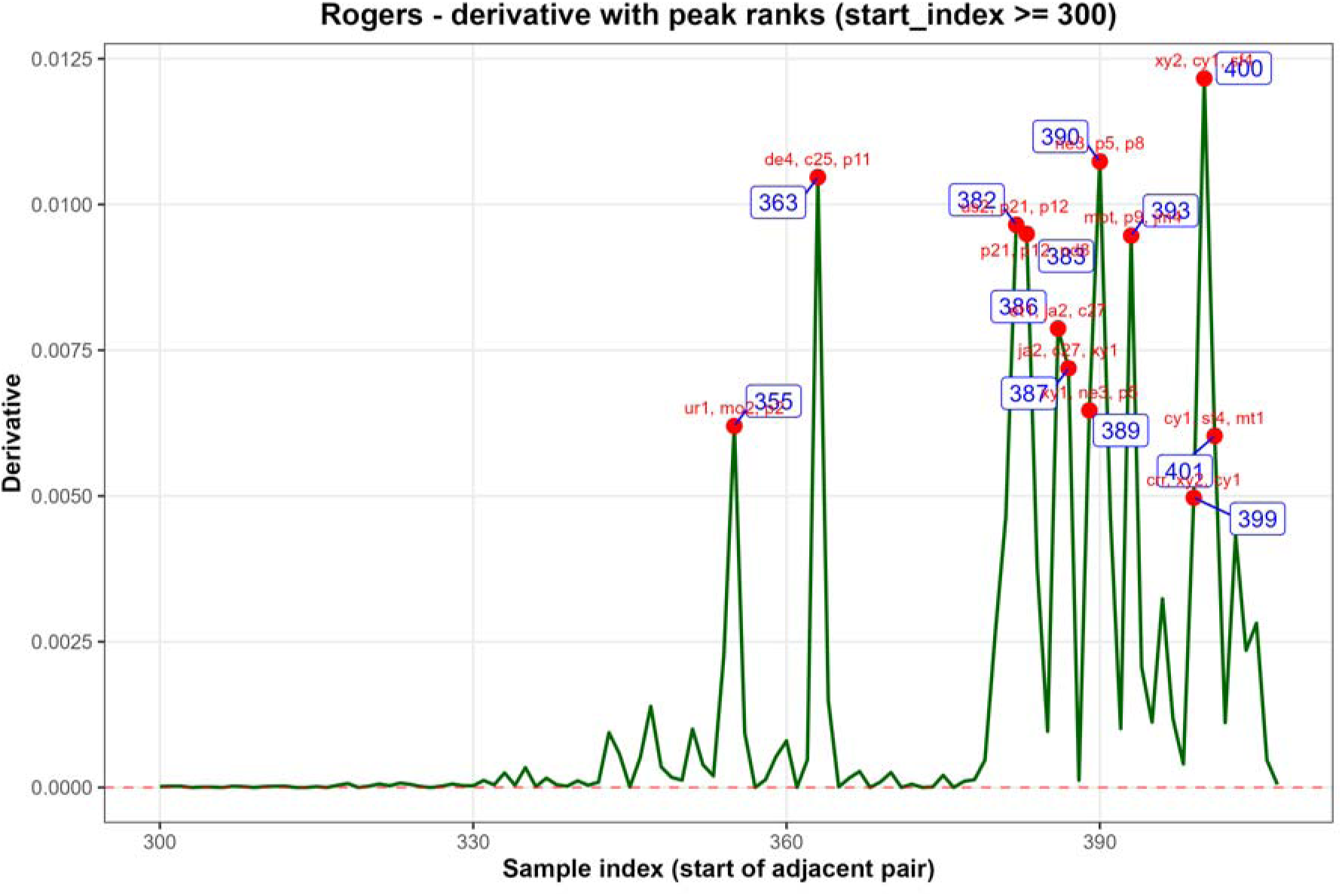
Representative derivative curve (Rogers method, with peak markers). This figure shows the first-derivative curve for the Rogers method, with the horizontal axis representing the start position (dimensionless integer) and the vertical axis representing the derivative value (dimensionless). Red dots mark the detected peaks, blue labels indicate the numerical values of the start positions, and red text annotates the sample names involved at those positions. A set of major peaks (e.g., 355, 363, 381, 400) and several minor peaks are visible in the figure. The key point of this figure is that the higher the peak, the more dramatic the similarity change at that position; the annotated sample names help associate specific evolutionary events (e.g., archaic hominin samples). Derivative curves for the other methods are provided in the supplementary material (Figure S2).

**Figure 3.**
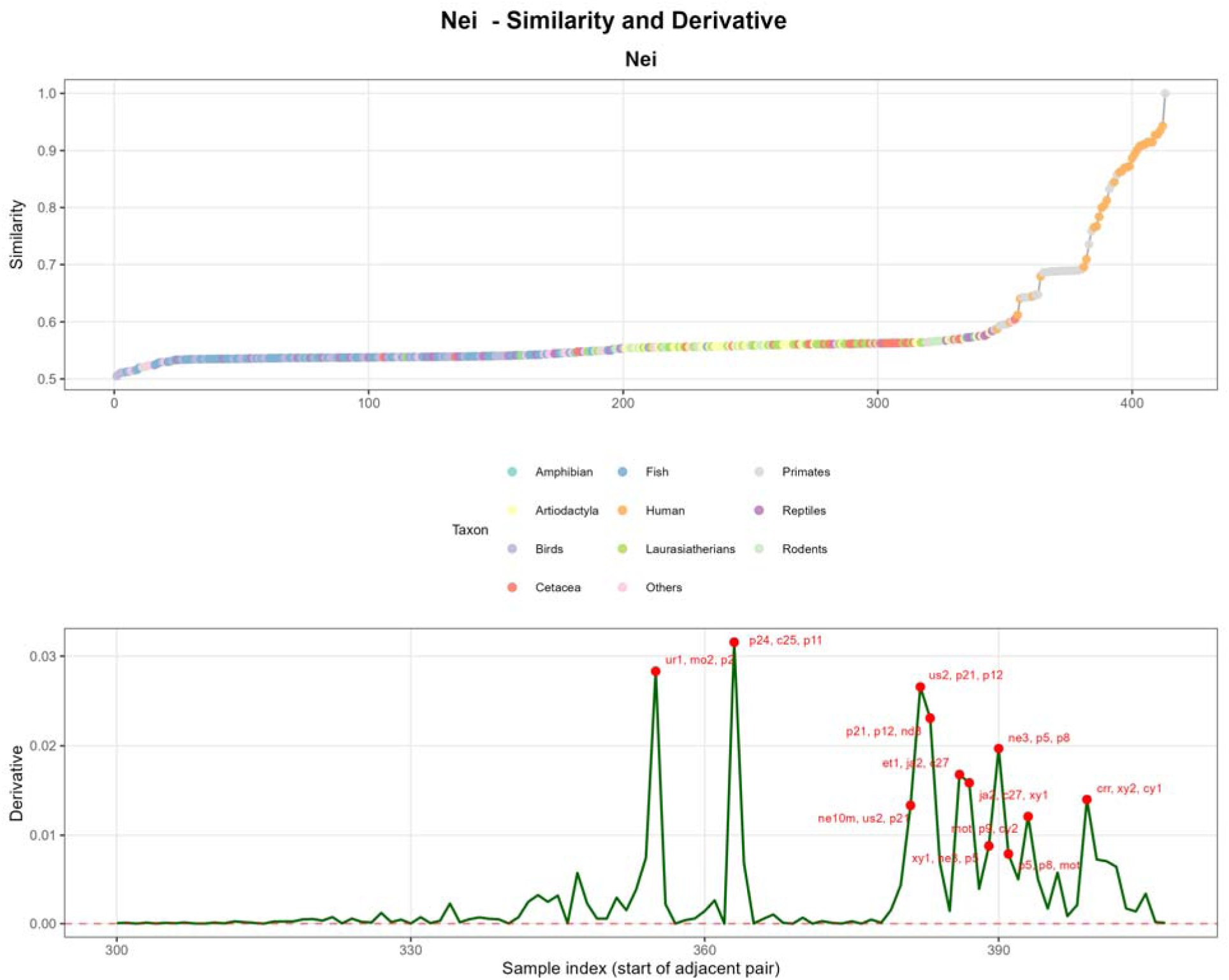
Representative combined plot (Nei method: similarity + derivative). The upper panel shows the similarity curve, and the lower panel shows the first-derivative curve of that curve. In the derivative plot, the horizontal axis represents the start position (i.e., the index of adjacent sample pairs, dimensionless integer), and the vertical axis represents the derivative value (dimensionless, indicating the difference in similarity between adjacent samples). Red dots mark the detected peaks (the top 12 with the largest absolute values). Note that the horizontal axis of the lower panel starts from rank 300 to clearly display the peak positions. By comparing the upper and lower panels, the correspondence between steep-slope positions and derivative peaks can be clearly observed. In this figure, the three most prominent positive peaks (at 355, 363, and 381) correspond to positions where significant genetic transitions (similarity jumps/candidate clustering boundaries) occurred. The value of this plot lies in its intuitive display of the relationship between peaks and the rate of similarity change, facilitating verification of the statistical significance of breakpoints. Combined plots for the other methods are provided in the supplementary material (Figure S3).

The basic content along the horizontal axis of the above similarity curve (Figure 1), from left to right, is as follows: before position 100, the samples are almost entirely fish and birds; from 100 to 200, they include fish, amphibians/reptiles, artiodactyls, cetaceans, laurasiatherians, and a small number of rodents; from 200 to 250, there is a dense representation of artiodactyls, with a few laurasiatherians, fish, and rodents; from 250 to 300, there are many laurasiatherians, along with a few artiodactyls, cetaceans, rodents, and reptiles; from 300 to 350, there are numerous reptiles, a relatively large number of cetaceans and laurasiatherians, and a small number of rodents and artiodactyls; starting at 350, with the exception of a very few cetacean and reptile samples, the composition immediately shifts entirely to primates and archaic hominins, continuing to the very end. The earliest archaic hominin samples are African and American specimens, followed by a large number of Eurasian archaic hominins, but the approximately 10–15 samples on the far right are mainly African archaic hominins plus the last few modern human samples (the final one being the control pp6).

Comparing Figures 4–5 and Figures 7–8, it can be observed that the three positions—355, 363, and 381 (382)—occur with the highest frequency, with multiple methods simultaneously detecting large peaks at these positions. These three peaks represent excellent windows for examining the sudden shift (as seen in Figures 4 and 6) in the polymorphism patterns of language-cognition genes from other animal groups (especially several living-fossil reptiles) towards archaic hominins. The annotated sample names reveal the following: near 355, there are Denisovan de4 (Denisova2, PRJEB20653), African archaic hominin mo2 (Iberomaurusian, PRJNA422662), and American archaic hominin ur1 (Uruguay (CH13), PRJEB48360); near 363, there are c25 (China (WGM70), PRJEB36297), de4, and several primate samples (p11, p24); near 381 (382), samples such as us2 (US ancient Anzick, PRJEB29074), ne10m (Mebrak, Nepal, PRJEB41752), et1 (Ancient Ethiopian Mota, PRJNA295861), nd8 (Neandertal Mezmaiskaya-2, PRJEB21881), and p21 (*Nomascus leucogenys*, Ensembl) appear. This distribution pattern suggests, first, that early hominins had already dispersed across Africa, Eurasia, and the Americas; and second, that these breakpoints (similarity transition/candidate clustering boundary points) may correspond to major evolutionary events such as hominid-hominin divergence and Neanderthal-Denisovan divergence. This figure provides an intuitive positional basis for the subsequent PeakRank7 grouping.

**Figure 4.**
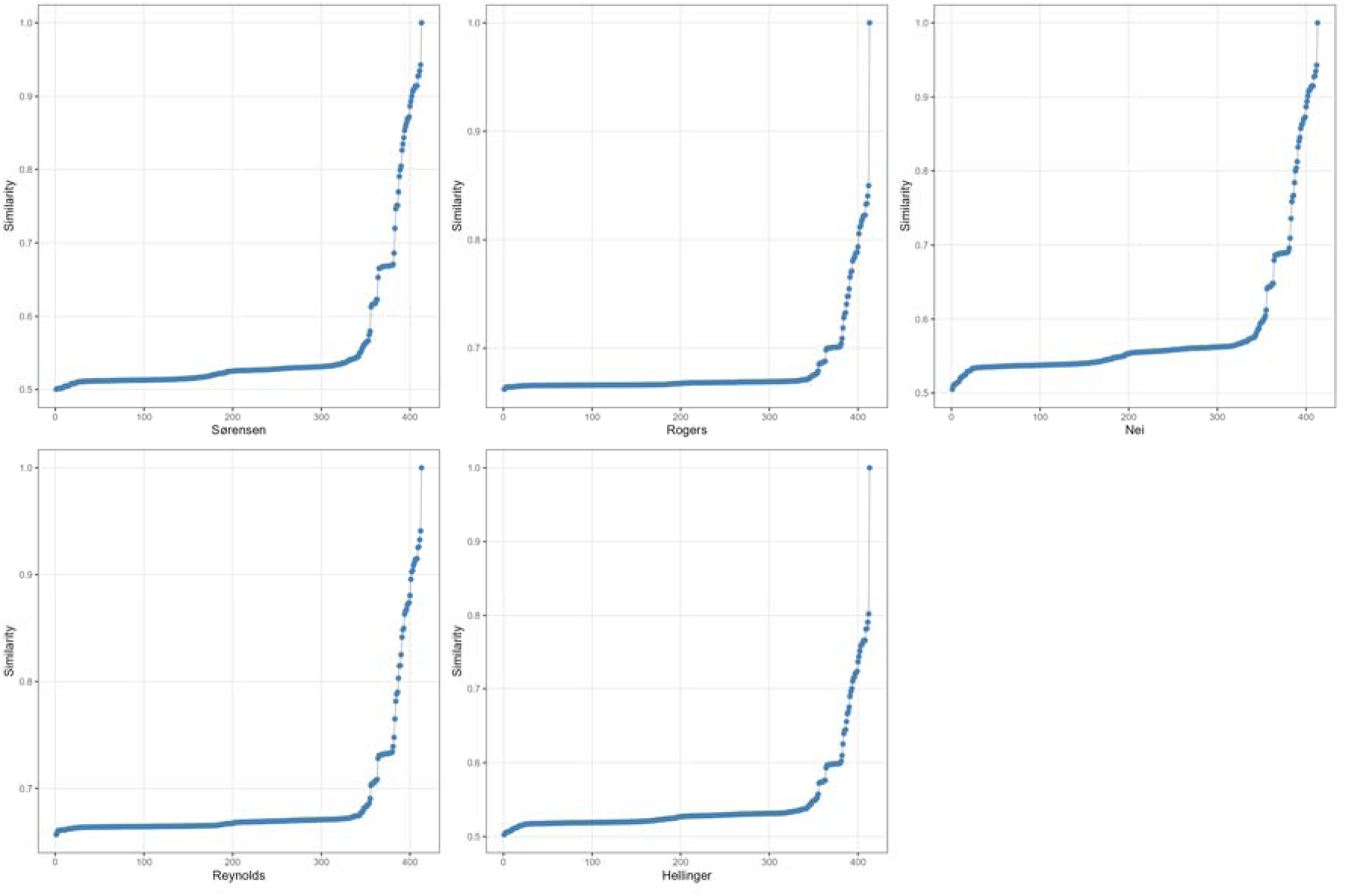
Overlay comparison of similarity curves from the five methods. This figure displays the similarity curves of the five methods side by side, with each subplot corresponding to one method. The horizontal axis represents the sample rank order after sorting (dimensionless integer), and the vertical axis represents similarity (0–1). All five curves show an upward trend in the later portion, but with different slopes: Rogers and Reynolds are steeper, Sørensen and Hellinger are gentler, and Nei is intermediate. Despite differences in ordering, the regions of peak concentration (approximately 350–400) are highly overlapping across methods, indicating that these breakpoints are robust evolutionary hotspots (similarity transition/candidate clustering boundary points).

**Figure 5.**
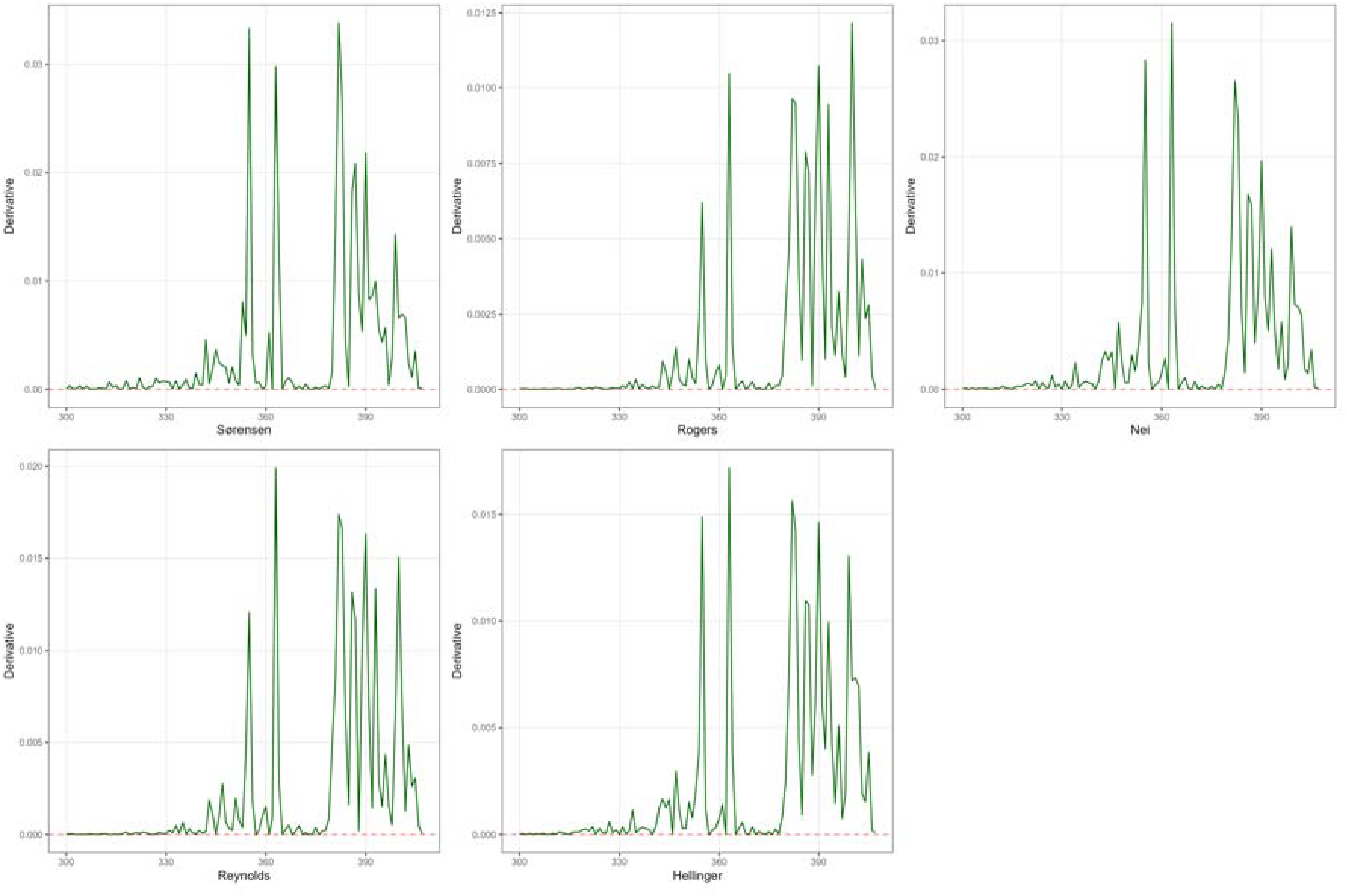
Overlay comparison of derivative curves from the five methods. This figure displays the derivative curves of the five methods, with the horizontal axis representing the start position (dimensionless integer) and the vertical axis representing the derivative value (dimensionless). All methods show distinct positive peaks in the 350–400 interval, though peak heights and exact positions shift slightly. Sørensen and Nei exhibit the highest peaks (>0.03), reflecting their strong sensitivity to genetic differences. Through overlay comparison, it is confirmed that 355, 363, and 381 (382) are the three major peaks commonly shared across the five methods. The significance of this figure lies in helping to filter out the most robust breakpoints and to distinguish universal changes from method-dependent variations.

**Figure 6.**
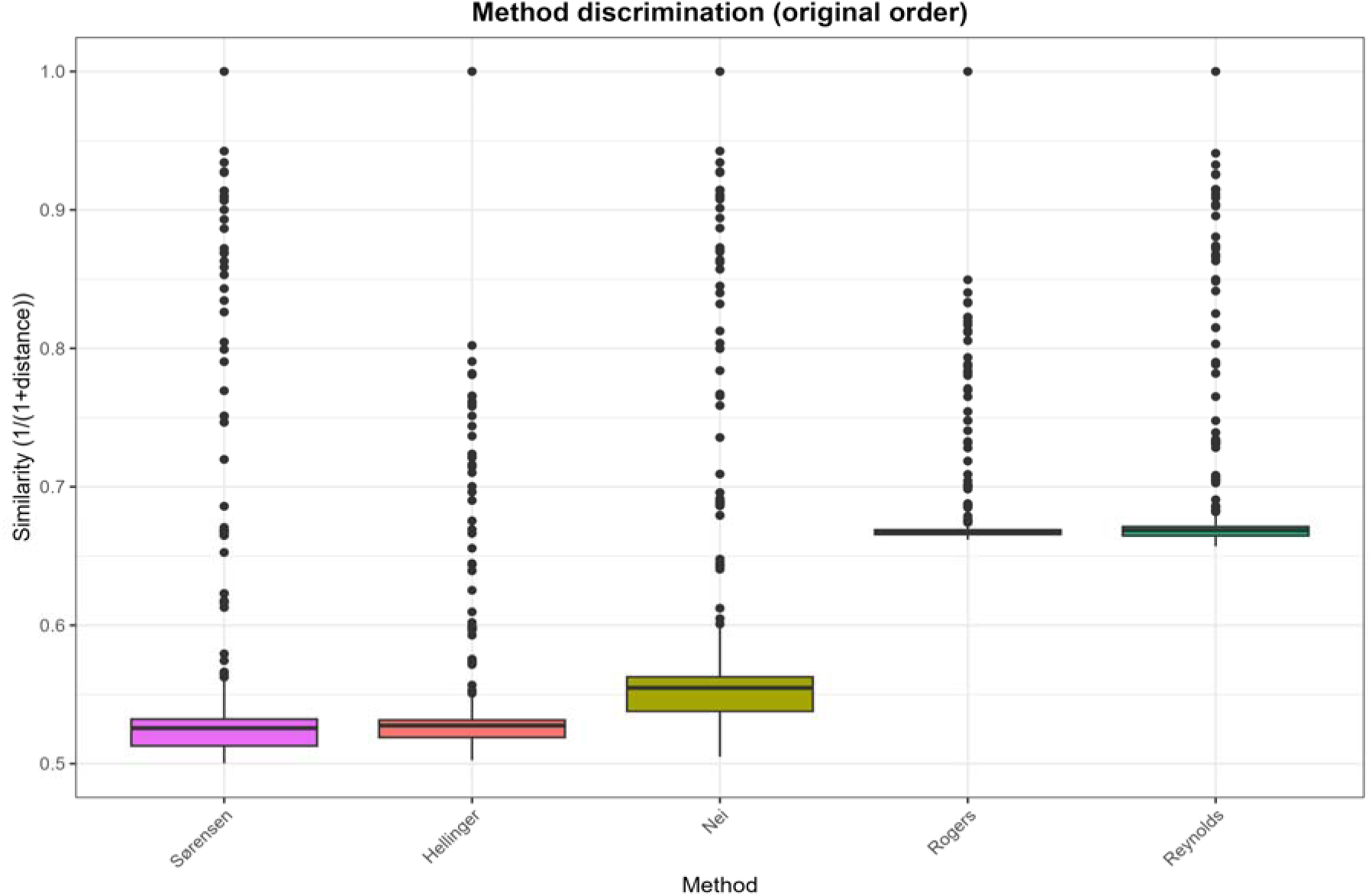
Boxplot of similarity distributions for the five methods. The horizontal axis represents the five methods, and the vertical axis represents similarity values (0–1). Boxes indicate the interquartile range (IQR), with the central line denoting the median, and whiskers extending to 1.5× IQR. Sørensen and Hellinger have the lowest median similarity (∼0.53), Rogers and Reynolds the highest (∼0.67), and Nei falls in between. All distributions are right-skewed (with longer upper whiskers), indicating the presence of a small number of samples (humans and primates) that are highly similar to the reference genome.

**Figure 7.**
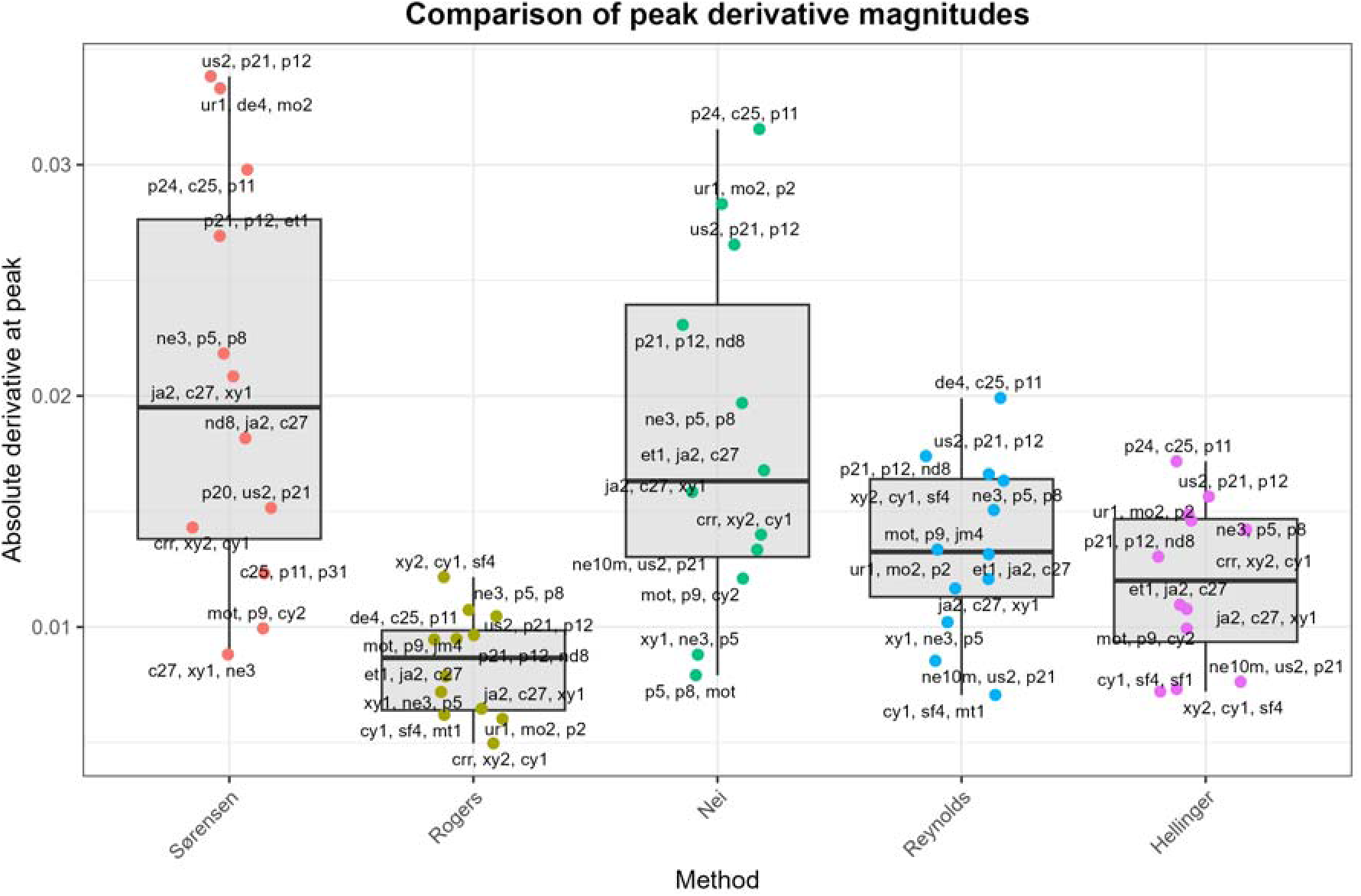
Distribution of absolute values of peak derivatives across methods. The horizontal axis represents the method, and the vertical axis represents the absolute value of the peak derivative (dimensionless). The boxplot shows the central tendency and dispersion of the intensity. Sørensen and Nei exhibit significantly higher peak intensities than the other methods (median ∼0.018 vs. 0.016), indicating their greater sensitivity to dramatic changes. All methods show outliers (beyond the upper whisker), and these extreme values correspond to the top peaks in terms of derivative magnitude, i.e., the breakpoints of primary interest.

**Figure 8.**
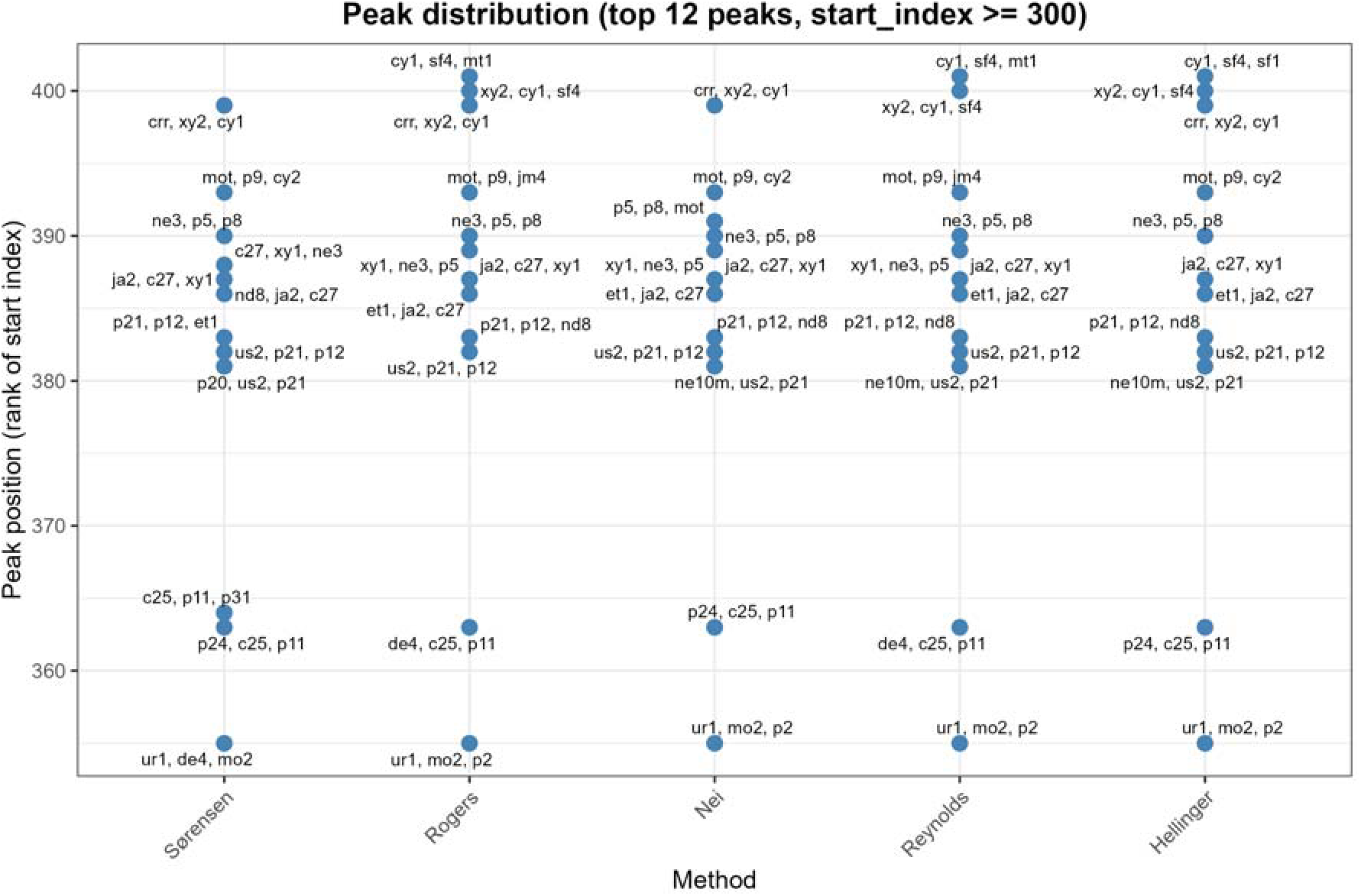
Distribution of peak positions across methods. The horizontal axis represents the method, and the vertical axis represents the start index (dimensionless integer). Each point denotes a peak, with sample names annotated alongside the points. The majority of peaks are concentrated in the 300–420 interval.

### 3.3 Peak analysis

#### 3.3.1 Grouping of peak apex samples

Table 3 groups all peaks whose apex samples are identical, facilitating the identification of recurrent breakpoint samples. By comparing each peak apex sample with its immediate left neighbour, the newly emerged SNV loci are obtained, and subsequently the scores and proportions of the two gene categories at these SNV loci are derived (Table 4).

**Table 3.** Peak sample information.

| Method | Rank | Start_Index | Peak_Sample | Method | Rank | Start_Index | Peak_Sample | Method | Rank | Start_Index | Peak_Sample |
| --- | --- | --- | --- | --- | --- | --- | --- | --- | --- | --- | --- |
| Sørensen | 1 | 382 | p21 | Sørensen | 6 | 387 | c27 | Sørensen | 11 | 393 | p9 |
| Rogers | 4 | 382 | p21 | Rogers | 8 | 387 | c27 | Rogers | 6 | 393 | p9 |
| Nei | 3 | 382 | p21 | Nei | 7 | 387 | c27 | Nei | 10 | 393 | p9 |
| Reynolds | 2 | 382 | p21 | Reynolds | 9 | 387 | c27 | Reynolds | 6 | 393 | p9 |
| Hellinger | 2 | 382 | p21 | Hellinger | 8 | 387 | c27 | Hellinger | 9 | 393 | p9 |
| Sørensen | 3 | 363 | c25 | Sørensen | 7 | 386 | ja2 | Rogers | 1 | 400 | cy1 |
| Rogers | 3 | 363 | c25 | Rogers | 7 | 386 | ja2 | Reynolds | 5 | 400 | cy1 |
| Nei | 1 | 363 | c25 | Nei | 6 | 386 | ja2 | Hellinger | 12 | 400 | cy1 |
| Reynolds | 1 | 363 | c25 | Reynolds | 7 | 386 | ja2 | Rogers | 9 | 389 | ne3 |
| Hellinger | 1 | 363 | c25 | Hellinger | 7 | 386 | ja2 | Nei | 11 | 389 | ne3 |
| Sørensen | 4 | 383 | p12 | Sørensen | 8 | 381 | us2 | Reynolds | 10 | 389 | ne3 |
| Rogers | 5 | 383 | p12 | Nei | 9 | 381 | us2 | Rogers | 10 | 355 | mo2 |
| Nei | 4 | 383 | p12 | Reynolds | 11 | 381 | us2 | Nei | 2 | 355 | mo2 |
| Reynolds | 3 | 383 | p12 | Hellinger | 10 | 381 | us2 | Reynolds | 8 | 355 | mo2 |
| Hellinger | 5 | 383 | p12 | Sørensen | 9 | 399 | xy2 | Hellinger | 3 | 355 | mo2 |
| Sørensen | 5 | 390 | p5 | Rogers | 12 | 399 | xy2 | Rogers | 11 | 401 | sf4 |
| Rogers | 2 | 390 | p5 | Nei | 8 | 399 | xy2 | Reynolds | 12 | 401 | sf4 |
| Nei | 5 | 390 | p5 | Hellinger | 6 | 399 | xy2 | Hellinger | 11 | 401 | sf4 |
| Reynolds | 4 | 390 | p5 | Sørensen | 10 | 364 | p11 | Sørensen | 2 | 355 | de4 |
| Hellinger | 4 | 390 | p5 | Sørensen | 12 | 388 | xy1 | Nei | 12 | 391 | p8 |

**Table 4.** Proportions of the two gene categories at peaks.

| Category | Gene number | Total score | Proportion |
| --- | --- | --- | --- |
| Language genes | 19 | 256,974 | 56.8% |
| Cognition genes | 18 | 244,847 | 54.1% |
| Total genes | 33 | 452,233 | 100% |

#### 3.3.2 Frequency analysis of peak apex samples (PeakRank)

The PeakRank analysis identified individuals that frequently appeared as apex samples across all methods (occurrence count ≥3). These samples represent evolutionary breakpoints (similarity transition/candidate clustering boundary points) that are consistently detected across methods. Table 5 lists these samples and their occurrence frequencies across the five similarity curves. Among them, the archaic hominin samples with the highest frequencies are c25, c27, and ja2 (see meta_data for details), while four primate samples also reached the highest frequency: p12 (*Pongo abelii*), p21 (*Nomascus leucogenys*), p5 (*Gorilla gorilla*), and p9 (*Pan troglodytes*). Except for gibbon p21, the other three are the great apes with the highest levels of intellectual evolution. The p11 (*Piliocolobus tephrosceles*, Ugandan red colobus, an Old World monkey) and p8 (*Pan paniscus*, bonobo) also appeared as peaks (Table 3), but with lower frequencies compared to the aforementioned three apes.

**Table 5.** Frequently occurring peak apex samples.

| Peak_Sample | Frequency | Peak_Sample | Frequency | Peak_Sample | Frequency |
| --- | --- | --- | --- | --- | --- |
| c25 | 5 | ne3 | 3 | sf4 | 3 |
| c27 | 5 | p12 | 5 | us2 | 4 |
| cy1 | 3 | p21 | 5 | xy2 | 4 |
| ja2 | 5 | p5 | 5 |  |  |
| mo2 | 4 | p9 | 5 |  |  |

#### 3.3.3 Major peak apex samples for each method

Table 6 lists the apex samples corresponding to the top 3 peaks (ranked by absolute derivative value) for each method; these samples represent the most significant evolutionary breakpoints detected by each method.

**Table 6.** Top 3 peaks (ranked by absolute derivative value) for five methods.

| Method | Rank | Start_Index | Peak_Sample | Derivative | Method | Rank | Start_Index | Peak_Sample | Derivative |
| --- | --- | --- | --- | --- | --- | --- | --- | --- | --- |
| Sørensen | 1 | 382 | p21 | 0.03382492 | Reynolds | 1 | 363 | c25 | 0.01990855 |
| Sørensen | 2 | 355 | de4 | 0.03330728 | Reynolds | 2 | 382 | p21 | 0.01739306 |
| Sørensen | 3 | 363 | c25 | 0.02978674 | Reynolds | 3 | 383 | p12 | 0.01661396 |
| Rogers | 1 | 400 | cy1 | 0.01216098 | Hellinger | 1 | 363 | c25 | 0.01717041 |
| Rogers | 2 | 390 | p5 | 0.01073752 | Hellinger | 2 | 382 | p21 | 0.01564249 |
| Rogers | 3 | 363 | c25 | 0.01046650 | Hellinger | 3 | 355 | mo2 | 0.01487329 |
| Nei | 1 | 363 | c25 | 0.03154363 |  |  |  |  |  |
| Nei | 2 | 355 | mo2 | 0.02830479 |  |  |  |  |  |
| Nei | 3 | 382 | p21 | 0.02654473 |  |  |  |  |  |

It is observed that the two samples with the highest occurrence among the top 3 peaks across all methods are p21 and c25; the former is a modern gibbon sample, and the latter is an East Asian archaic hominin sample. Gibbons diverged from the hominid lineage approximately 16 million years ago [35]; nevertheless, their polymorphism patterns at the 13,415 SNVs of the 33 language-cognition genes remain highly similar to those of many archaic hominin samples, with their peak positions located along the horizontal axis between 380 and 385 in Figure 8.

#### 3.3.4 Occurrence frequency of archaic hominin samples among peaks

Among all peaks, archaic hominin samples (age >0 BP) appeared as apex samples a total of 38 times, involving 11 distinct archaic hominin individuals (Table 7). The high-frequency occurrence of these archaic hominin samples suggests that language-cognition genes underwent strong selection during archaic hominin evolution.

**Table 7.**
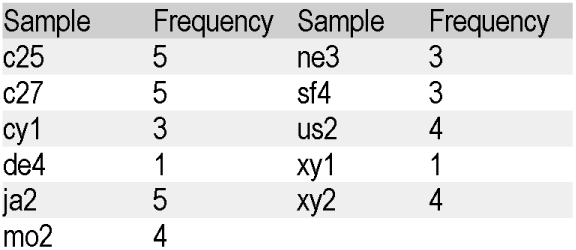
Peak frequency for archaic hominin samples.

### 3.4 PeakRank7: pairwise differential intersection analysis

To systematically dissect the genetic changes at multiple breakpoints within the 355–401 interval, we defined 13 peak groups (group1–group13), each containing 3–5 peaks derived from different methods. For each peak within a group, we compared the SNV differences between its apex sample and its immediate left neighbor (i.e., the preceding sample) to obtain the set of differential loci for that peak (F1, F2, …, Fk). The intersection of these sets was then taken to yield F_INTERSECTION, i.e., the differential loci shared by all peaks in the group. For each F_INTERSECTION, we counted the number and proportion of loci belonging to language genes and cognition genes (Table 8), and listed the top six loci. Note that Table 8 is based on statistics across all 13,415 SNVs, whereas Table 4, although also searched within the 13,415 SNVs, focuses on the top 600 SNV loci with the highest scores.

**Table 8.**
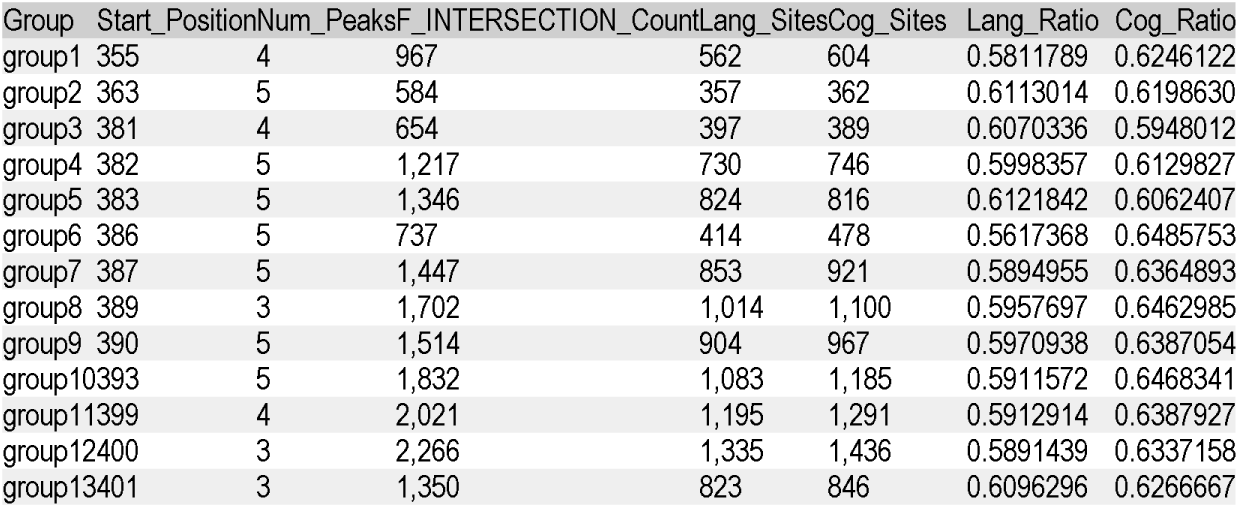
Statistics for F_INTERSECTION (pairwise differential intersection analysis)

| Group | Start_Position | Num_Peaks | F_INTERSECTION_Count | Lang_Sites | Cog_Sites | Lang_Ratio | Cog_Ratio |
| --- | --- | --- | --- | --- | --- | --- | --- |
| group1 | 355 | 4 | 967 | 562 | 604 | 0.5811789 | 0.6246122 |
| group2 | 363 | 5 | 584 | 357 | 362 | 0.6113014 | 0.6198630 |
| group3 | 381 | 4 | 654 | 397 | 389 | 0.6070336 | 0.5948012 |
| group4 | 382 | 5 | 1,217 | 730 | 746 | 0.5998357 | 0.6129827 |
| group5 | 383 | 5 | 1,346 | 824 | 816 | 0.6121842 | 0.6062407 |
| group6 | 386 | 5 | 737 | 414 | 478 | 0.5617368 | 0.6485753 |
| group7 | 387 | 5 | 1,447 | 853 | 921 | 0.5894955 | 0.6364893 |
| group8 | 389 | 3 | 1,702 | 1,014 | 1,100 | 0.5957697 | 0.6462985 |
| group9 | 390 | 5 | 1,514 | 904 | 967 | 0.5970938 | 0.6387054 |
| group10 | 393 | 5 | 1,832 | 1,083 | 1,185 | 0.5911572 | 0.6468341 |
| group11 | 399 | 4 | 2,021 | 1,195 | 1,291 | 0.5912914 | 0.6387927 |
| group12 | 400 | 3 | 2,266 | 1,335 | 1,436 | 0.5891439 | 0.6337158 |
| group13 | 401 | 3 | 1,350 | 823 | 846 | 0.6096296 | 0.6266667 |

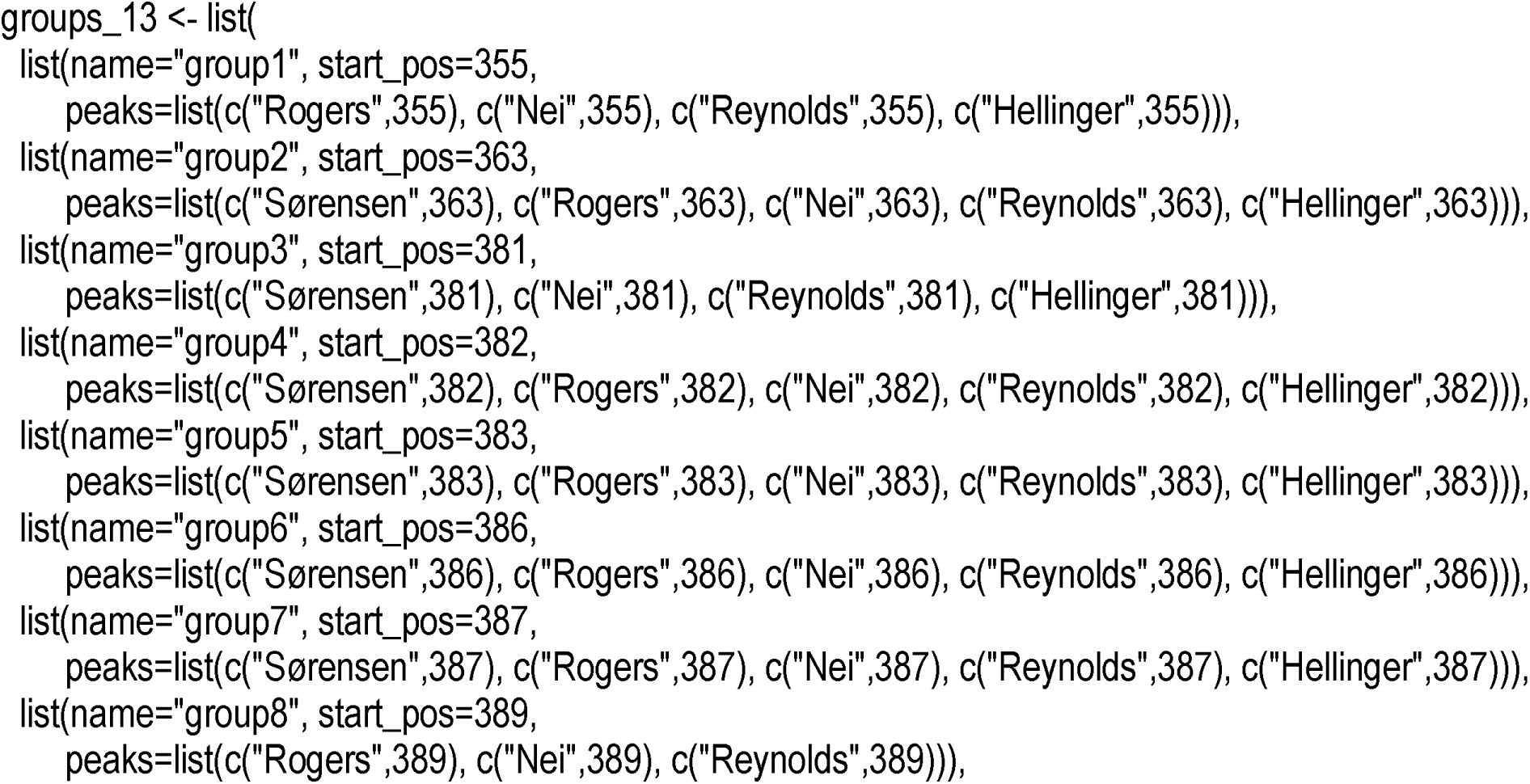
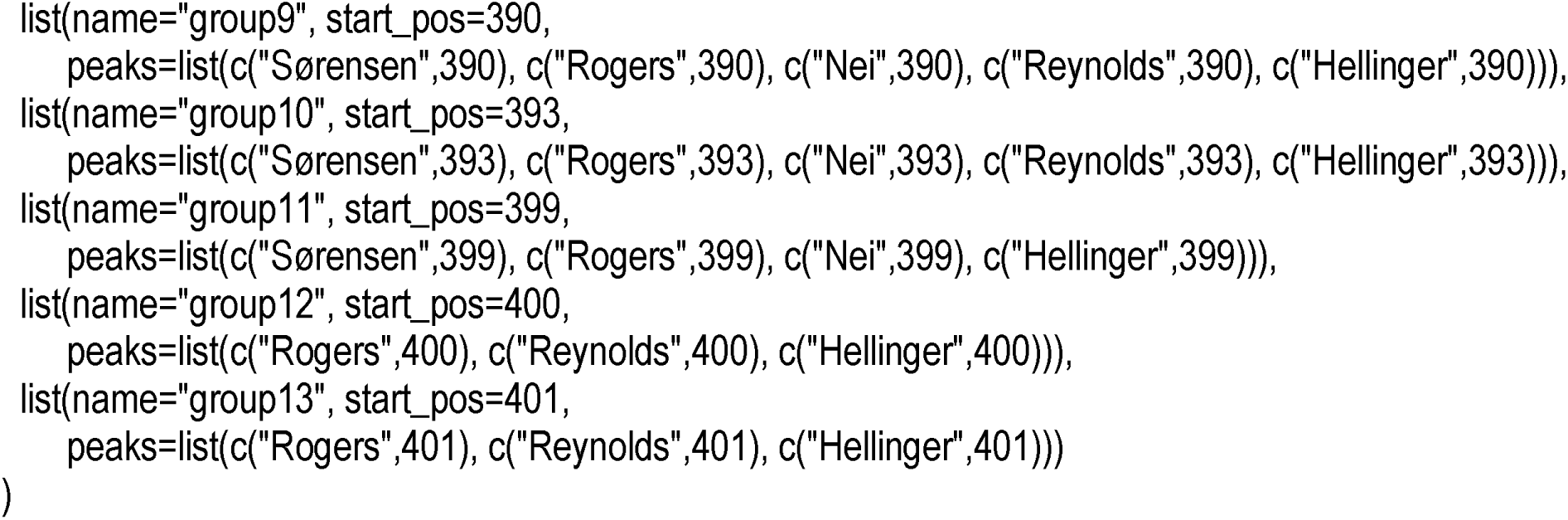

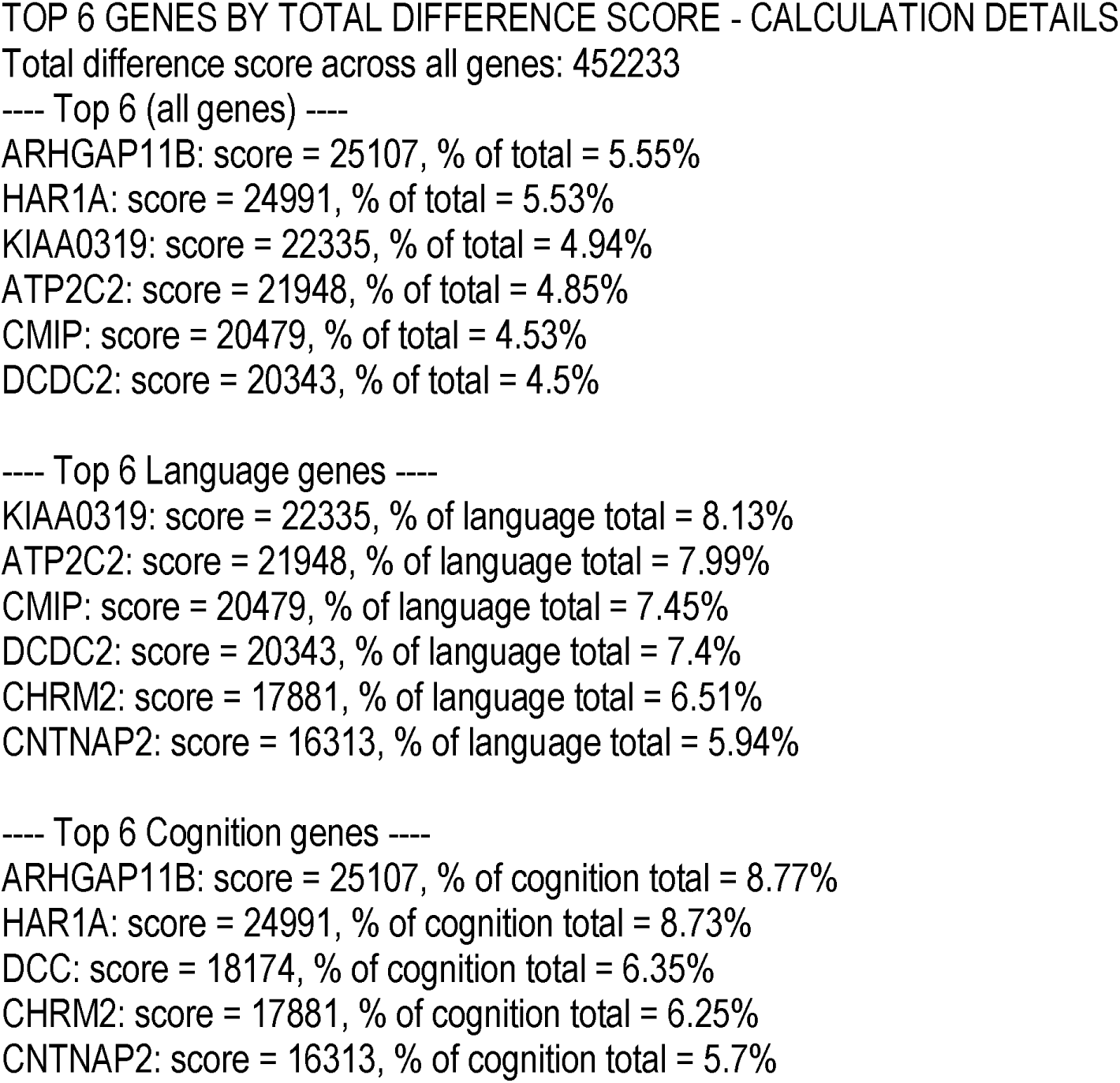
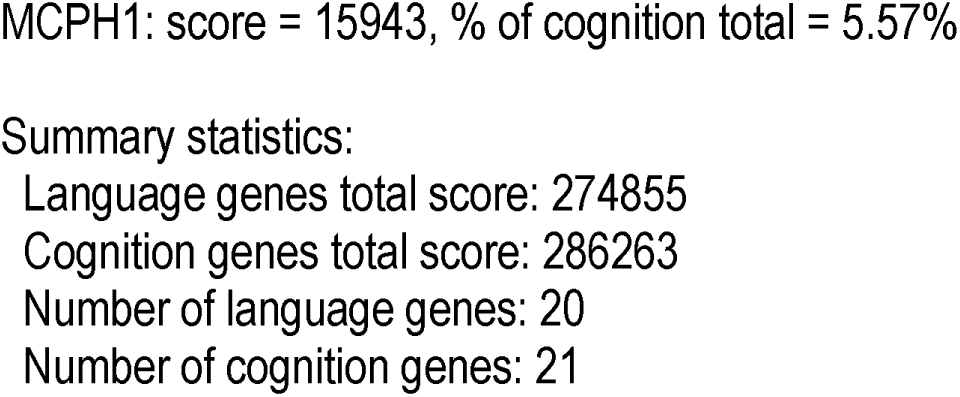

#### 3.4.1 Differential sets between adjacent F_INTERSECTION (newly emerged loci)

Table 9 lists, from group 2 to group 13, the newly added SNV loci for each group relative to the previous group (i.e., loci present in the latter group’s F_INTERSECTION but absent in the former group’s F_INTERSECTION), along with the numbers of associated genes. This reflects the stepwise acquisition of new mutations during evolution. The early, middle, and late stages each appear to have a maximum number of newly added loci, namely 719, 872, and 1,152, respectively (see also Figure 10). Although the sample arrangement along the horizontal axis of the similarity curves cannot theoretically be taken as a true evolutionary timeline for the SNV polymorphism patterns of cognition genes, Figures 1–3 clearly show that the samples on the horizontal axis are essentially ordered in the general direction from fish → amphibians → reptiles → birds → rodents → primates. Therefore, the horizontal axis from left to right can be roughly approximated as an evolutionary timeline, and starting from around coordinate 350, humans suddenly accelerated into distinct early-, middle-, and late-stage evolutionary phases.

**Figure 9.**
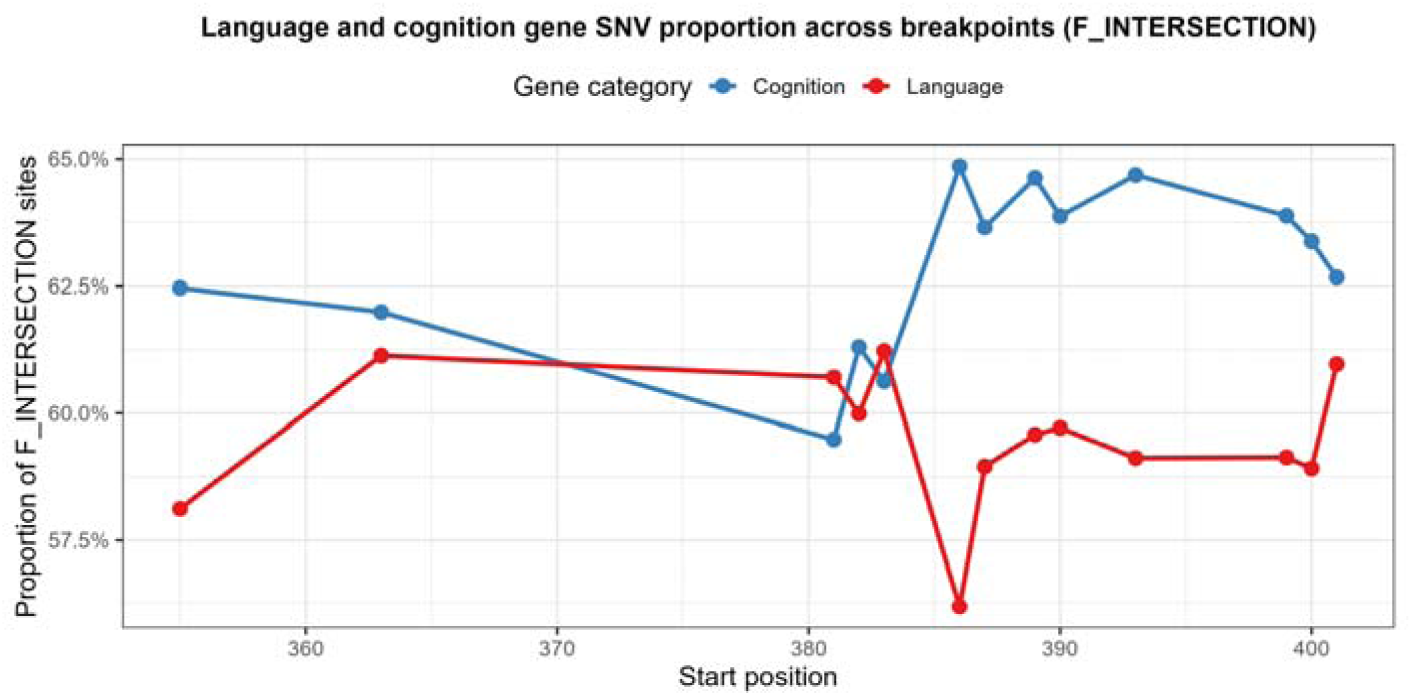
Proportions of language genes and cognition genes in F_INTERSECTION as a function of start position. The horizontal axis represents the start position of each peak group (dimensionless integer, 355–401), and the vertical axis represents the proportion of F_INTERSECTION loci that belong to language genes or cognition genes (ratio, 0–1). The red curve denotes language genes, and the blue curve denotes cognition genes. Overall, the proportion of language genes ranges from 56.2% to 61.2%, while that of cognition genes ranges from 59.5% to 64.9%. The language-gene proportion peaks at position 383 (61.2%), and the cognition-gene proportion peaks at position 386 (64.9%). These curves reflect the different dynamics of language and cognition genes along the evolutionary axis, providing quantitative evidence for the co-ordination and divergence of the two gene categories (raw data are given in Table 8).

**Figure 10.**
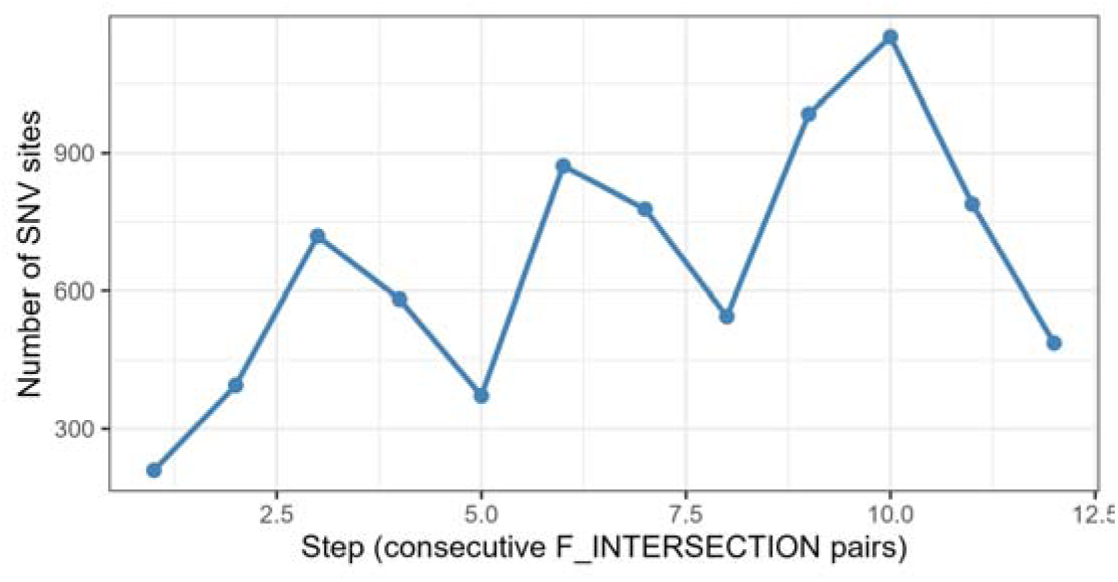
New SNV sites gained in consecutive F_INTERSECTION differences. The horizontal axis represents the step number (1–12), corresponding to the adjacent difference sets between the 13 F_INTERSECTION sets; the vertical axis represents the number of SNV loci (integer) contained in each difference set. This figure shows the trend in the number of newly acquired differential loci between adjacent peak groups.

**Table 9.**
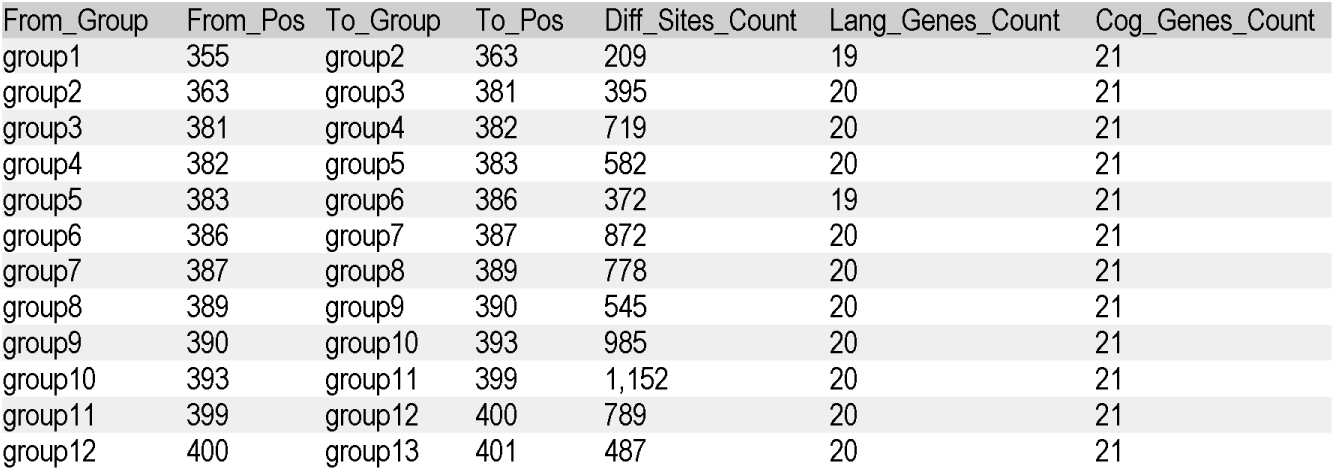
Statistics of differential loci between consecutive F_INTERSECTION sets.

Figure 11 shows the differences in activity levels of various genes during evolution. For example, *ATP2C2, EP300, NOTCH2NLA, SRGAP2C, TPK1*, and *GNB5* all have relatively high counts across multiple steps, whereas some genes appear only at specific steps, reflecting a potential division of labour between core genes and stage-specific genes. Among them, the NBPF8 gene stands out. The NBPF gene family, characterized by human-specific Olduvai (DUF1220) domain expansions at 1q21.1, has been hypothesized to contribute to human brain expansion and cognitive evolution. While the family-wide copy-number expansion is supported by comparative genomic and cellular studies, NBPF8, one member of this family, lacks direct experimental evidence for independent roles in human cognitive evolution. NBPF8 includes pseudogenic isoforms, and its protein-coding status and functional contribution remain unvalidated. Most evolutionary conclusions for NBPF are derived from the total Olduvai domain dosage rather than individual NBPF genes such as NBPF8 [36–38]. However, Figures 11, 12, and 13 all show that this gene exhibits a gradually increasing number of SNVs across different evolutionary stages of human cognitive function SNV polymorphism patterns, providing new evidence for an association between NBPF8 gene and cognitive evolution.

**Figure 11.**
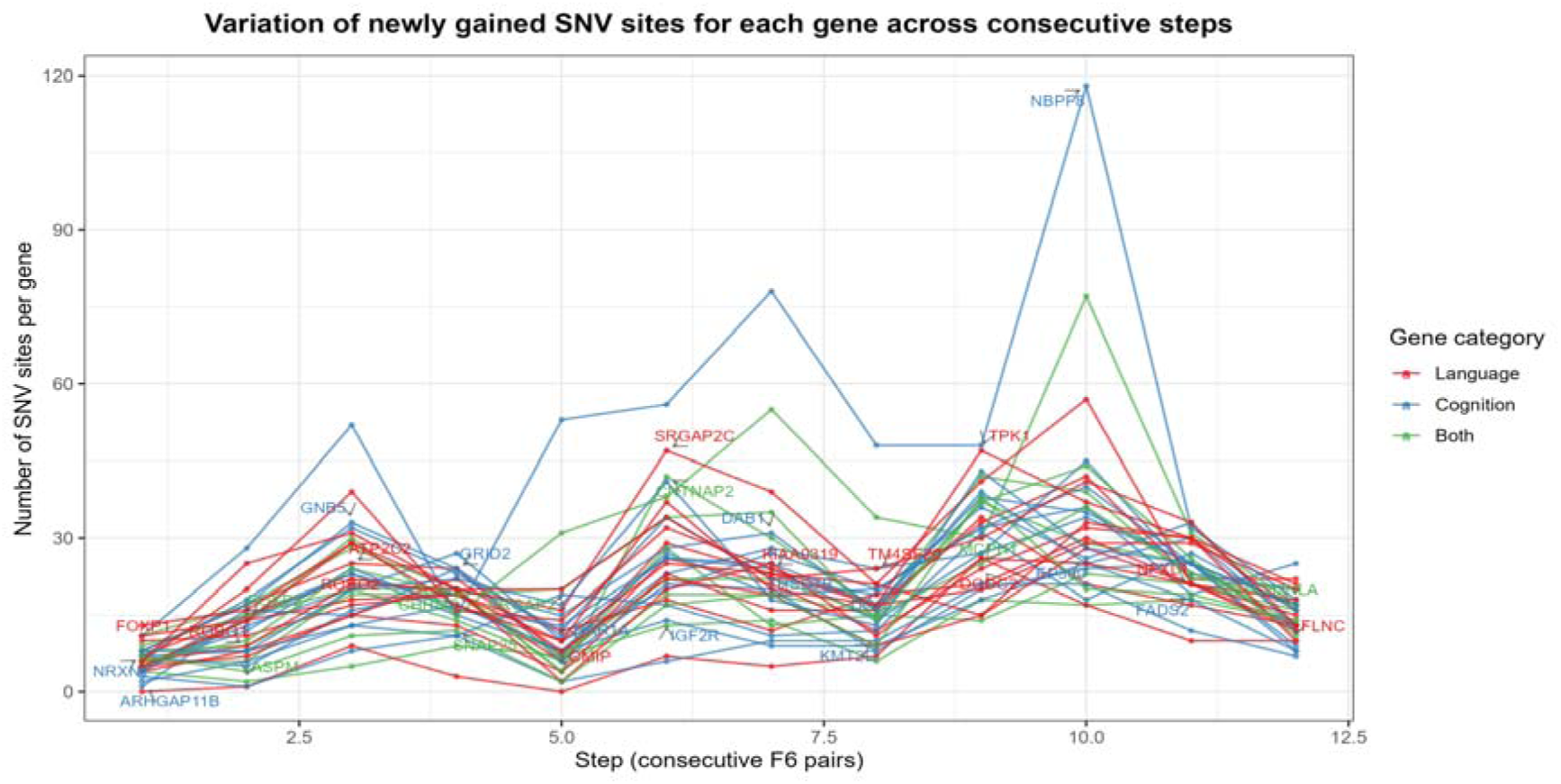
Variation of newly gained SNV sites for each gene across consecutive steps. The horizontal axis represents the step number (1–12), and the vertical axis represents the number of SNV loci (integer) for each gene that appear in the difference set at that step. A total of 33 lines are shown, representing the 33 language/cognition genes; red indicates language genes, green indicates cognition genes, and blue indicates genes that belong to both categories. Each line is labelled with the corresponding gene name.

**Figure 12.**
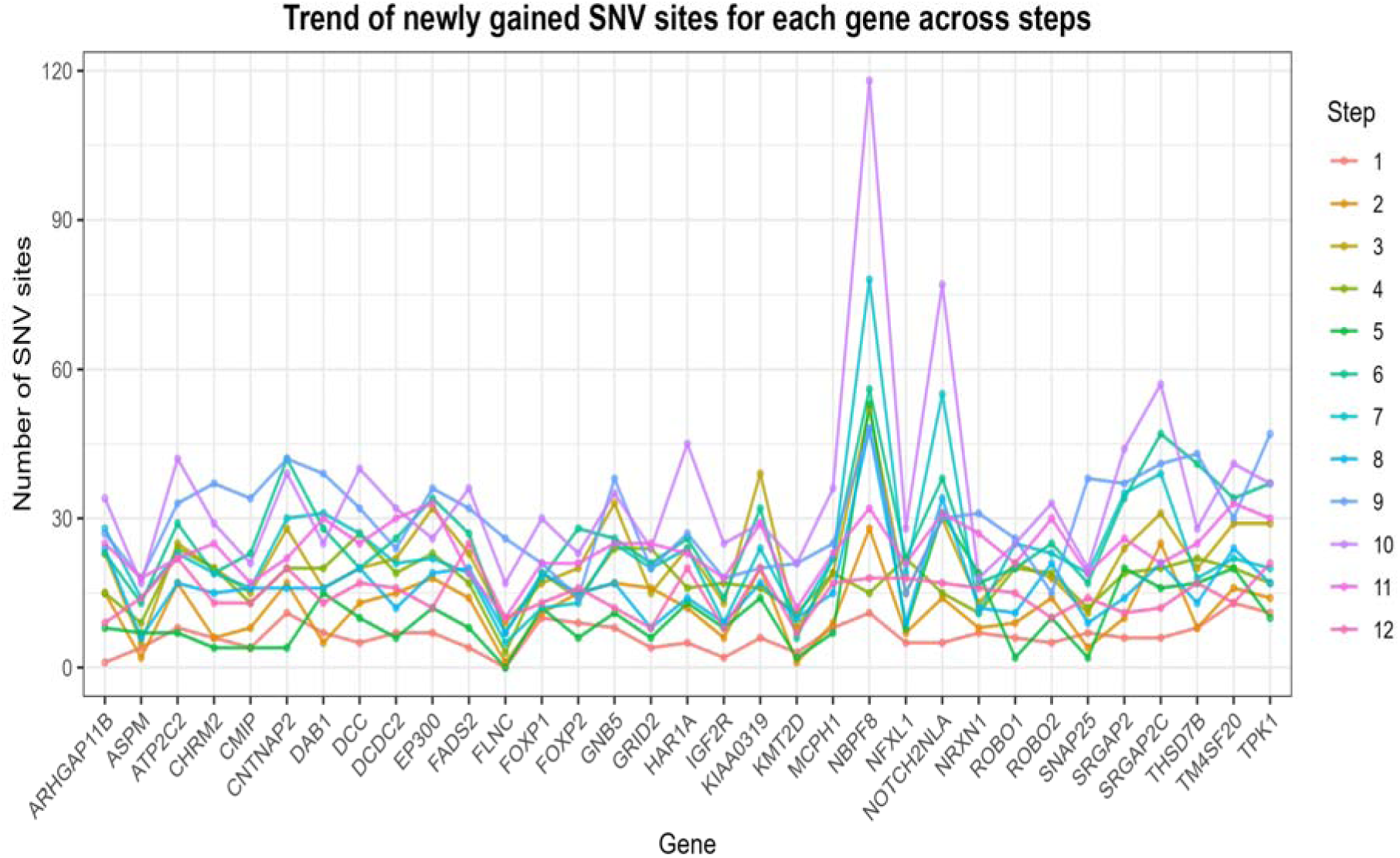
Trend of newly gained SNV sites for each gene across steps. The horizontal axis represents the 33 genes (sorted alphabetically), and the vertical axis represents the number of SNV loci (integer). Each line corresponds to one step (12 steps in total). This figure is a transpose of Figure 11, facilitating the observation of fluctuations for each gene across different steps. It can be seen that certain genes show relatively high and stable counts across multiple steps, while others exhibit peaks only at specific steps. This pattern suggests that the language-cognition gene network comprises both conserved core members and dynamically regulated auxiliary members.

**Figure 13.**
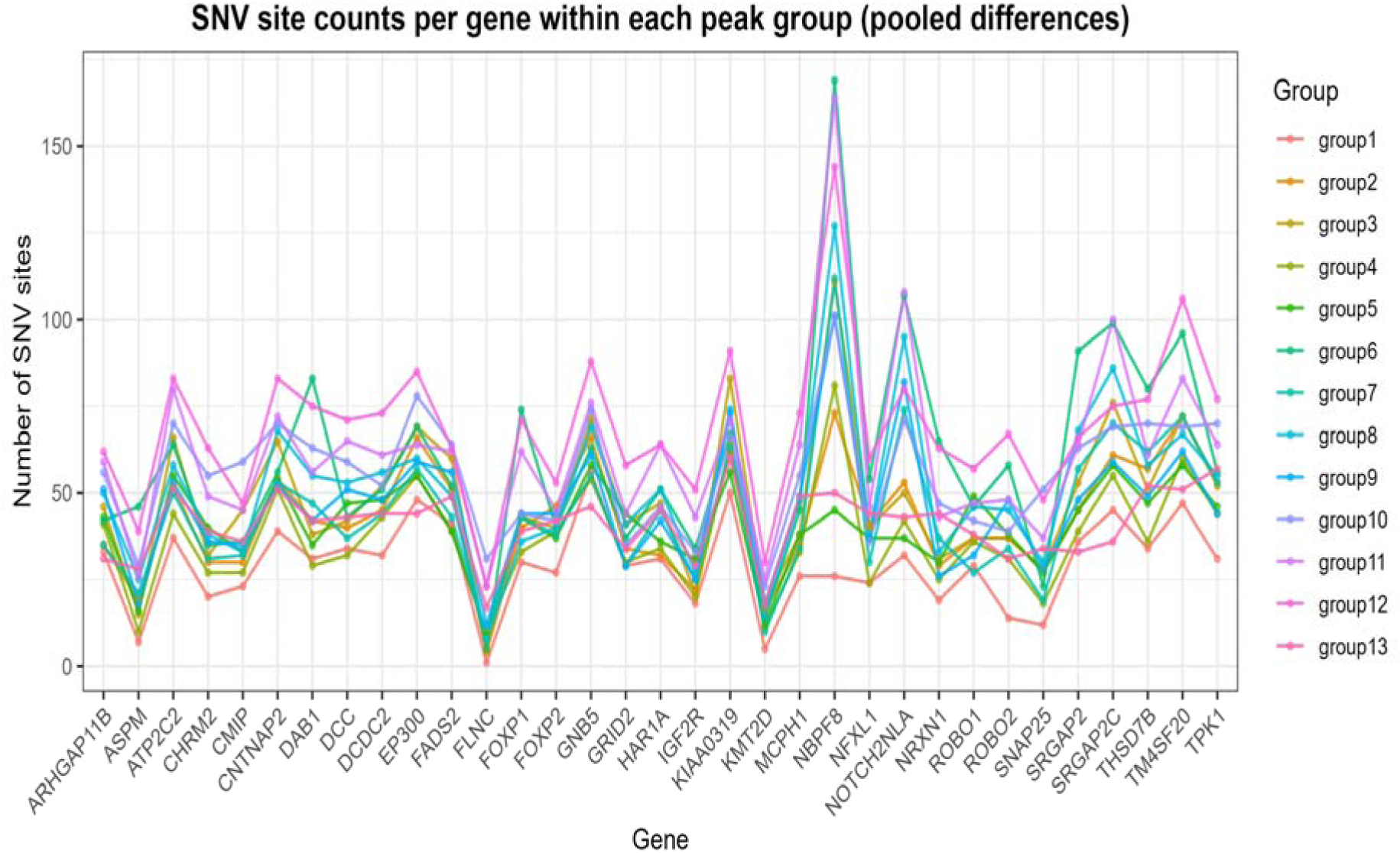
SNV site counts per gene within each peak group (pooled differences). The horizontal axis represents the 33 genes, and the vertical axis represents the number of SNV loci (integer). Each line corresponds to one peak group (13 groups in total). Unlike Figure 12, this figure uses the union of differential loci from all peaks within each group, rather than the adjacent F_INTERSECTION difference sets; thus, it reflects the cumulative differential gene counts at each breakpoint position (group) itself. It can be observed that the profiles of gene counts are similar across groups but vary in magnitude, indicating that core genes contribute at all breakpoints, yet some groups show higher overall counts, which may correspond to more drastic genetic changes. Given that from group1 to group13 along the horizontal axis of the similarity curve essentially represents the temporal direction of evolution, Figure 13 implies that the macro-evolutionary rate of cognition-gene polymorphism patterns is accelerating over time.

#### 3.4.2 Statistical test for enrichment of language/cognition genes

To assess whether the proportions of language genes and cognition genes in F_INTERSECTION deviate from the background composition, we performed hypergeometric tests (reporting both one-sided tests for the enrichment direction and two-sided tests without a preset direction). The background gene set was defined as the deduplicated genes annotated by all SNP loci, totalling 33 genes, among which 20 are language genes, 21 are cognition genes, and 8 belong to both categories. The tests were conducted at the gene level; overlapping genes were counted separately in the two categories of analysis. It should be particularly noted that in most of the sets shown in Tables 10–13, all 33 background genes are included, or only one gene is missing; therefore, their composition is almost identical to the background. Under this setting, the tests cannot provide independent evidence of enrichment.

**Table 10.** Hypergeometric enrichment test results for language/cognition genes in F_INTERSECTION (one-sided)

| Group | Start | F_INTERSECTION<br>Total genes | Langu<br>genes | Cogni<br>genes | Common<br>genes | P(Langu<br>One-sided) | OR<br>(Langu) | CI(Langu<br>Lower bound) | CI(Langu<br>Upper bound) | P(Cogni<br>One-sided) | OR<br>(Cogni) | CI(Cogni<br>Lower bound) | CI(Cogni<br>Upper bound) |
| --- | --- | --- | --- | --- | --- | --- | --- | --- | --- | --- | --- | --- | --- |
| group1 | 355 | 33 | 20 | 21 | 8 | 1 | 0 | 0 | Inf | 1 | 0 | 0 | Inf |
| group2 | 363 | 33 | 20 | 21 | 8 | 1 | 0 | 0 | Inf | 1 | 0 | 0 | Inf |
| group3 | 381 | 33 | 20 | 21 | 8 | 1 | 0 | 0 | Inf | 1 | 0 | 0 | Inf |
| group4 | 382 | 33 | 20 | 21 | 8 | 1 | 0 | 0 | Inf | 1 | 0 | 0 | Inf |
| group5 | 383 | 33 | 20 | 21 | 8 | 1 | 0 | 0 | Inf | 1 | 0 | 0 | Inf |
| group6 | 386 | 33 | 20 | 21 | 8 | 1 | 0 | 0 | Inf | 1 | 0 | 0 | Inf |
| group7 | 387 | 33 | 20 | 21 | 8 | 1 | 0 | 0 | Inf | 1 | 0 | 0 | Inf |
| group8 | 389 | 33 | 20 | 21 | 8 | 1 | 0 | 0 | Inf | 1 | 0 | 0 | Inf |
| group9 | 390 | 33 | 20 | 21 | 8 | 1 | 0 | 0 | Inf | 1 | 0 | 0 | Inf |
| group10 | 393 | 33 | 20 | 21 | 8 | 1 | 0 | 0 | Inf | 1 | 0 | 0 | Inf |
| group11 | 399 | 33 | 20 | 21 | 8 | 1 | 0 | 0 | Inf | 1 | 0 | 0 | Inf |
| group12 | 400 | 33 | 20 | 21 | 8 | 1 | 0 | 0 | Inf | 1 | 0 | 0 | Inf |
| group13 | 401 | 33 | 20 | 21 | 8 | 1 | 0 | 0 | Inf | 1 | 0 | 0 | Inf |

**Table 11.**
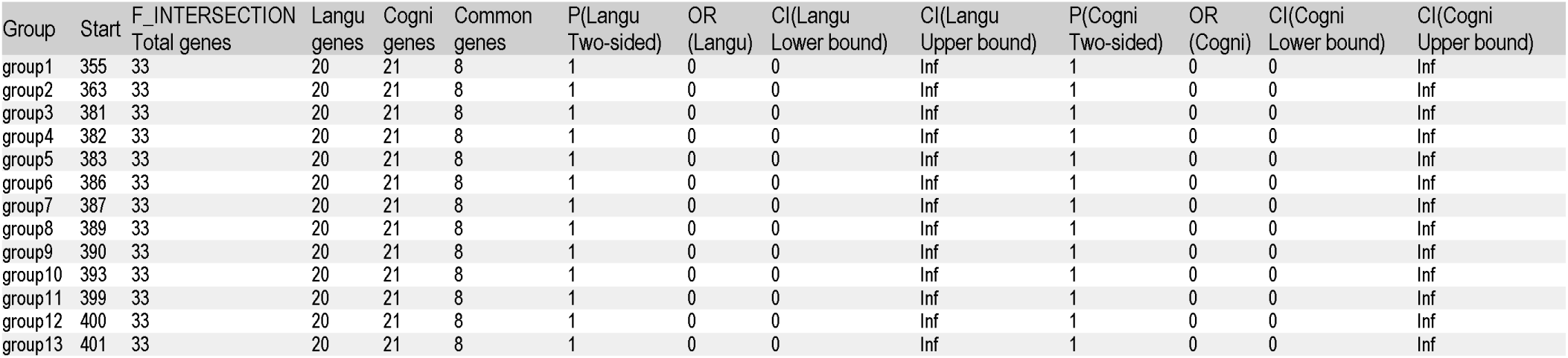
Hypergeometric enrichment test results for language/cognition genes in F_INTERSECTION (two-sided)

| Group | Start | F_INTERSECTION<br>Total genes | Langu<br>genes | Cogni<br>genes | Common<br>genes | P(Langu<br>Two-sided) | OR<br>(Langu) | CI(Langu<br>Lower bound) | CI(Langu<br>Upper bound) | P(Cogni<br>Two-sided) | OR<br>(Cogni) | CI(Cogni<br>Lower bound) | CI(Cogni<br>Upper bound) |
| --- | --- | --- | --- | --- | --- | --- | --- | --- | --- | --- | --- | --- | --- |
| group1 | 355 | 33 | 20 | 21 | 8 | 1 | 0 | 0 | Inf | 1 | 0 | 0 | Inf |
| group2 | 363 | 33 | 20 | 21 | 8 | 1 | 0 | 0 | Inf | 1 | 0 | 0 | Inf |
| group3 | 381 | 33 | 20 | 21 | 8 | 1 | 0 | 0 | Inf | 1 | 0 | 0 | Inf |
| group4 | 382 | 33 | 20 | 21 | 8 | 1 | 0 | 0 | Inf | 1 | 0 | 0 | Inf |
| group5 | 383 | 33 | 20 | 21 | 8 | 1 | 0 | 0 | Inf | 1 | 0 | 0 | Inf |
| group6 | 386 | 33 | 20 | 21 | 8 | 1 | 0 | 0 | Inf | 1 | 0 | 0 | Inf |
| group7 | 387 | 33 | 20 | 21 | 8 | 1 | 0 | 0 | Inf | 1 | 0 | 0 | Inf |
| group8 | 389 | 33 | 20 | 21 | 8 | 1 | 0 | 0 | Inf | 1 | 0 | 0 | Inf |
| group9 | 390 | 33 | 20 | 21 | 8 | 1 | 0 | 0 | Inf | 1 | 0 | 0 | Inf |
| group10 | 393 | 33 | 20 | 21 | 8 | 1 | 0 | 0 | Inf | 1 | 0 | 0 | Inf |
| group11 | 399 | 33 | 20 | 21 | 8 | 1 | 0 | 0 | Inf | 1 | 0 | 0 | Inf |
| group12 | 400 | 33 | 20 | 21 | 8 | 1 | 0 | 0 | Inf | 1 | 0 | 0 | Inf |
| group13 | 401 | 33 | 20 | 21 | 8 | 1 | 0 | 0 | Inf | 1 | 0 | 0 | Inf |

**Table 12.** Enrichment test results for language/cognition genes in adjacent F_INTERSECTION difference sets (one-sided)

| Step | Group<br>from | To<br>group | Differential<br>genes total | Langu<br>genes | Cogni<br>genes | Common<br>genes | P(Langu<br>One-sided) | OR<br>(Langu) | CI(Langu<br>Lower bound) | CI(Langu<br>Upper bound) | P(Cogni<br>One-sided) | OR<br>(Cogni) | CI(Cogni<br>Lower bound) | CI(Cogni<br>Upper bound) |
| --- | --- | --- | --- | --- | --- | --- | --- | --- | --- | --- | --- | --- | --- | --- |
| 1 | group1 | group2 | 32 | 19 | 21 | 8 | 1 | 0 | 0 | Inf | 0.3636364 | Inf | 0.09210247 | Inf |
| 2 | group2 | group3 | 33 | 20 | 21 | 8 | 1 | 0 | 0 | Inf | 1.0000000 | 0 | 0.00000000 | Inf |
| 3 | group3 | group4 | 33 | 20 | 21 | 8 | 1 | 0 | 0 | Inf | 1.0000000 | 0 | 0.00000000 | Inf |
| 4 | group4 | group5 | 33 | 20 | 21 | 8 | 1 | 0 | 0 | Inf | 1.0000000 | 0 | 0.00000000 | Inf |
| 5 | group5 | group6 | 32 | 19 | 21 | 8 | 1 | 0 | 0 | Inf | 0.3636364 | Inf | 0.09210247 | Inf |
| 6 | group6 | group7 | 33 | 20 | 21 | 8 | 1 | 0 | 0 | Inf | 1.0000000 | 0 | 0.00000000 | Inf |
| 7 | group7 | group8 | 33 | 20 | 21 | 8 | 1 | 0 | 0 | Inf | 1.0000000 | 0 | 0.00000000 | Inf |
| 8 | group8 | group9 | 33 | 20 | 21 | 8 | 1 | 0 | 0 | Inf | 1.0000000 | 0 | 0.00000000 | Inf |
| 9 | group9 | group10 | 33 | 20 | 21 | 8 | 1 | 0 | 0 | Inf | 1.0000000 | 0 | 0.00000000 | Inf |
| 10 | group10 | group11 | 33 | 20 | 21 | 8 | 1 | 0 | 0 | Inf | 1.0000000 | 0 | 0.00000000 | Inf |
| 11 | group11 | group12 | 33 | 20 | 21 | 8 | 1 | 0 | 0 | Inf | 1.0000000 | 0 | 0.00000000 | Inf |
| 12 | group12 | group13 | 33 | 20 | 21 | 8 | 1 | 0 | 0 | Inf | 1.0000000 | 0 | 0.00000000 | Inf |

**Table 13.** Enrichment test results for language/cognition genes in adjacent F_INTERSECTION difference sets (two-sided)

| Step | Group from | To group | Differential genes total | Langu genes | Cogni genes | Common genes | P(Langu two-sided) | OR (Langu) | CI(Langu Lower bound) | CI(Langu Upper bound) | P(Cogni two-sided) | OR (Cogni) | CI(Cogni Lower bound) | CI(Cogni Upper bound) |
| --- | --- | --- | --- | --- | --- | --- | --- | --- | --- | --- | --- | --- | --- | --- |
| 1 | group1 | group2 | 32 | 19 | 21 | 8 | 1 | 0 | 0 | 59.94474 | 0.3636364 | Inf | 0.0448716 | Inf |
| 2 | group2 | group3 | 33 | 20 | 21 | 8 | 1 | 0 | 0 | Inf | 1.0000000 | 0 | 0.0000000 | Inf |
| 3 | group3 | group4 | 33 | 20 | 21 | 8 | 1 | 0 | 0 | Inf | 1.0000000 | 0 | 0.0000000 | Inf |
| 4 | group4 | group5 | 33 | 20 | 21 | 8 | 1 | 0 | 0 | Inf | 1.0000000 | 0 | 0.0000000 | Inf |
| 5 | group5 | group6 | 32 | 19 | 21 | 8 | 1 | 0 | 0 | 59.94474 | 0.3636364 | Inf | 0.0448716 | Inf |
| 6 | group6 | group7 | 33 | 20 | 21 | 8 | 1 | 0 | 0 | Inf | 1.0000000 | 0 | 0.0000000 | Inf |
| 7 | group7 | group8 | 33 | 20 | 21 | 8 | 1 | 0 | 0 | Inf | 1.0000000 | 0 | 0.0000000 | Inf |
| 8 | group8 | group9 | 33 | 20 | 21 | 8 | 1 | 0 | 0 | Inf | 1.0000000 | 0 | 0.0000000 | Inf |
| 9 | group9 | group10 | 33 | 20 | 21 | 8 | 1 | 0 | 0 | Inf | 1.0000000 | 0 | 0.0000000 | Inf |
| 10 | group10 | group11 | 33 | 20 | 21 | 8 | 1 | 0 | 0 | Inf | 1.0000000 | 0 | 0.0000000 | Inf |
| 11 | group11 | group12 | 33 | 20 | 21 | 8 | 1 | 0 | 0 | Inf | 1.0000000 | 0 | 0.0000000 | Inf |
| 12 | group12 | group13 | 33 | 20 | 21 | 8 | 1 | 0 | 0 | Inf | 1.0000000 | 0 | 0.0000000 | Inf |

Each F_INTERSECTION group in Table 10 contains all 33 background genes, and the numbers of language genes and cognition genes are also completely consistent with the background values. Consequently, all one-sided P-values are 1, indicating no positive enrichment of either language genes or cognition genes.

The two-sided tests in Table 11 are consistent with the one-sided test conclusions: the gene sets in each group are completely identical in composition to the background, and all two-sided P-values are 1. This result does not support any significant deviation in the proportions of language or cognition genes at any breakpoint, nor can it be used to infer directional enrichment signals at key nodes. The adjacent difference sets in Table 12 predominantly contain all 33 background genes; only steps 1 and 5 are missing one language gene, but cognition genes remain all 21. The corresponding one-sided P-values do not reach significance (minimum cognitive P = 0.364), thus providing no statistical evidence for enrichment of language genes in early steps or cognition genes in later steps.

The two-sided results in Table 13 further indicate that the language/cognition gene compositions of the adjacent difference sets do not deviate significantly from the background; the minimum two-sided P-value is 0.364, still above 0.05. Therefore, based on the current definition of background and difference sets, these results cannot be interpreted as non-random fluctuations, cumulative small-effect events, or stage-specific enrichment.

#### 3.4.3 Hypergeometric enrichment test at the SNV locus level

We further examined whether language/cognition-associated loci are enriched in F_INTERSECTION at the SNV locus level. The background was defined as all detectable SNV loci, and the success items were loci corresponding to language or cognition genes. The same hypergeometric test (one-sided and two-sided) was applied. The total number of background loci was 13,415, of which 8,126 were language-associated, 8,555 were cognition-associated, and 3,265 belonged to both categories. The one-sided test results (Table 14) showed no evidence of enrichment. In the two-sided tests (Table 15), the raw test results indicated a depletion trend in three subsets: language genes group 6 (*P* = 0.0131, OR = 0.82, 95% CI 0.70–0.96); cognition genes group 3 (*P* = 0.0217, OR = 0.82, 95% CI 0.70–0.97) and group 5 (*P* = 0.0120, OR = 0.86, 95% CI 0.76–0.96). After further multiple-testing correction using the Benjamini–Hochberg method, the corresponding q-values for the above three groups were 0.1698, 0.1413, and 0.1413, respectively, all greater than 0.05. For the remaining subsets, the raw P-values were all >0.05, and the corrected q-values were also >0.05. In summary, Table 14 and Table 15 did not provide statistically robust evidence of enrichment or depletion; only a sample-level depletion trend was observed in a few subsets.

**Table 14.** Hypergeometric enrichment test results for language/cognition loci in F_INTERSECTION (one-sided)

| Group | Start | F_INTERSECTION Loci (total) | Langu loci | Cogni loci | Common loci | P(Langu one-sided) | OR(Langu one-sided) | CI(Langu Lower bound) | CI(Langu Upper bound) | P(Cogni one-sided) | OR(Cogni one-sided) | CI(Cogni Lower bound) | CI(Cogni Upper bound) |
| --- | --- | --- | --- | --- | --- | --- | --- | --- | --- | --- | --- | --- | --- |
| group1 | 355 | 967 | 562 | 604 | 199 | 0.9505548 | 0.8961874 | 0.8000340 | Inf | 0.8191774 | 0.9413047 | 0.8386034 | Inf |
| group2 | 363 | 584 | 357 | 362 | 135 | 0.4061192 | 1.0249084 | 0.8856440 | Inf | 0.8314192 | 0.9232944 | 0.7973325 | Inf |
| group3 | 381 | 654 | 397 | 389 | 132 | 0.4887482 | 1.0059140 | 0.8762869 | Inf | 0.9910029 | 0.8262014 | 0.7200920 | Inf |
| group4 | 382 | 1,217 | 730 | 746 | 259 | 0.6810821 | 0.9734291 | 0.8784910 | Inf | 0.9715767 | 0.8904148 | 0.8029930 | Inf |
| group5 | 383 | 1,346 | 824 | 816 | 294 | 0.3145680 | 1.0307455 | 0.9340406 | Inf | 0.9945803 | 0.8616392 | 0.7808257 | Inf |
| group6 | 386 | 737 | 414 | 478 | 155 | 0.9944391 | 0.8255337 | 0.7260230 | Inf | 0.2769454 | 1.0515380 | 0.9204894 | Inf |
| group7 | 387 | 1,447 | 853 | 921 | 327 | 0.9134241 | 0.9272138 | 0.8435313 | Inf | 0.5517334 | 0.9942845 | 0.9027036 | Inf |
| group8 | 389 | 1,702 | 1,014 | 1,100 | 412 | 0.8221008 | 0.9537081 | 0.8731363 | Inf | 0.2222400 | 1.0438951 | 0.9536288 | Inf |
| group9 | 390 | 1,514 | 904 | 967 | 357 | 0.7750585 | 0.9603174 | 0.8751151 | Inf | 0.4766880 | 1.0050596 | 0.9141427 | Inf |
| group10 | 393 | 1,832 | 1,083 | 1,185 | 436 | 0.9184548 | 0.9322728 | 0.8559599 | Inf | 0.1972730 | 1.0472300 | 0.9593562 | Inf |
| group11 | 399 | 2,021 | 1,195 | 1,291 | 465 | 0.9279959 | 0.9317915 | 0.8584730 | Inf | 0.4653049 | 1.0057328 | 0.9249579 | Inf |
| group12 | 400 | 2,266 | 1,335 | 1,436 | 505 | 0.9632460 | 0.9204239 | 0.8512101 | Inf | 0.6753814 | 0.9796398 | 0.9045711 | Inf |
| group13 | 401 | 1,350 | 823 | 846 | 319 | 0.3893333 | 1.0185173 | 0.9231782 | Inf | 0.8204832 | 0.9487051 | 0.8591126 | Inf |

**Table 15.** Hypergeometric enrichment test results for language/cognition loci in F_INTERSECTION (two-sided)

| Group | Start | F_INTERSECTION<br>Loci (total) | Langu<br>loci | Cogni<br>loci | Common<br>loci | P(Langu<br>Two-sided) | OR(Langu<br>Two-sided) | CI(Langu<br>Lower bound) | CI(Langu<br>Upper bound) | P(Cogni<br>Two-sided) | OR(Cogni<br>Two-sided) | CI(Cogni<br>Lower bound) | CI(Cogni<br>Upper bound) |
| --- | --- | --- | --- | --- | --- | --- | --- | --- | --- | --- | --- | --- | --- |
| group1 | 355 | 967 | 562 | 604 | 199 | 0.10844459 | 0.8961874 | 0.7832451 | 1.0260368 | 0.38540510 | 0.9413047 | 0.8207444 | 1.0806856 |
| group2 | 363 | 584 | 357 | 362 | 135 | 0.79516248 | 1.0249084 | 0.8620008 | 1.2204937 | 0.37871407 | 0.9232944 | 0.7759257 | 1.1004921 |
| group3 | 381 | 654 | 397 | 389 | 132 | 0.96728882 | 1.0059140 | 0.8541020 | 1.1862640 | 0.02173140 | 0.8262014 | 0.7019210 | 0.9735918 |
| group4 | 382 | 1,217 | 730 | 746 | 259 | 0.66683107 | 0.9734291 | 0.8617390 | 1.1002846 | 0.06069792 | 0.8904148 | 0.7875964 | 1.0073305 |
| group5 | 383 | 1,346 | 824 | 816 | 294 | 0.61728582 | 1.0307455 | 0.9169560 | 1.1593929 | 0.01203027 | 0.8616392 | 0.7665513 | 0.9690822 |
| group6 | 386 | 737 | 414 | 478 | 155 | 0.01305807 | 0.8255337 | 0.7088268 | 0.9620393 | 0.55439519 | 1.0515380 | 0.8980604 | 1.2333772 |
| group7 | 387 | 1,447 | 853 | 921 | 327 | 0.19021622 | 0.9272138 | 0.8286751 | 1.0379386 | 0.93079377 | 0.9942845 | 0.8865221 | 1.1159718 |
| group8 | 389 | 1,702 | 1,014 | 1,100 | 412 | 0.38112515 | 0.9537081 | 0.8587238 | 1.0595501 | 0.43401519 | 1.0438951 | 0.9375644 | 1.1630205 |
| group9 | 390 | 1,514 | 904 | 967 | 357 | 0.46806207 | 0.9603174 | 0.8599496 | 1.0729503 | 0.95473316 | 1.0050596 | 0.8980345 | 1.1256289 |
| group10 | 393 | 1,832 | 1,083 | 1,185 | 436 | 0.17280662 | 0.9322728 | 0.8422864 | 1.0321859 | 0.38814445 | 1.0472300 | 0.9436921 | 1.1628127 |
| group11 | 399 | 2,021 | 1,195 | 1,291 | 465 | 0.15211725 | 0.9317915 | 0.8453555 | 1.0274092 | 0.92001800 | 1.0057328 | 0.9104980 | 1.1115098 |
| group12 | 400 | 2,266 | 1,335 | 1,436 | 505 | 0.07709018 | 0.9204239 | 0.8387785 | 1.0102935 | 0.66621487 | 0.9796398 | 0.8911205 | 1.0774210 |
| group13 | 401 | 1,350 | 823 | 846 | 319 | 0.76908295 | 1.0185173 | 0.9063191 | 1.1452844 | 0.38664131 | 0.9487051 | 0.8432909 | 1.0680583 |

#### 3.4.4 Locus frequency enrichment analysis

We further counted the frequency of each locus appearing across all 13 F_INTERSECTION groups, and used a binomial test to determine which loci appeared significantly more often than expected by chance (assuming the probability of each locus being selected equals the average group size divided by the total number of loci), followed by FDR correction (Table 16).

**Table 16.**
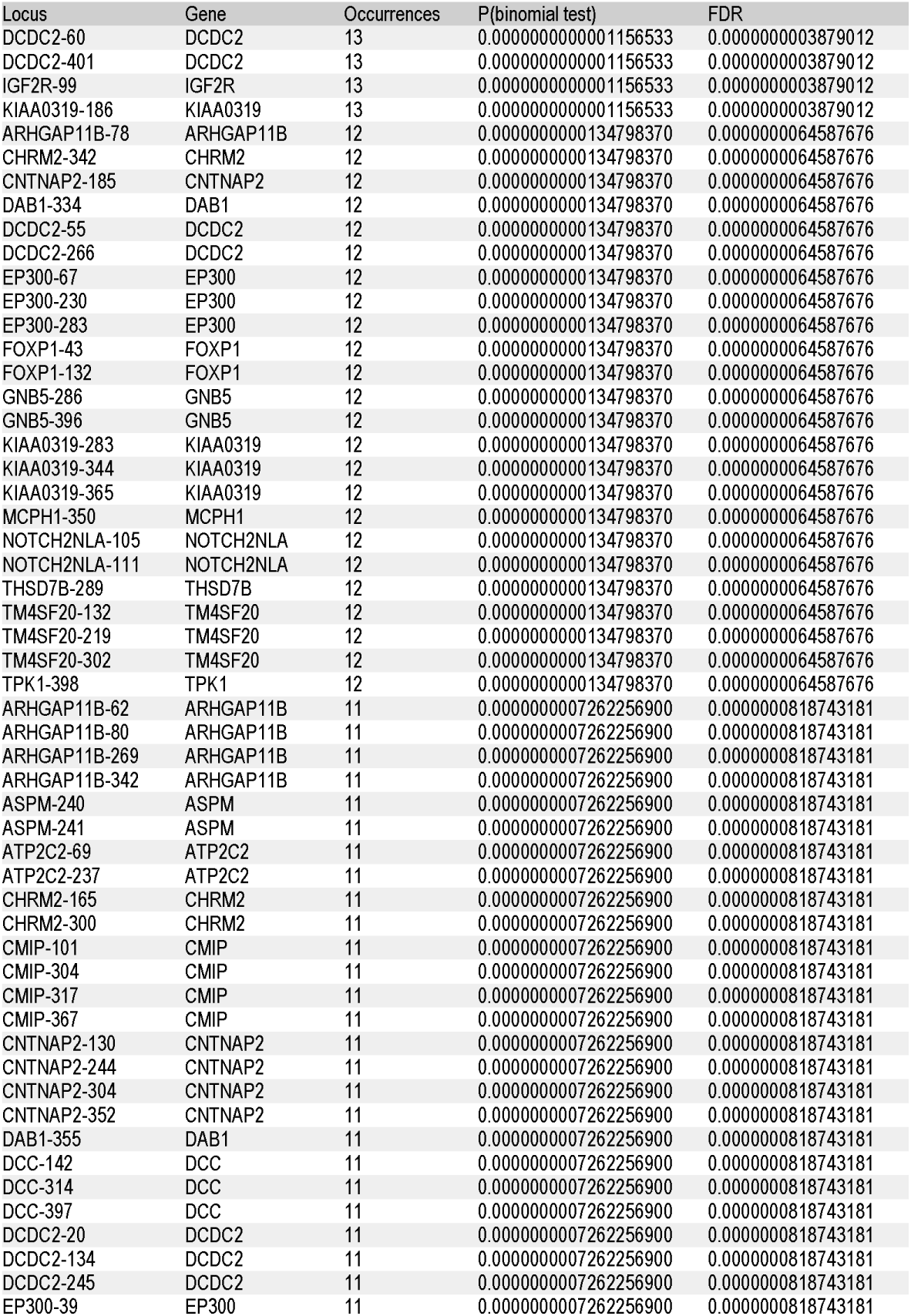
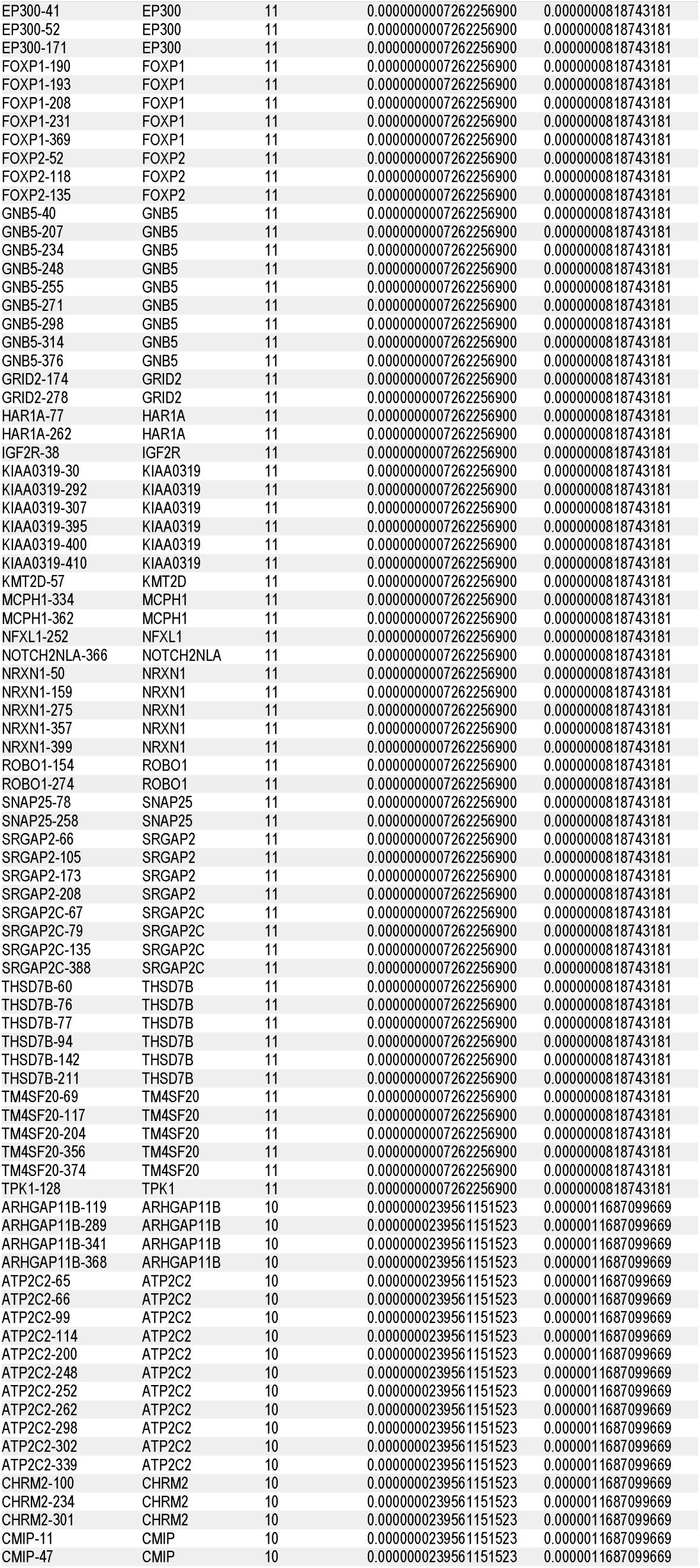
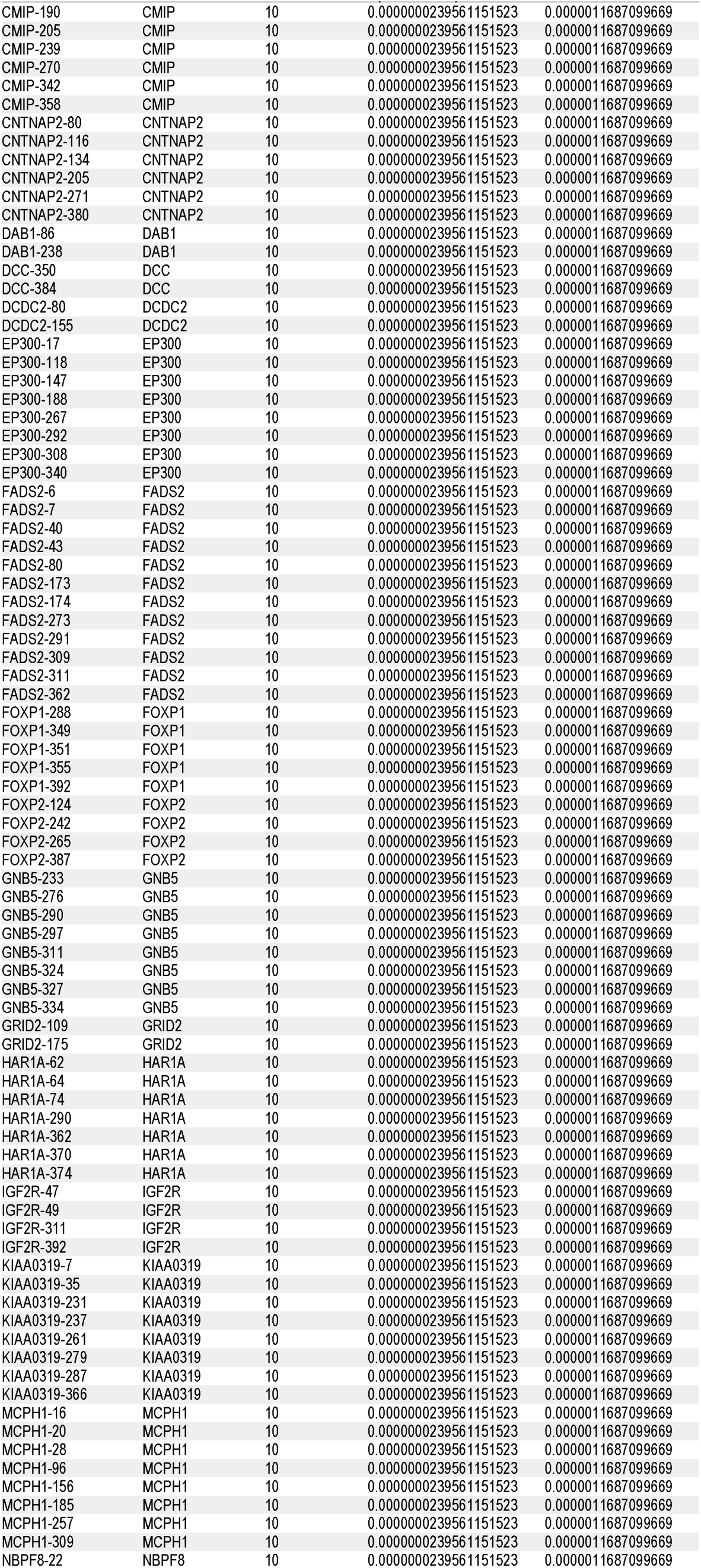
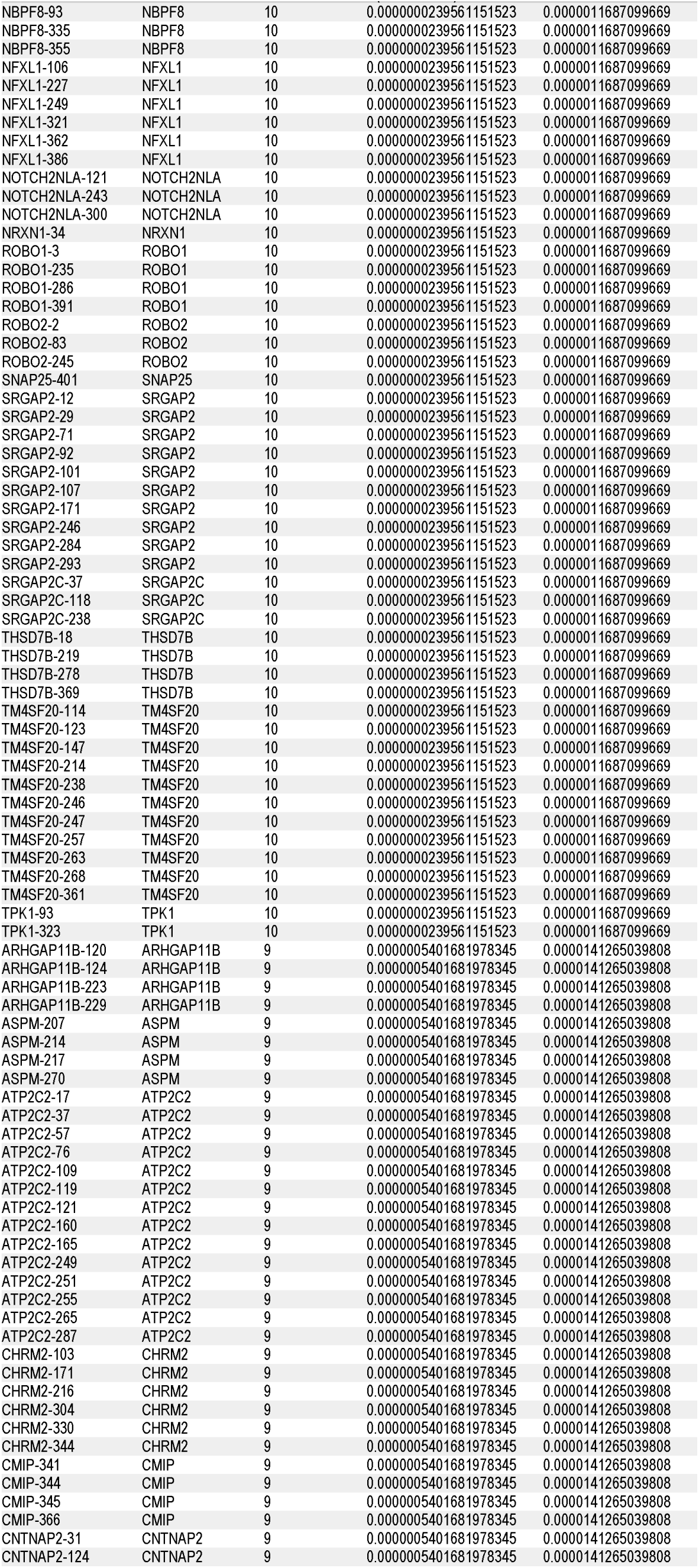
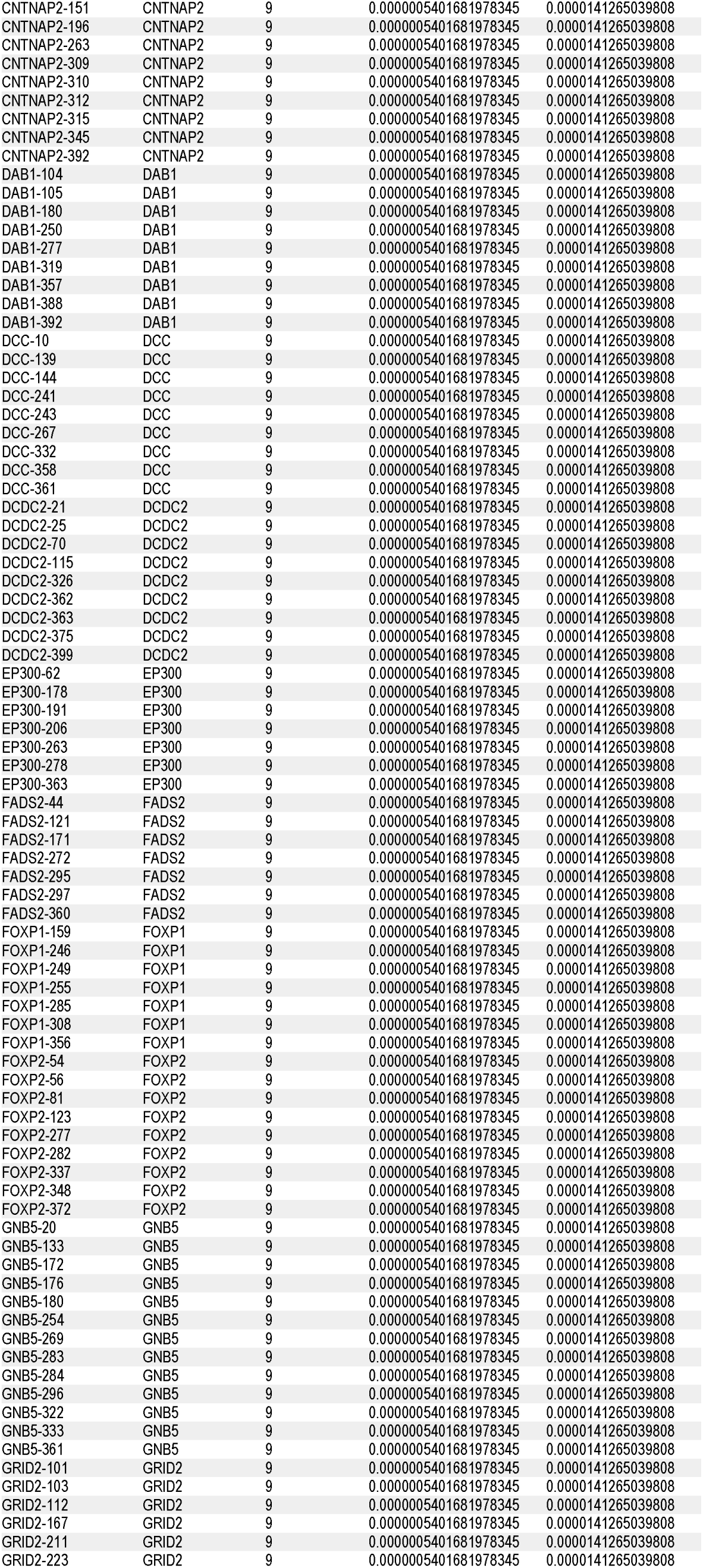
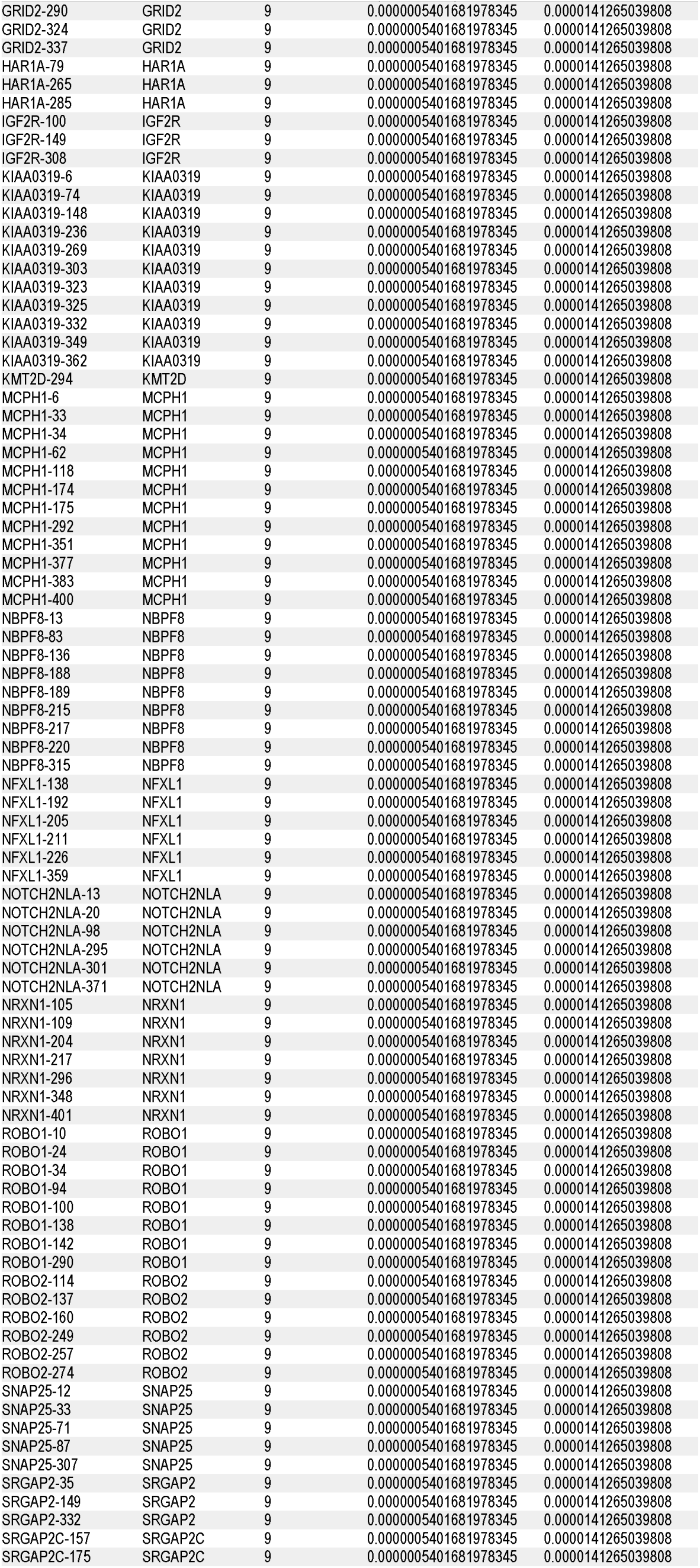
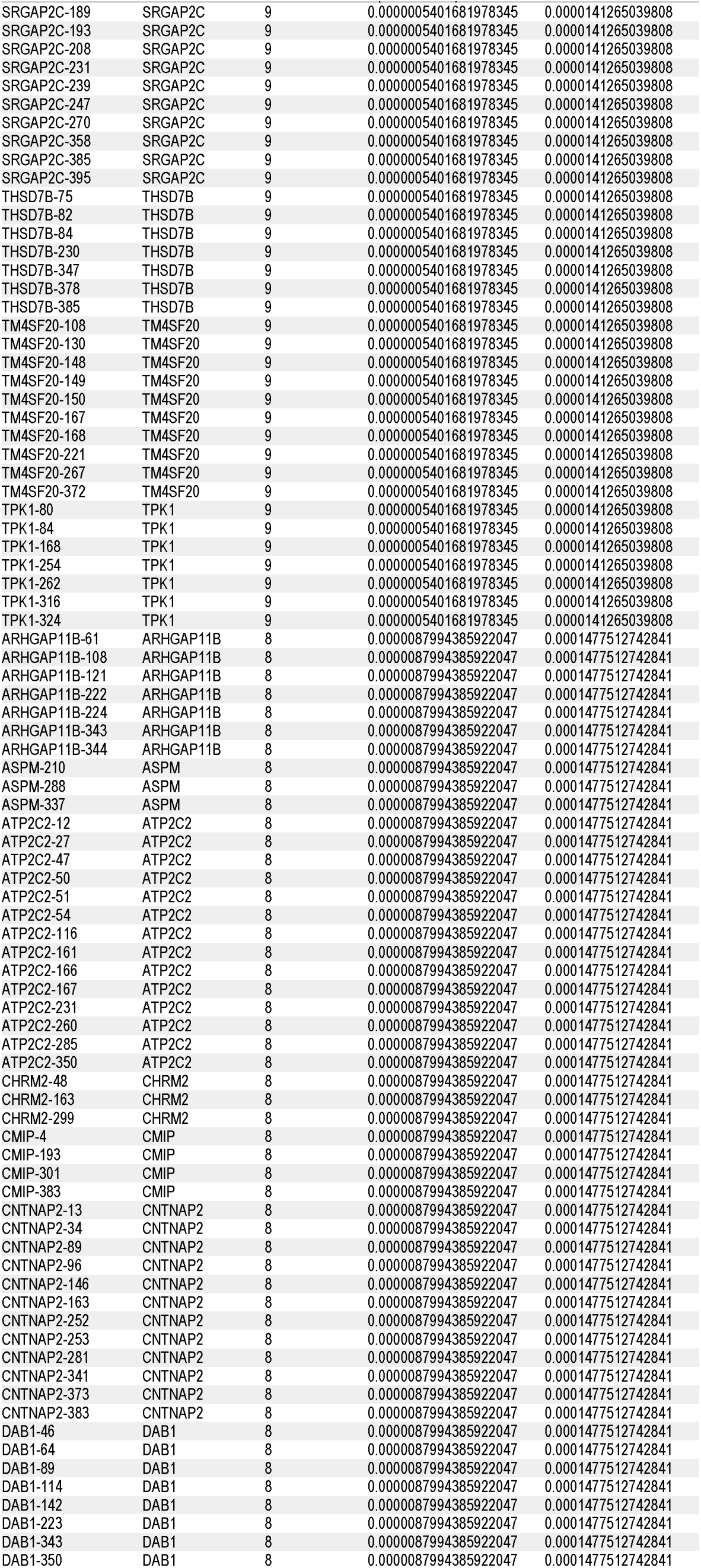
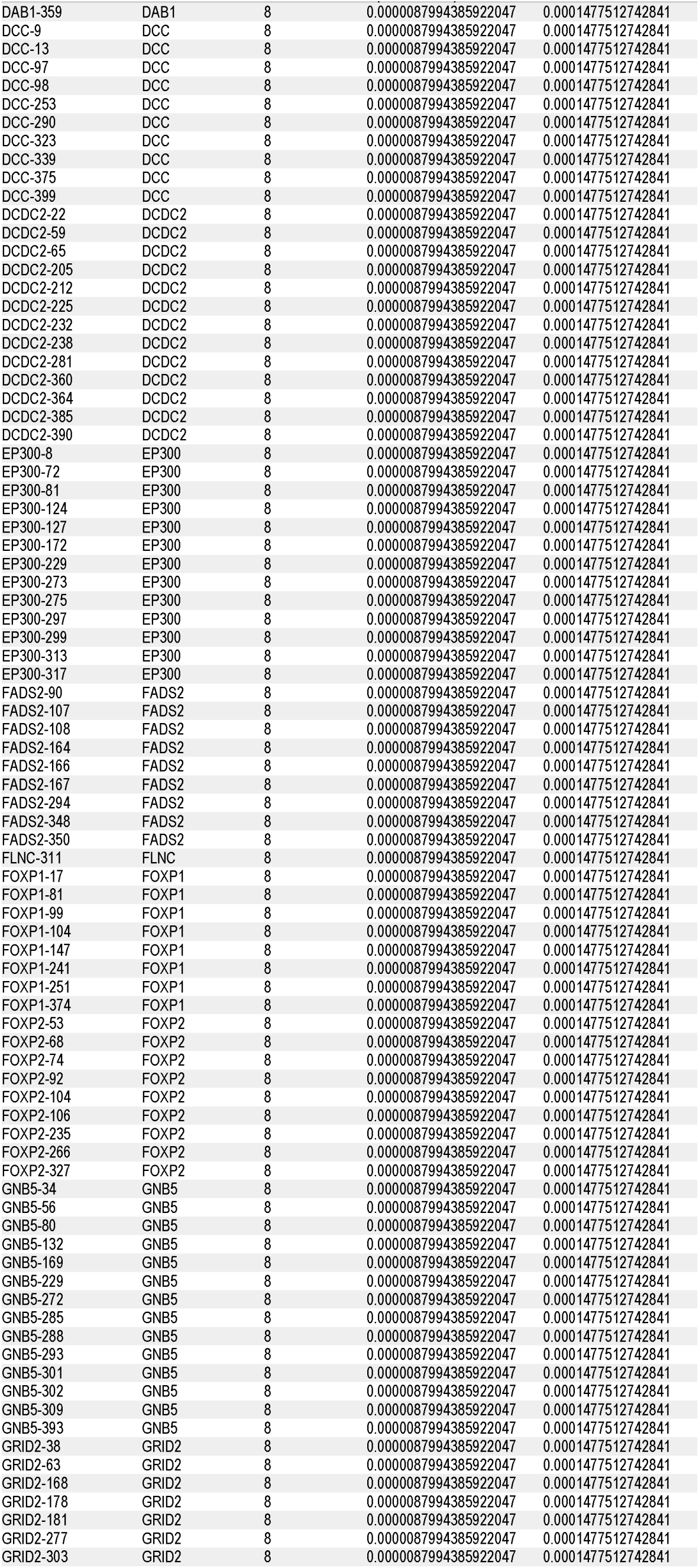
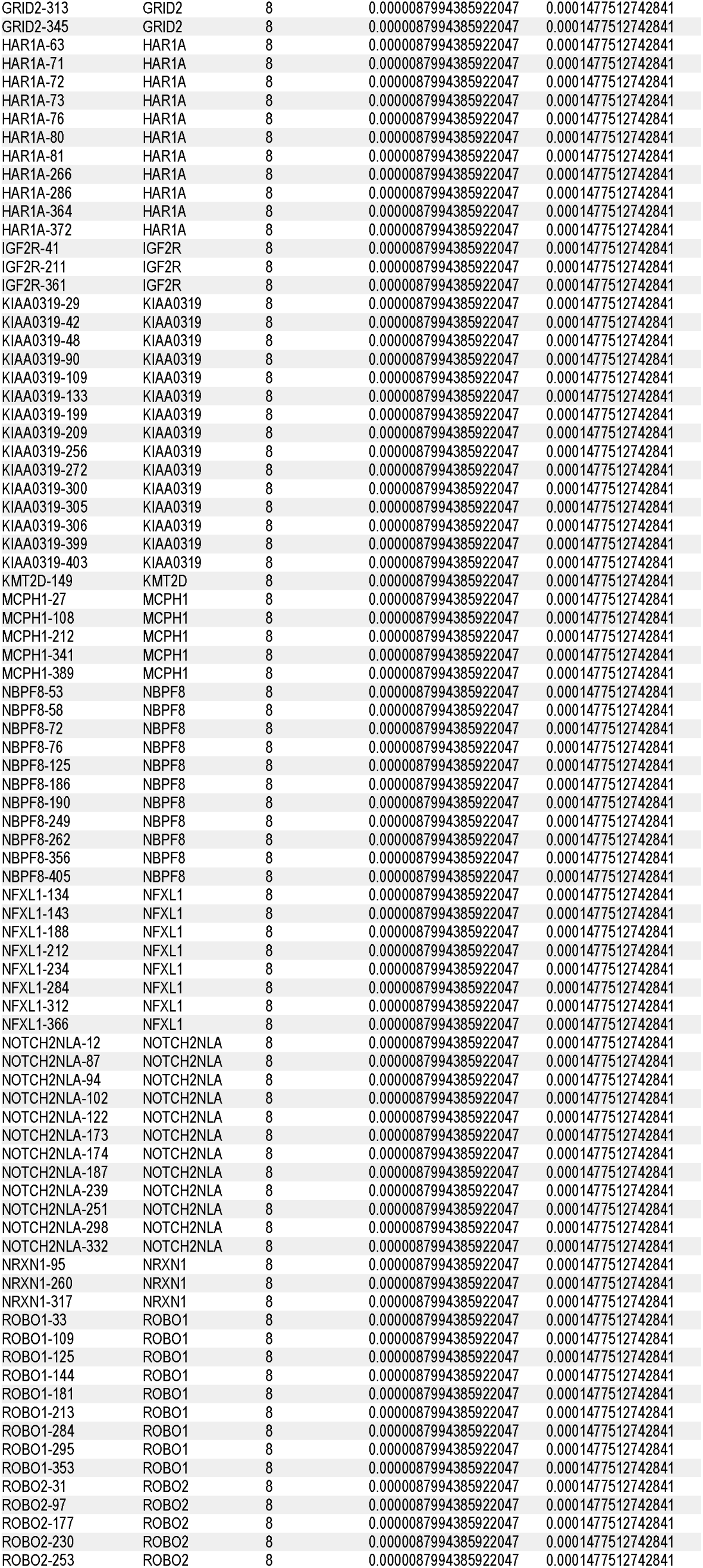
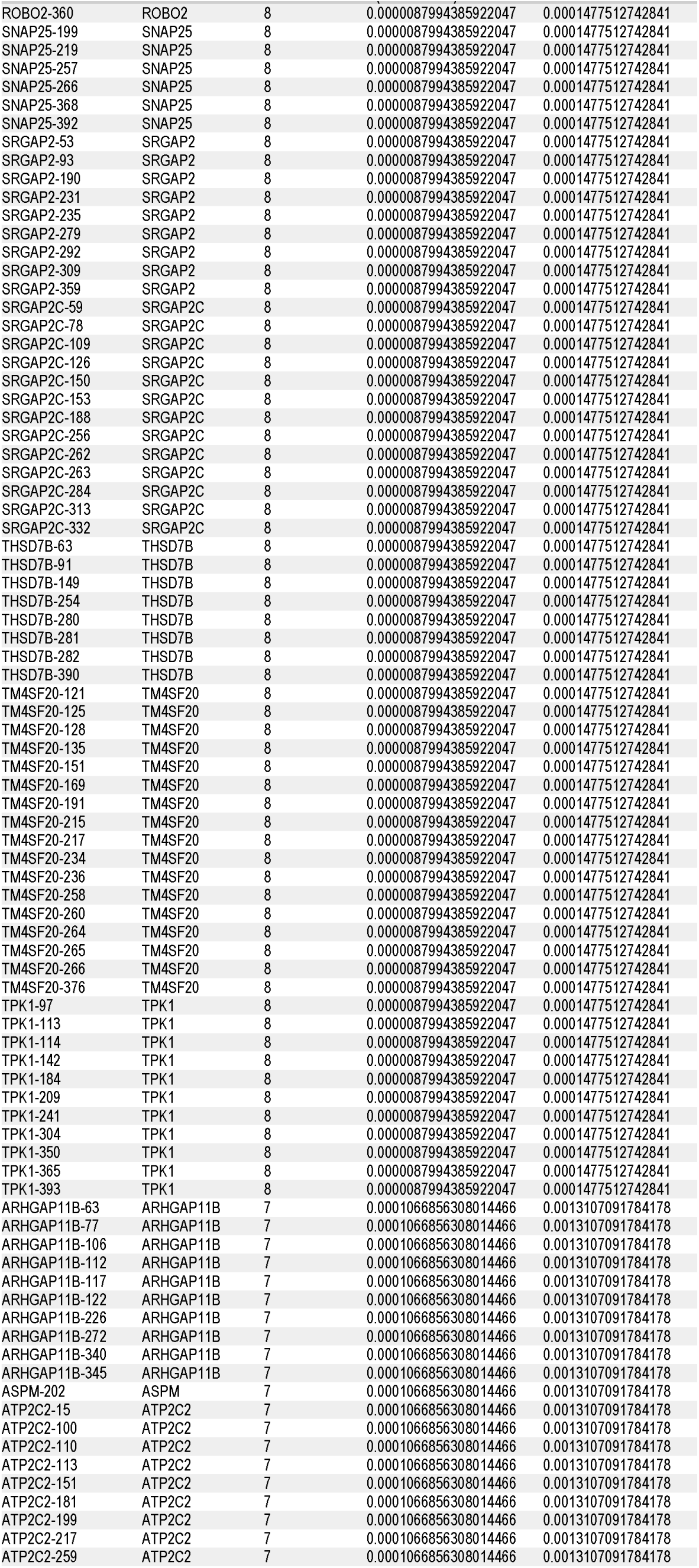
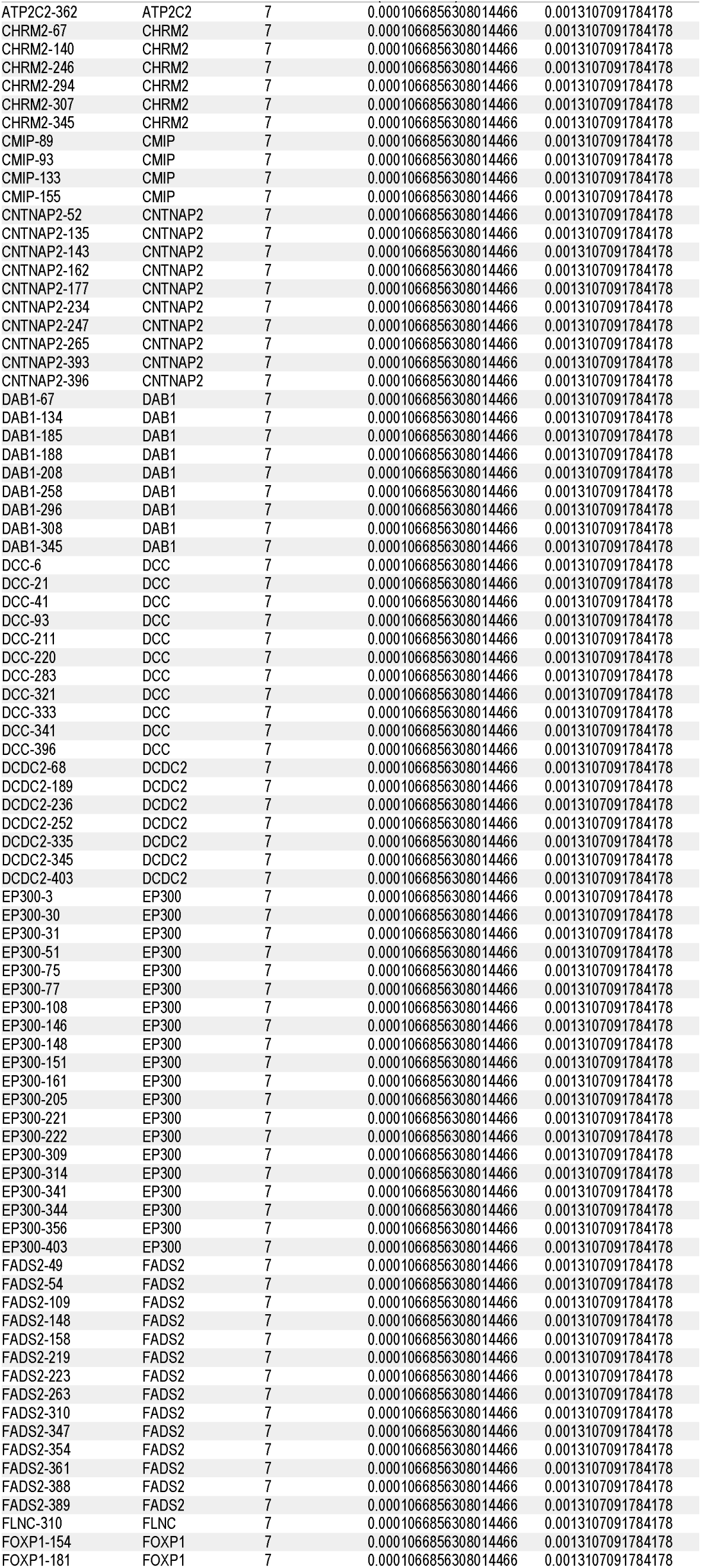
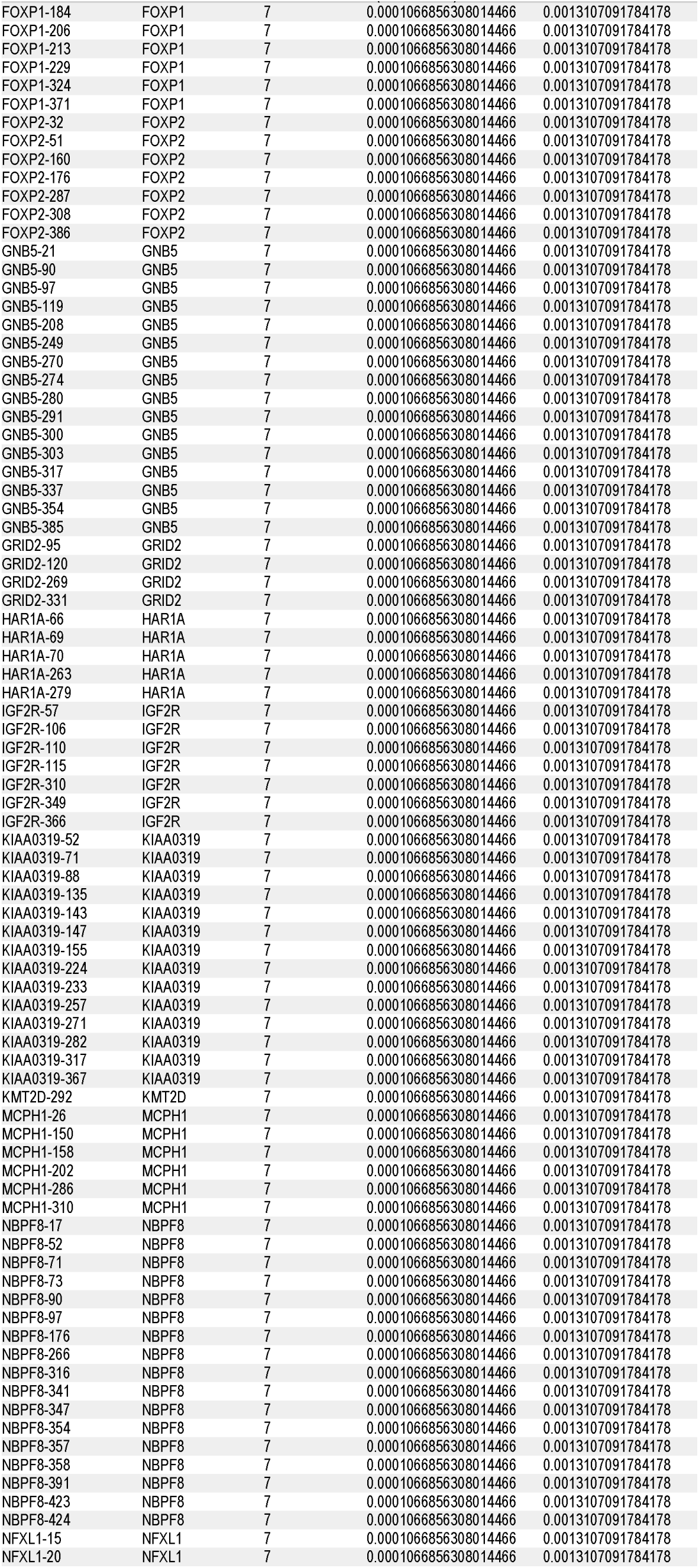
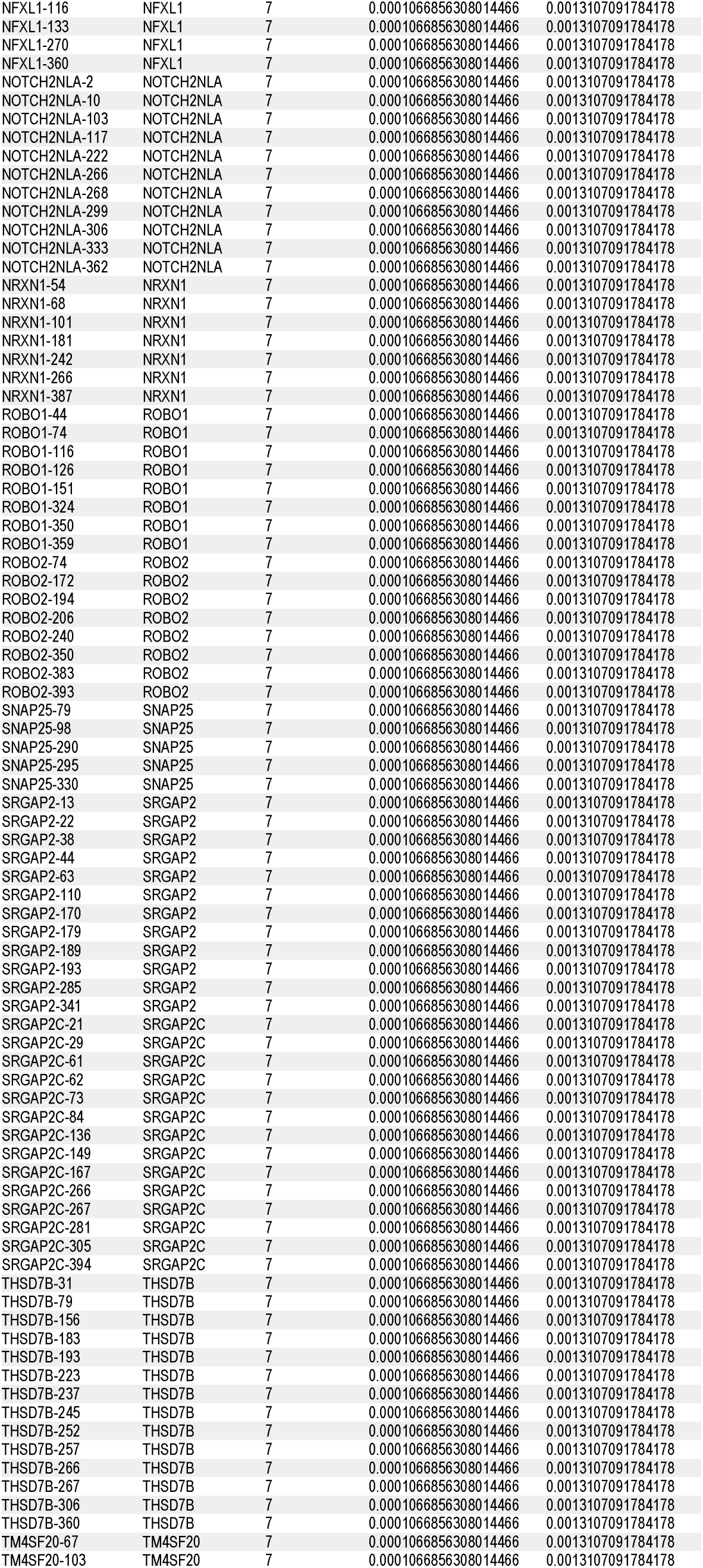
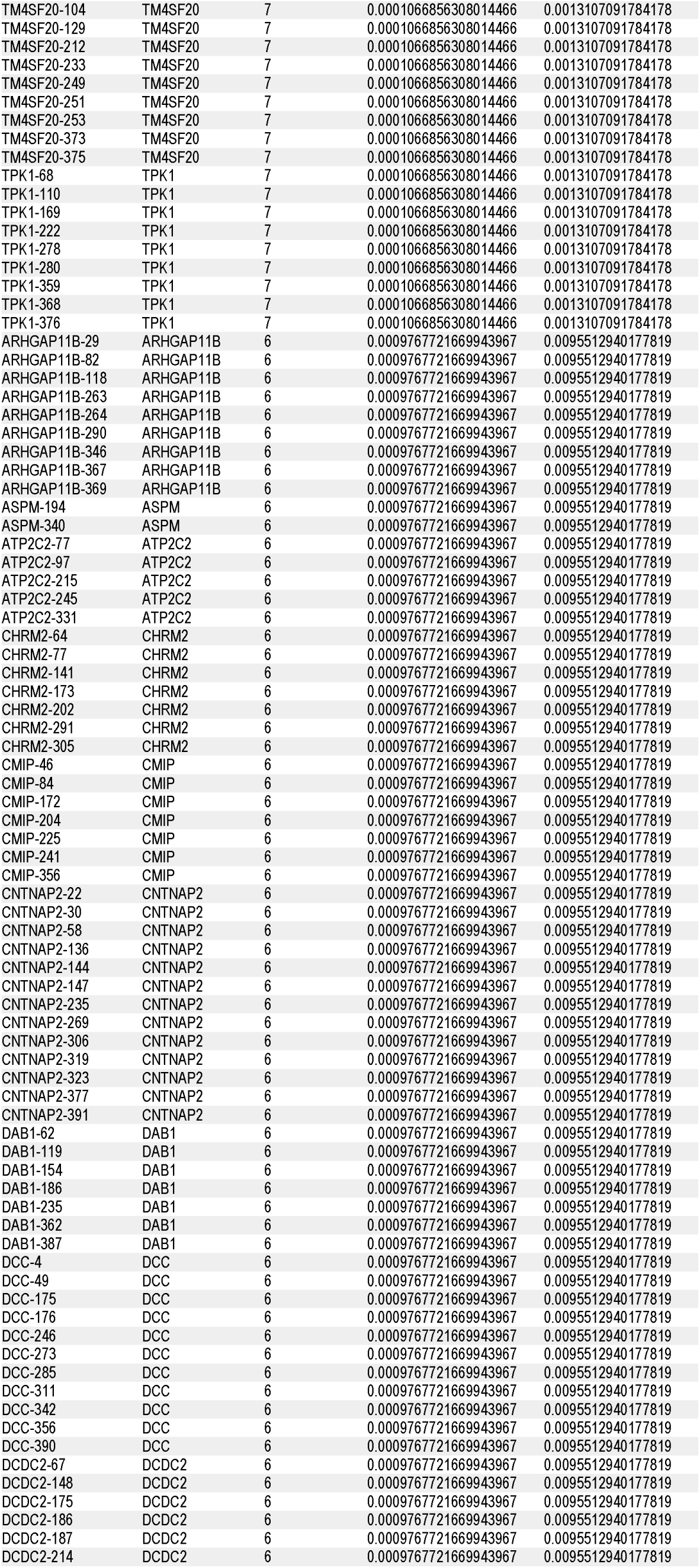
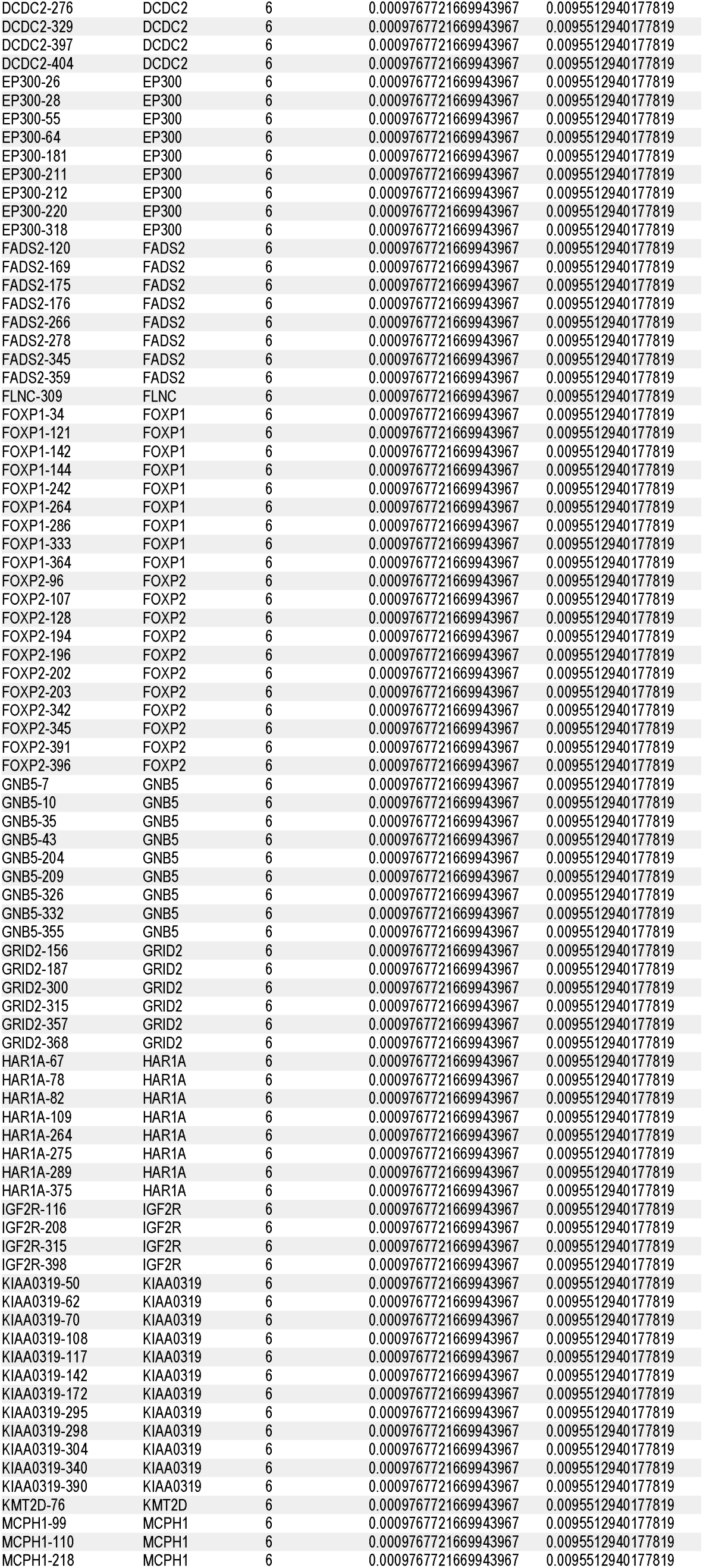
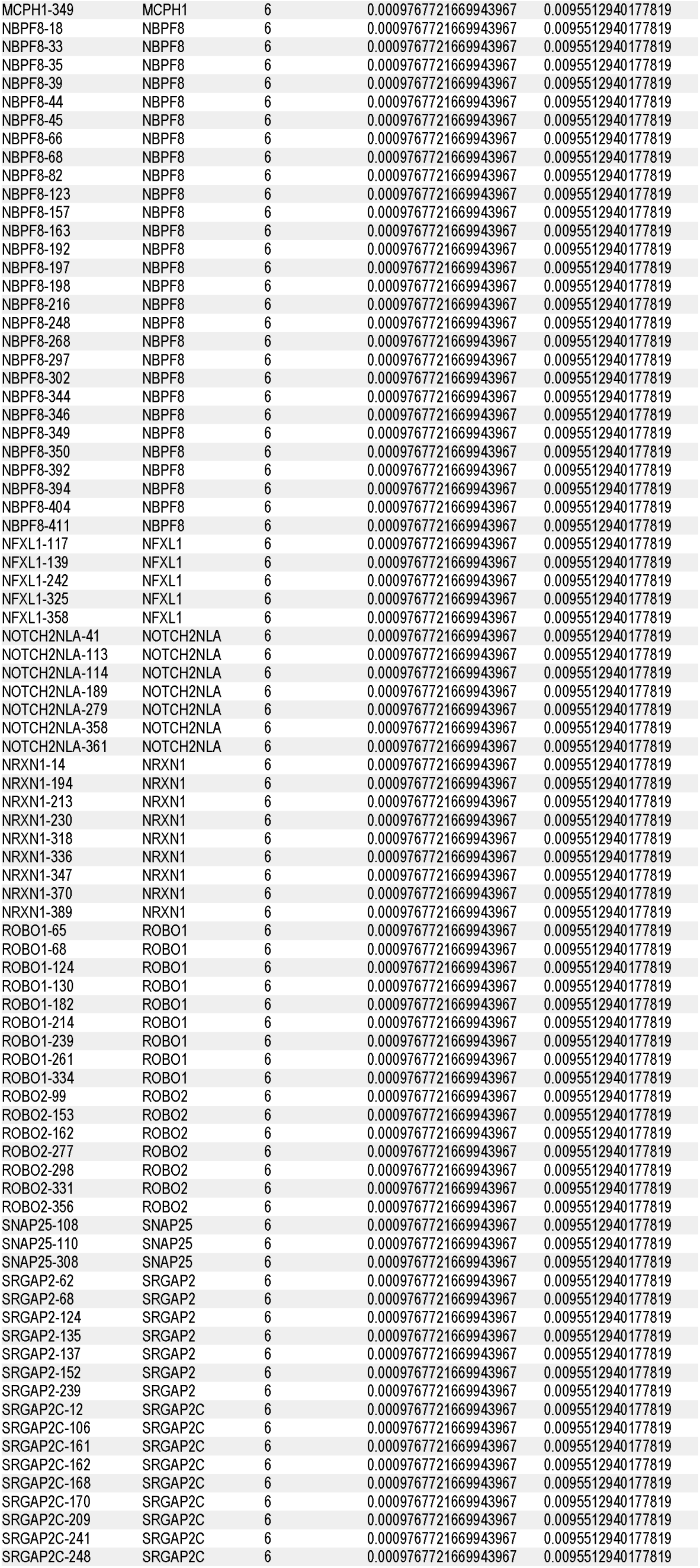
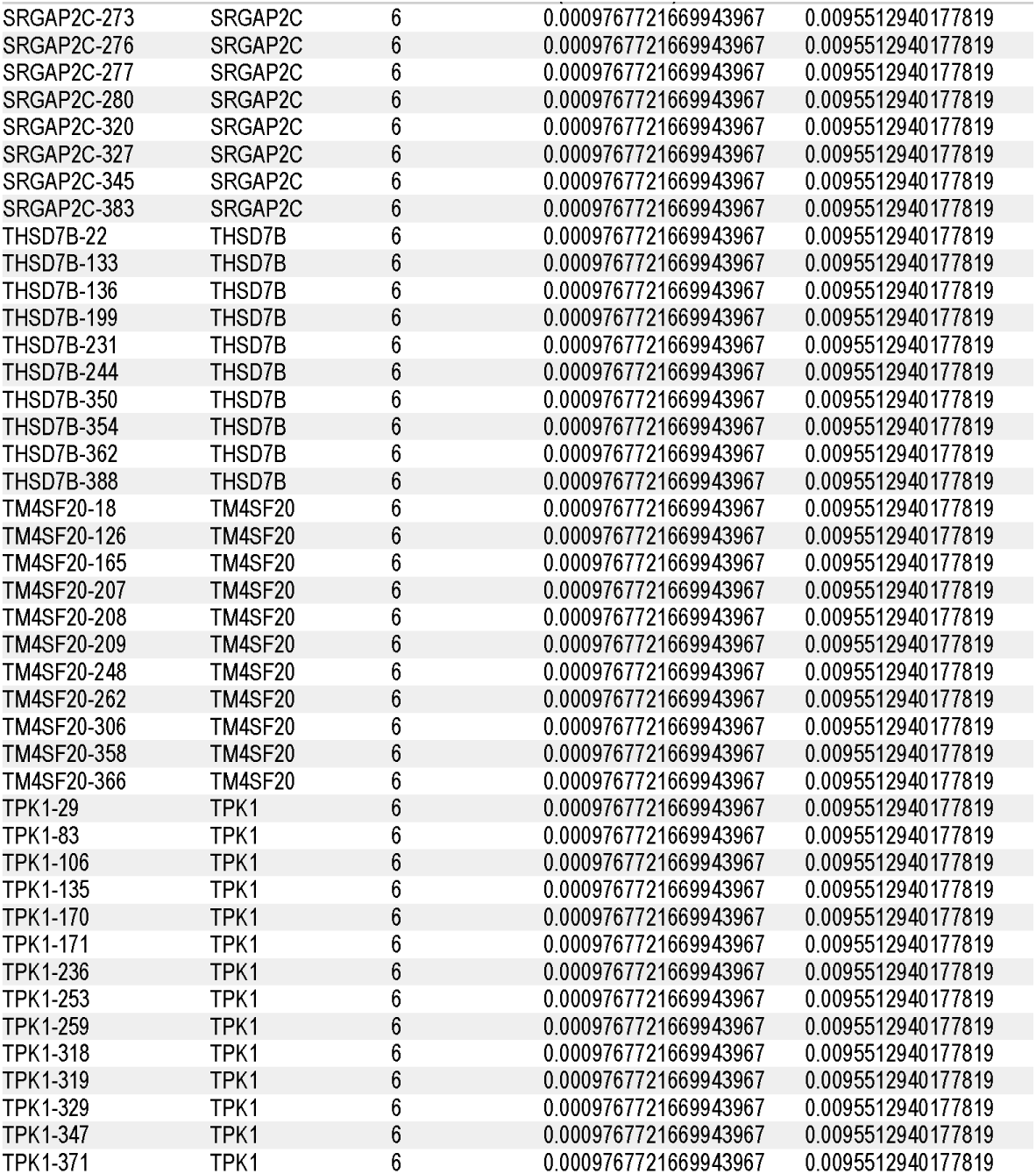
Significantly enriched loci (FDR < 0.05)

A total of 1,372 significantly frequent loci were identified, accounting for 10.23% of all loci. These loci can be further stratified:

• 6–8 occurrences: moderately high frequency;

• 9–13 occurrences: extremely high frequency, appearing across nearly all 13 groups. A stratified hypergeometric enrichment analysis can be performed to examine the distribution of language- and cognition-gene loci across different frequency intervals (see below).

#### 3.4.5 Stratified hypergeometric enrichment analysis by locus frequency

For the three sets of loci derived from Table 16, we conducted two-sided hypergeometric tests separately to determine whether language-/cognition-associated loci were significantly enriched or depleted in each: the first set comprised all 1,372 significant loci with FDR < 0.05; the second set comprised the extremely high-frequency loci with occurrence ≥9; and the third set comprised the moderately high-frequency loci with occurrence 6–8. The background was all 13,415 SNV loci.

The results in Table 17 show that, in the full set of high-frequency loci, language-gene SNVs did not deviate significantly from the background (*P* = 0.41, OR = 0.95, 95% CI 0.84–1.07); for cognition-gene SNVs, a marginal depletion trend was observed (*P* = 0.054, OR = 0.89, 95% CI 0.79–1.00), with the upper bound of the confidence interval exactly equal to 1, not reaching statistical significance. In the extremely high-frequency subset (9–13 occurrences), language genes showed no significant enrichment or depletion (*P* = 0.40, OR = 1.08, 95% CI 0.89–1.30); cognition genes exhibited a sample-level depletion tendency (*P* = 0.09, OR = 0.85, 95% CI 0.71–1.02), with the confidence interval crossing 1. In the moderately high-frequency subset (6–8 occurrences), language genes showed a weak depletion trend (*P* = 0.09, OR = 0.88, 95% CI 0.76–1.02); cognition genes did not differ significantly from the background (*P* = 0.28, OR = 0.92, 95% CI 0.79–1.06). The confidence intervals for all tests above contained 1, and after Benjamini–Hochberg correction for multiple testing, none of the signals met the significance criterion. In summary, although we identified a set of SNVs that recur across multiple similarity intervals (Table 16), we did not obtain robust statistical evidence that this high-frequency locus set is enriched or depleted for language- or cognition-gene-associated SNVs; only sample-level marginal depletion tendencies were observed in some stratified subsets.

**Table 17.**
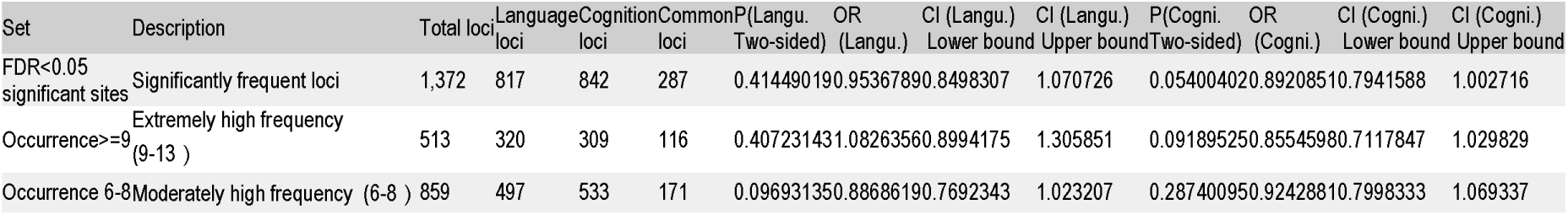
Enrichment tests for language/cognition loci in locus sets of different frequency tiers (two-sided)

#### 3.4.6 Summary of adjacent F_INTERSECTION differences

By comparing the F_INTERSECTION sets of the 13 groups adjacently, a total of 12 differential steps were obtained (Tables 8–9). Overall, the number of newly emerged SNV loci between adjacent groups averaged 657.1, ranging from 209 to 1,152. The largest change occurred from group10 (position 393) to group11 (position 399), with 1,152 newly added loci; the associated genes include all 33 language-cognition genes: *ARHGAP11B; ASPM; ATP2C2; CHRM2; CMIP; CNTNAP2; DAB1; DCC; DCDC2; EP300; FADS2; FLNC; FOXP1; FOXP2; GNB5; GRID2; HAR1A; IGF2R; KIAA0319; KMT2D; MCPH1; NBPF8; NFXL1; NOTCH2NLA; NRXN1; ROBO1; ROBO2; SNAP25*; *SRGAP2*; *SRGAP2C*; *THSD7B*; *TM4SF20*; *TPK1*. Notably, throughout the 355–401 interval, there were three growth peaks in newly added loci, as shown in Figure 10. In addition, in the early steps (355→363 and 363→381), the numbers of newly added loci were relatively low (209 and 395, respectively), and the proportions of language genes and cognition genes were close to each other; while in the later steps (390→393 and 399→400), the proportion of cognition genes increased (Figure 9), suggesting that fine-tuning of cognitive functions may have become more prominent in later evolution. Through a series of enrichment analyses (Tables 10–17), it was found that, at both the gene level and the SNV locus level, the peak positions along the similarity curves did not provide robust statistical evidence for enrichment or depletion of either gene category or their associated SNVs.

## 4 Discussion

The results of this study need to be examined within the broader theoretical context of the language–cognition relationship. The enrichment analysis in this study failed to detect significant enrichment or depletion of either language genes or cognition genes at key evolutionary nodes, with the proportions of the two gene categories in F_Intersection remaining close to background levels throughout. This “negative result” is itself theoretically meaningful: it is compatible with the “language–cognition unity” hypothesis, and does not support treating the two as independent modules that can be clearly separated at the molecular evolutionary level. Of course, this inference requires caution: the categorical division of gene annotation (language genes vs. cognition genes) is itself artificial, and cross-boundary gene functions are the norm (e.g., FOXP2 participates in speech motor control while also influencing cortical development and synaptic plasticity). Therefore, a more accurate formulation is: under the analytical framework of this paper, the evolutionary patterns of language- and cognition-related SNVs did not exhibit distinguishable systematic differences, which suggests that using “language ability” as a proxy indicator for AI cognitive capacity may be theoretically problematic.

In this study, using a redesigned allele presence/absence similarity measure, we successfully detected multiple breakpoints in the later evolutionary stages (rank positions ≥300). These breakpoints are mainly concentrated in primate and human samples, indicating that language-cognition genes underwent accelerated evolution in these lineages. The results from different methods showed high consistency (correlation coefficients >0.9), supporting the robustness of the approach. The PeakRank7 analysis, by comparing the differences between each peak apex and its left neighbour and taking the intersection, revealed that, across consecutive breakpoints (355–401), the proportion of language-gene SNVs gained was overall slightly lower than that of cognition genes (Figure 9). The adjacent F_INTERSECTION difference set analysis further uncovered the newly emerged loci at each step, and these gradually accumulated genetic changes provide temporal clues for understanding the evolution of language and cognitive abilities.

The uniqueness of our method lies in combining traditional allele presence/absence coding with derivative analysis of similarity curves, while incorporating cross-validation across multiple methods and a group-intersection strategy. Unlike commonly used PCA or phylogenetic trees, our approach directly quantifies the degree of sharing between each sample and the reference genome across all SNV loci, avoiding assumptions about population structure. At the same time, we adopted five different distance metrics, covering models ranging from simple matching to allele-frequency-based ones, thereby enhancing the robustness of the results. Similar approaches, such as “genetic similarity sliding windows” and “allele-sharing proportion curves,” have been used to study viral evolution (e.g., mutation accumulation in SARS-CoV-2), but their application to macro-evolution across multiple species has been limited. The rationale of our method is that the appearance of derivative peaks marks a transition from slow to rapid similarity increase, which typically corresponds to the acquisition and fixation of numerous new alleles, rather than just single-site substitutions. Therefore, using steep slopes as a measure of multi-SNV pattern changes can better reflect evolutionary events at the genomic level than point mutations alone [39–43].

The steep slopes and small plateaus on the similarity curve constitute a “fingerprint” of the evolutionary process. Although steep slopes may partly arise from sampling bias (e.g., lack of genomic data for representative intermediate species), we still consider them to have substantial quantitative value. First, the slope of a steep segment reflects the change in similarity per unit ranking step; this value is not affected by absolute species richness, so it can serve as a relative indicator of evolutionary rate. Second, even under insufficient sampling, the appearance of steep slopes suggests that significant genetic clustering already exists among the available samples, and these clusters often coincide with known evolutionary events (e.g., speciation or migration). Small plateaus represent intervals of relatively stable similarity, possibly corresponding to extended periods of ecological niche conservatism or genetic drift equilibrium. It is noteworthy that these small plateaus are not merely the result of redundant sampling of the same species, because our dataset is dominated by different species; yet plateaus still emerge, suggesting that multi-species evolution in nature may automatically produce a pattern of “punctuated equilibrium”—i.e., long-term stasis alternating with short-term rapid change. Similar phenomena have long been described in the fossil record (Gould & Eldredge, 1977) [44], and at the molecular level, studies have also reported synchronous allele-frequency changes across multiple species during specific periods (e.g., “species radiation” events) [44–48]. Our results suggest that the evolution of multi-SNV patterns may similarly follow a punctuated equilibrium pattern, where small plateaus represent conserved regions with strong functional constraints, and steep slopes represent bursts of innovation when constraints are relaxed.

In our results, the peak apex samples with the highest frequencies are c25 (China (WGM70) PRJEB36297), c27 (HJTM115, China, PRJEB36297), and ja2 (Funadomari23 female-2, PRJDB7235), all of which are archaic hominin samples from East Asia. Previous studies have shown that c25 and c27 are genetically associated with Denisovans [49–50]. Denisovans are an enigmatic archaic hominin group discovered in Siberia and East Asia, whose genomes have made significant contributions to the immune systems and high-altitude adaptations of modern humans. Our findings suggest that the evolution of language and cognition genes was not confined to an African “origin of *Homo sapiens*” scenario, but rather that important adaptive changes occurred independently in Eurasia. In light of the literature background of these high-frequency samples, we hypothesize that environmental conditions in East Asia may have facilitated rapid evolution of archaic hominin cognition-gene polymorphism patterns, thereby accumulating a relatively large number of language- and cognition-related mutations. This hypothesis is consistent with recent ancient DNA studies (e.g., Fu et al., 2013) [51–53], indicating that East Asia was one of the important evolutionary centres for modern humans. The substantial number of mutations accumulated in East Asia (Figure 8) likely occurred after early modern humans migrated out of Africa, although this will need to be carefully examined and confirmed in future research.

On the horizontal axis of the similarity curve, in the direction from 300 to 410, the first major peak (around 355) corresponds to sample mo2 (Iberomaurusian, Morocco, PRJNA422662), an archaic hominin from North Africa, which is consistent with the conclusion that modern humans originated in Africa. Subsequently, other African samples such as sf4 (flo023, South Africa, PRJEB98562) appear at peak positions, but the majority of peak samples come from Eurasia (e.g., c25, c27, ja2) and primates (p21, p5, p9, etc.). Notably, two archaic hominin samples from Yunnan, China, associated with Denisovans—xy1 (Xingyi_LN, Yunnan, PRJCA015361) and xy2 (Xingyi_BA, Yunnan, PRJCA015361)—frequently appear near the peaks. This may be related to Yunnan’s unique geographical location: situated at the southeastern edge of the Qinghai–Tibet Plateau, it possesses complex karst landforms that may have served as refugia for archaic hominins during glacial periods, thereby facilitating the accumulation of genetic diversity and mutations in language-cognition genes. Such a pattern of geographic isolation and diversification is consistent with the “refugium theory” and provides spatial evidence for the special role of East Asian archaic hominins in the evolution of language and cognition. Overall, the sequence of peaks from Africa to Eurasia may reflect multiple independent episodes of language-cognition gene intensification in different regions of Eurasia after modern humans migrated out of Africa. In addition, we need to pay special attention to sample xy2, as it appears late on the horizontal axis of the similarity curve in Figure 1, lying in close proximity to sf4 and cy1 (Canary Islands El Teide National Park, Morocco, PRJEB61655), with both being only one step away from pp6. However, these three samples originate from South Africa, Morocco, and Yunnan (China), respectively [54–58], which readily invites the following speculation: owing to food shortages, archaic humans may have migrated westwards from the glacial karst caves of Yunnan, moved south along the west coast of Africa to Morocco, and finally reached South Africa. Whether xy2 belongs to the same group of archaic hominins reported in the literature [59–60] is a question worthy of future investigation.

From the first derivative peak (Figure 2, located at the African archaic hominin sample mo2) onwards, the middle portion consists largely of Eurasian samples, but the final part still falls on African samples (Figures 3, 7–8). This implies that the African archaic hominins closest to modern humans either evolved locally, or that archaic hominins that evolved in Eurasia—especially in East Asia—returned to Africa in some way. If in the future we persistently fail to find sufficient African (archaic) human samples whose positions on Figures 1 and 2 lie between the first derivative peak and the last derivative peak, then a hypothesis would automatically emerge, namely that archaic hominins such as the first peak sample mo2 could directly evolve, bypassing many intermediate steps, into the African archaic humans closest to pp6 on the far right of Figure 1. Clearly, such a hypothesis would not hold, so we would have to consider an alternative hypothesis: that archaic hominins (erectines and early Homo sapiens) that evolved early in Africa migrated out of Africa and dispersed globally, after which some underwent profound evolution in Eurasia under specific environmental conditions and then returned to the African continent, eventually evolving into African archaic hominins very close to modern humans, such as sf1-6 (late Homo sapiens migrating out of Africa again). This hypothesis will need to be rigorously tested in the future by incorporating a large number of African archaic hominin samples. Of course, if all Eurasian archaic hominin samples are derived from erectines or early *Homo sapiens* that left Africa early (not limited to late Homo sapiens that migrated out of Africa), then it would be easy to explain why the peak samples in the 370–400 interval of the similarity curve are predominantly composed of Eurasian archaic hominins.

The series of PeakRank7 analyses show that the proportions of language genes and cognition genes in F_INTERSECTION are generally close, but language genes are slightly lower in most peaks. This pattern of “close overall but with minor differences” suggests that language and cognitive functions are highly coupled in evolution. From the F_INTERSECTION line plots, it can be seen that the proportion of language genes has two rising peaks in the early (355–363) and middle (381–390) stages, whereas cognition genes show a greater proportional increase in the later stage (393–401), which may reflect a transition from “acquisition of language abilities” to “fine-tuning of cognitive functions.” At the gene level, we counted the occurrence frequencies of each gene across multiple F_INTERSECTION sets and found no genes with exceptionally high or low frequencies; it appears that all 33 genes evolve along the evolutionary timeline at roughly similar frequencies. This mechanism may provide a new framework for understanding the evolution of human intelligence.

This study has several limitations. First, the SNV dataset used is mainly derived from public databases, and sample coverage is uneven across taxonomic groups, which may lead to overestimation or underestimation of certain breakpoints. Second, we relied solely on binary coding of allele presence/absence, ignoring information such as allele frequencies and linkage disequilibrium, potentially losing some fine-scale genetic signals. In addition, the annotation of language and cognition genes depends on existing literature, which may involve omissions or misclassifications, and we did not distinguish the effects of gene regulatory regions or non-coding regions. Future work will progressively address these design issues.

The findings of this study raise a deeper question: if language and cognition are difficult to separate at the molecular evolutionary level, is the current practice in large language model-based AI research of equating “language ability” with “cognitive ability” methodologically valid? Recent studies from MIT and other institutions have indeed shown that language networks exhibit robust responses to sublexical regularities [61], but this precisely supports a more conservative interpretation: what language networks process are the structural constraints of a symbolic system, rather than general-purpose cognitive reasoning. If the phenomenon of language loss with preserved cognition is further confirmed experimentally, then directly interpreting the “linguistic performance” of LLMs as “cognitive ability” would require more cautious theoretical argumentation [62]. Future research should, on the basis of this framework, incorporate more comparative data from non-linguistic cognitive tasks, as well as more molecular data from individuals with language disorders, in order to test the predictive differences between the “speaking-cognition” and “language-cognition” frameworks.

## 5 Conclusion

In this study, we developed a multi-species SNV similarity analysis framework based on binary allele presence/absence features and applied it to SNV data from 33 language-/cognition-related genes, spanning species from fish to humans. The main conclusions are as follows: (1) The similarity curves generated by the five distance measures were highly consistent, with major peaks concentrated at positions 355, 363, and 381. These breakpoints correspond to archaic hominin and primate samples, indicating that language-cognition genes underwent significant mutation accumulation at these evolutionary nodes. (2) Using a group-intersection strategy, the PeakRank7 analysis revealed shared differential loci across 13 consecutive breakpoints. The average proportion of language genes in these shared sets was higher than that of cognition genes, and the early breakpoints (355–363) showed the largest number of newly added loci, with language genes contributing prominently. (3) The high-frequency peak apex samples were predominantly East Asian archaic hominins (c25, c27, ja2), suggesting that Eurasia, especially East Asia, played a unique role in the evolution of language-cognition genes. (4) The steep-slope and plateau pattern of the similarity curves reflects punctuated equilibrium at the macro-evolutionary level, while the adjacent F_INTERSECTION difference sets revealed a stepwise acquisition of new loci. (5) All 33 language-cognition genes appeared at the 13 peaks, implying that the acquisition of language and cognitive abilities is the integrated outcome of repeated mutations, adjustments, and evolution of thousands of SNV loci. However, these peaks did not show enrichment for either language genes or cognition genes, nor for their associated SNVs at the locus level. In addition, among the 33 genes, the six that contributed the most newly gained loci across the 13 peaks were ARHGAP11B (cognition, 5.55% of contributed loci), HAR1A (cognition, 5.53%), KIAA0319 (cognition, 4.94%), ATP2C2 (language, 4.85%), CMIP (language, 4.53%), and DCDC2 (language, 4.5%). This preliminary observational analysis presents, from a unique perspective, an evolutionary profile of the polymorphism patterns across a total of 13,415 SNV loci from language and cognition genes.

From a broader perspective, this study offers a methodological caution for AI foundations research: using “language ability” as a proxy indicator for cognitive capacity may overlook the fundamental distinction between language as an internal symbolic computational system and language as an externalized communication tool. The molecular evolutionary data in this study failed to support the separability of language genes and cognition genes at key evolutionary nodes, which is compatible with the theoretical framework of “language–cognition unity.” Based on this framework, future artificial intelligence research may need to more carefully distinguish whether the capabilities exhibited by a model constitute mastery of the deep rules of a symbolic system (closer to I-language) or merely fitting to surface statistical patterns of language (closer to the imitation of externalized behavior). This distinction holds foundational theoretical value for understanding the cognitive significance and fundamental limitations of large language models.

## Acknowledgments

This study was supported by a State Language Commission Research Grant (YB135-117) and National Research Center for Foreign Language Education Grant (ZGWYJYJJ10A042). The authors are indebted to Shuaiyu Zhang for helping in SNV-searching software development.

