## Supplementary figures and images for "From Language-Cognition Coupling to AI Foundations: An Evolutionary Perspective on 13,415 SNVs in 33 Language/Cognition Genes"

### Hellinger.png

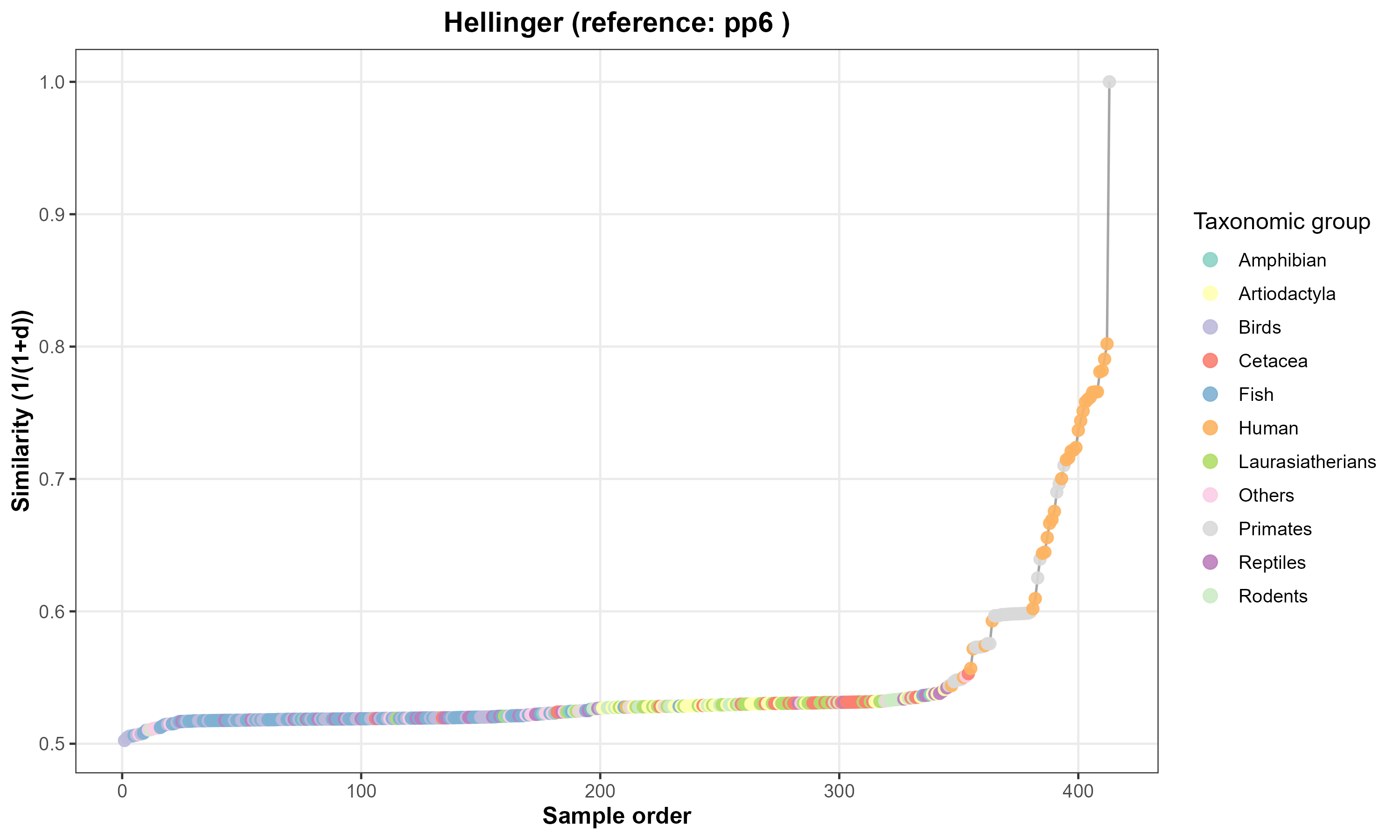

### Hellinger.png

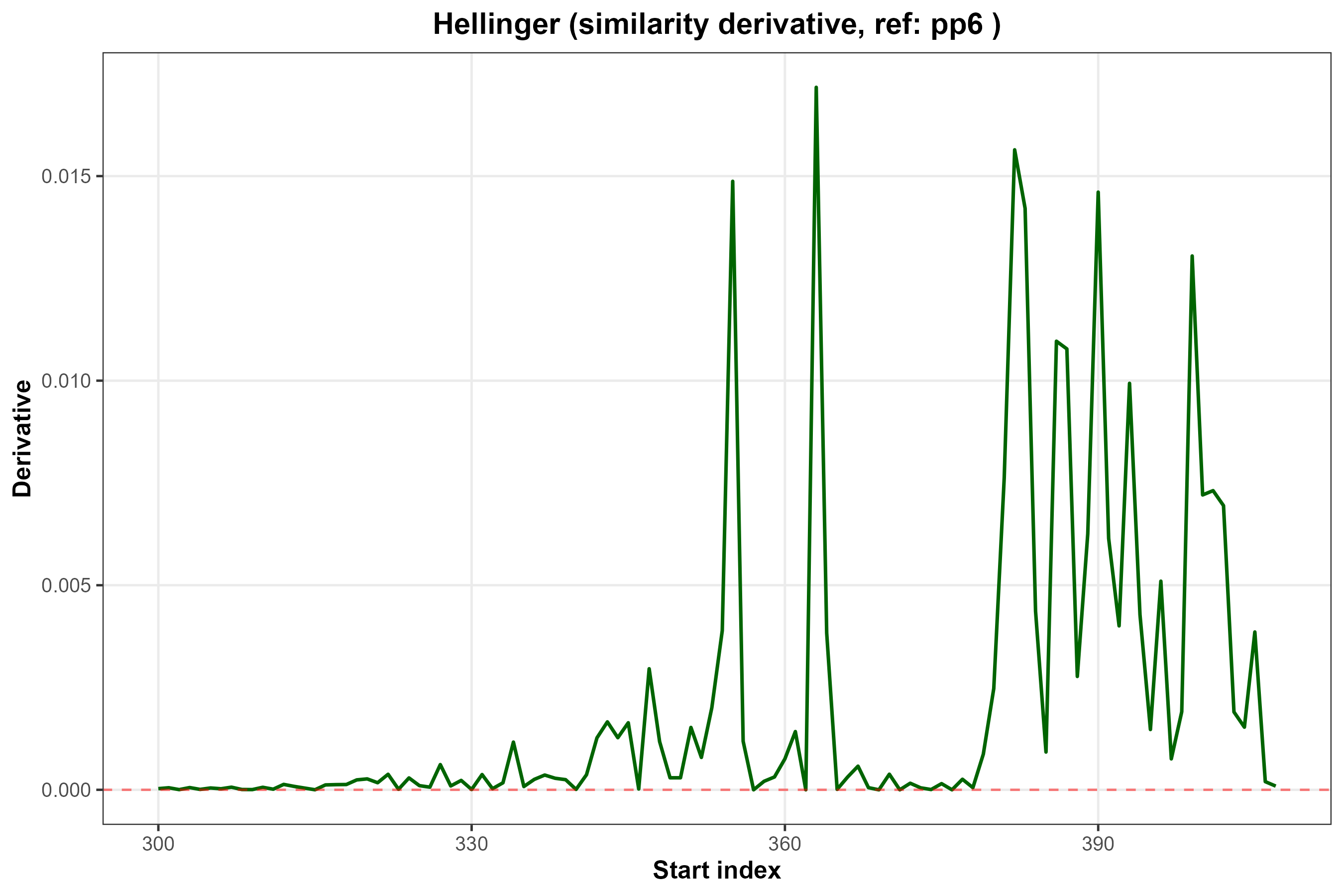

### Hellinger_combined.png

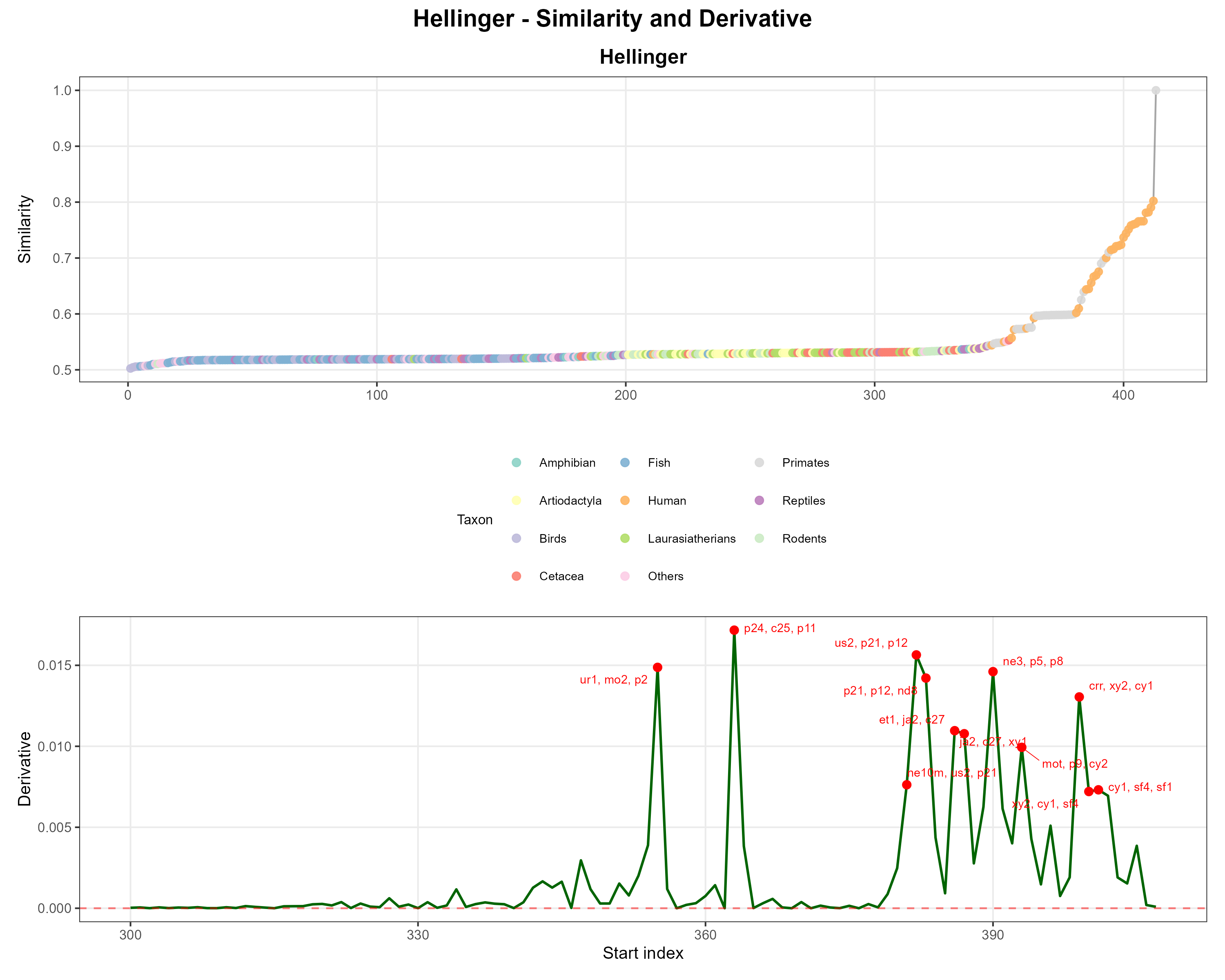

### Nei.png

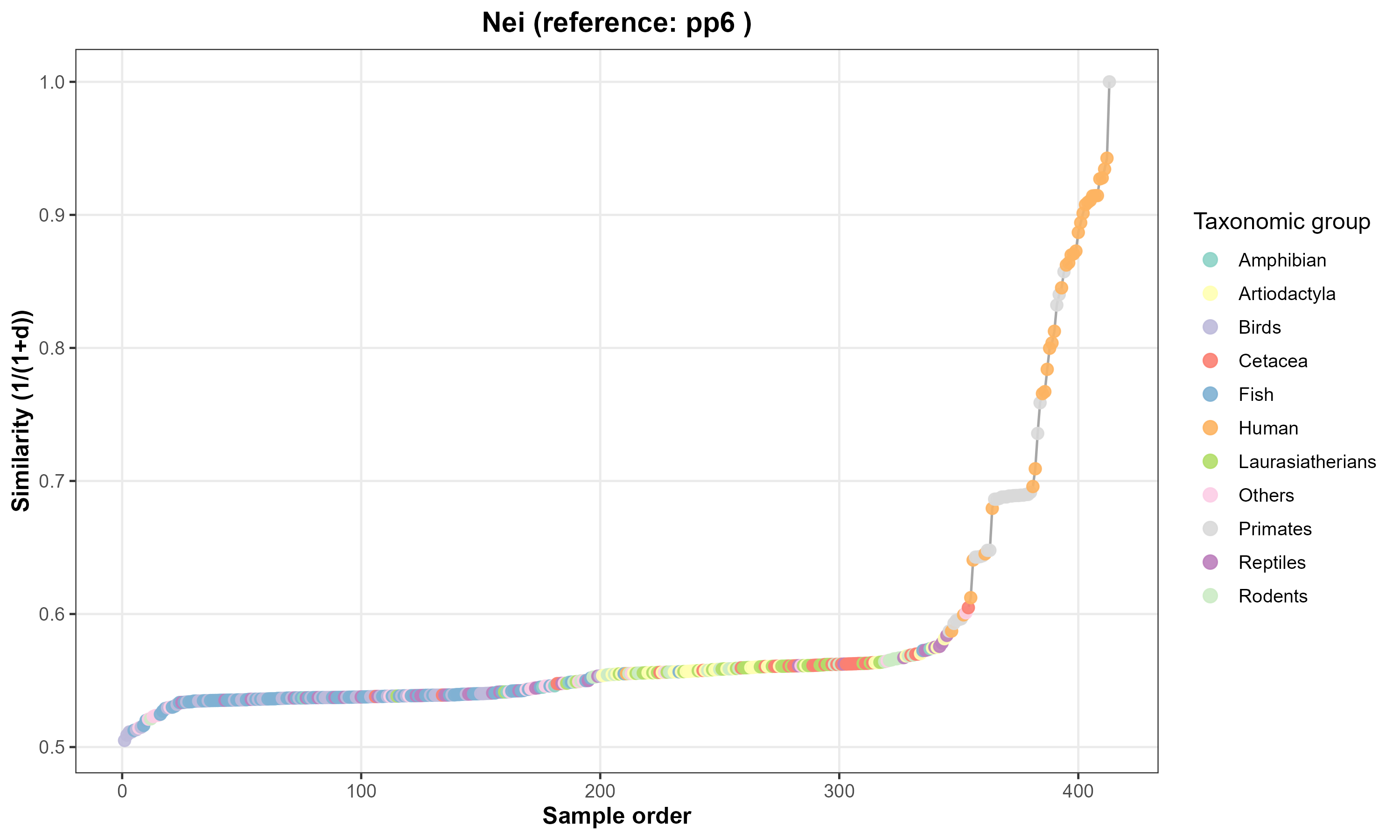

### Nei.png

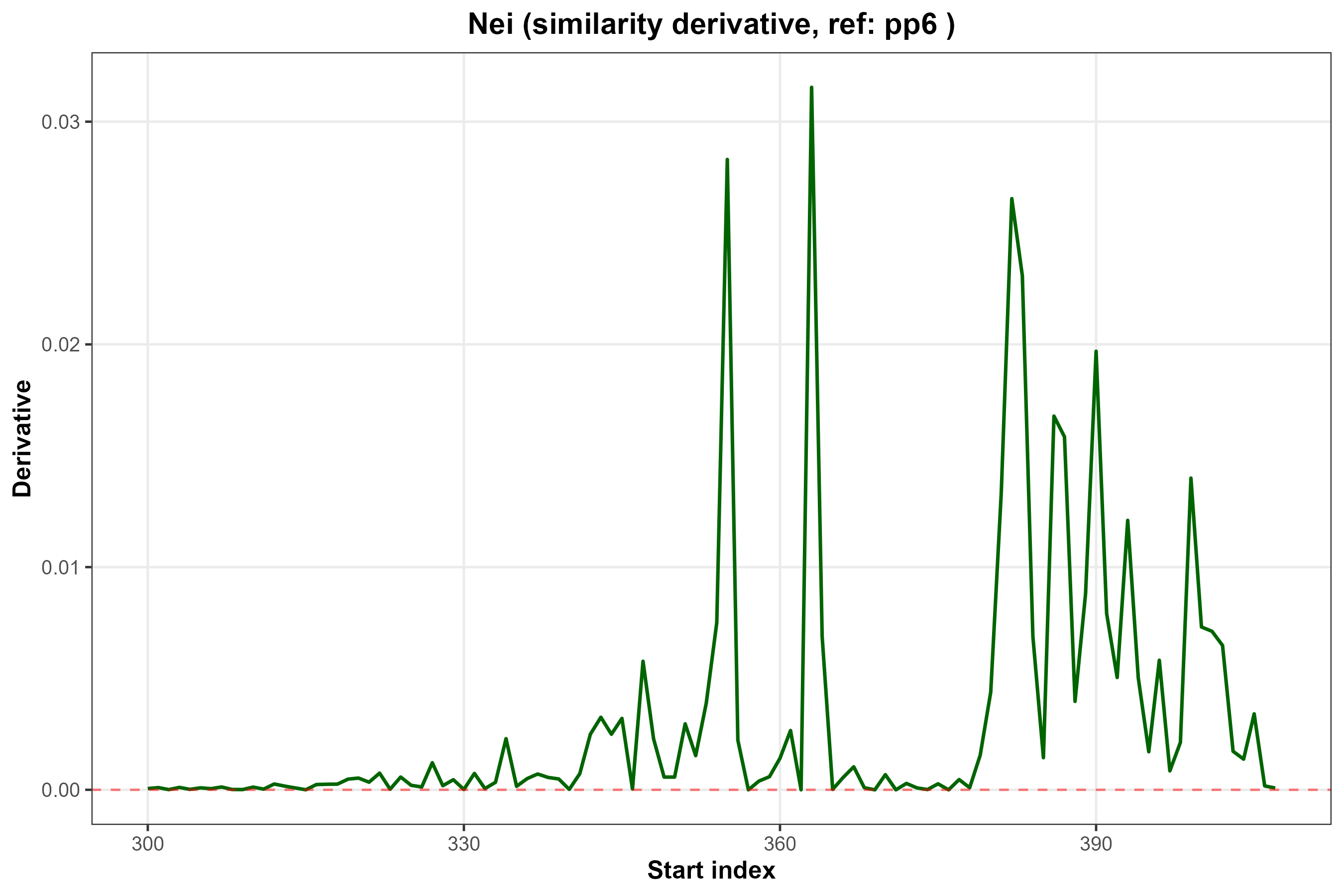

### Nei_combined.png

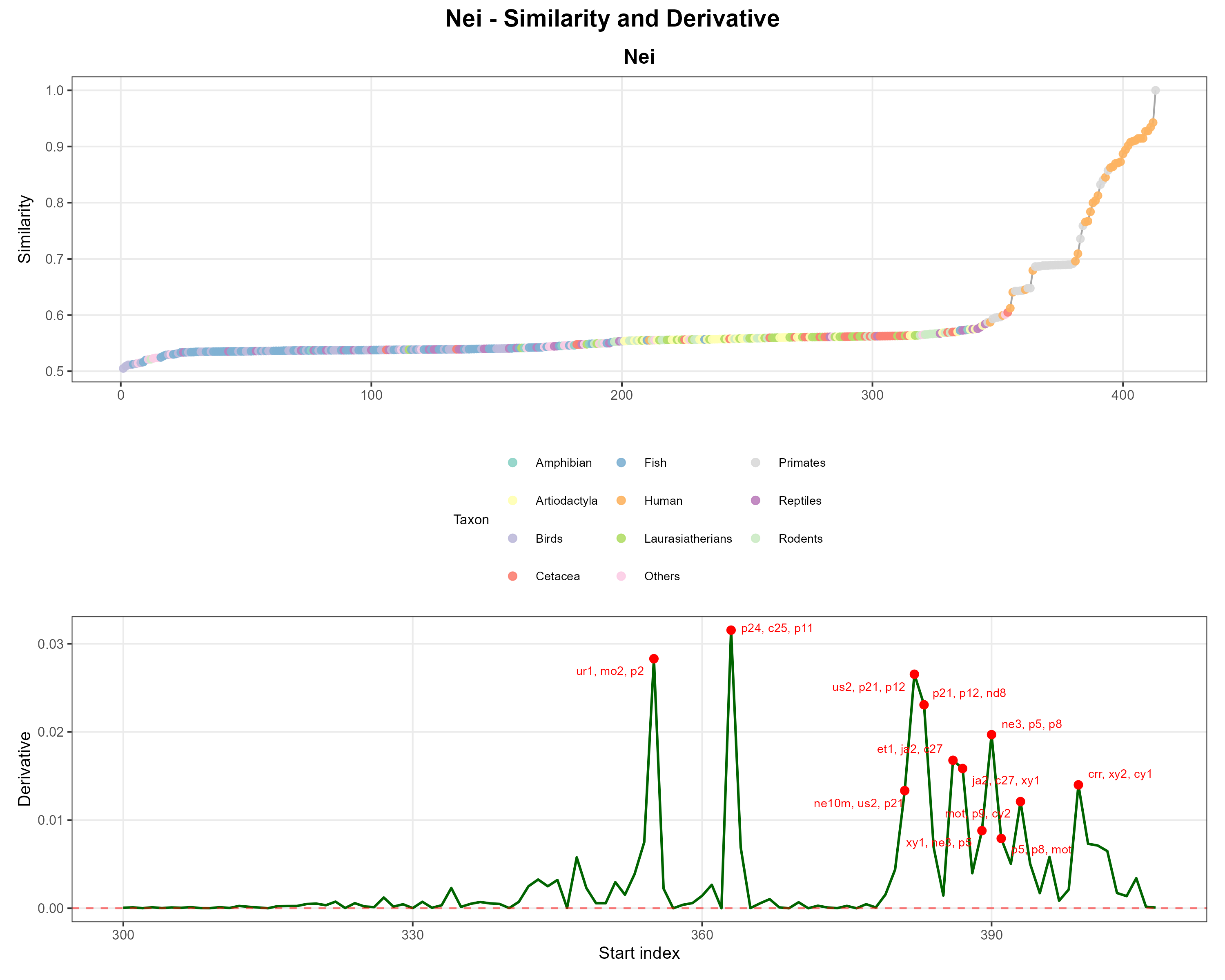

### Reynolds.png

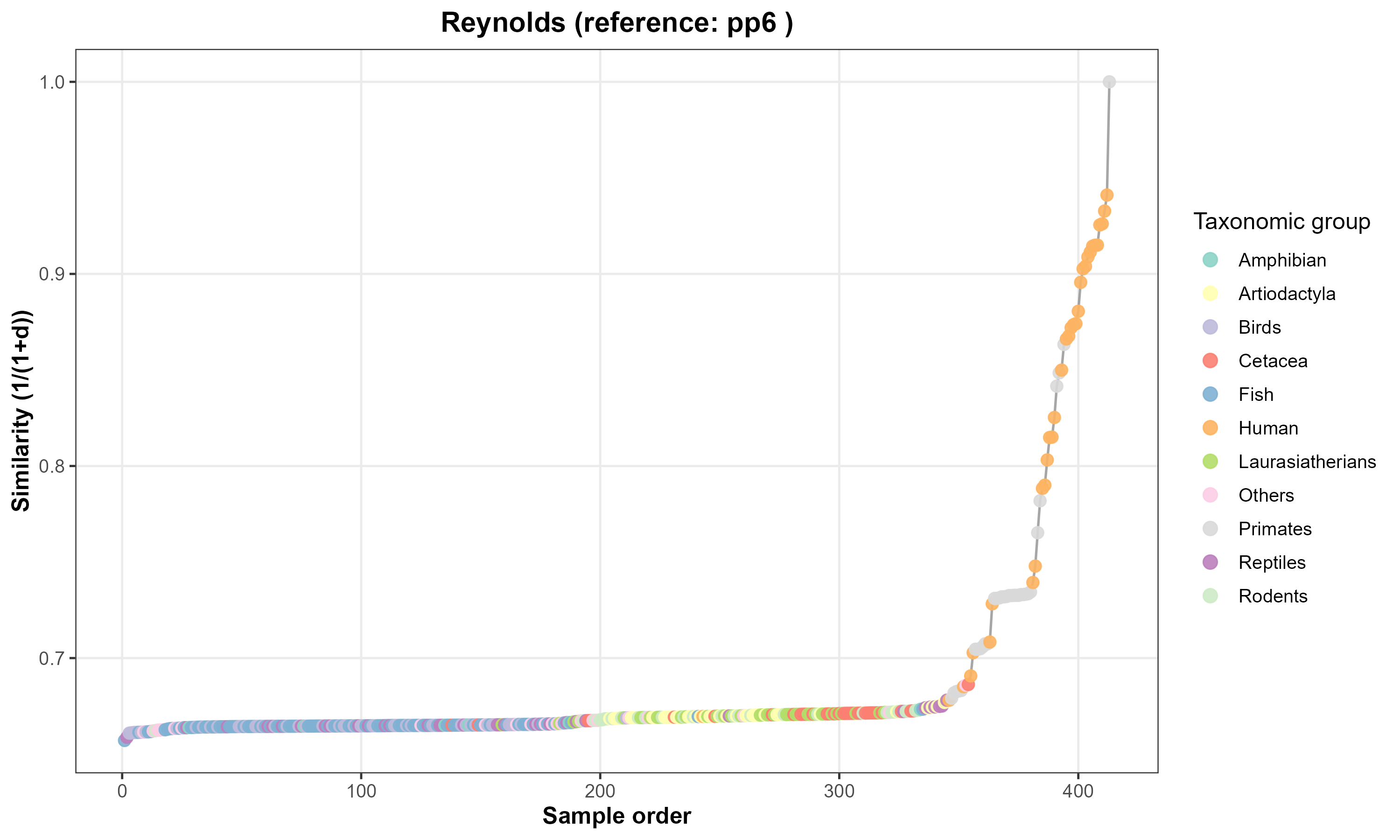

### Reynolds.png

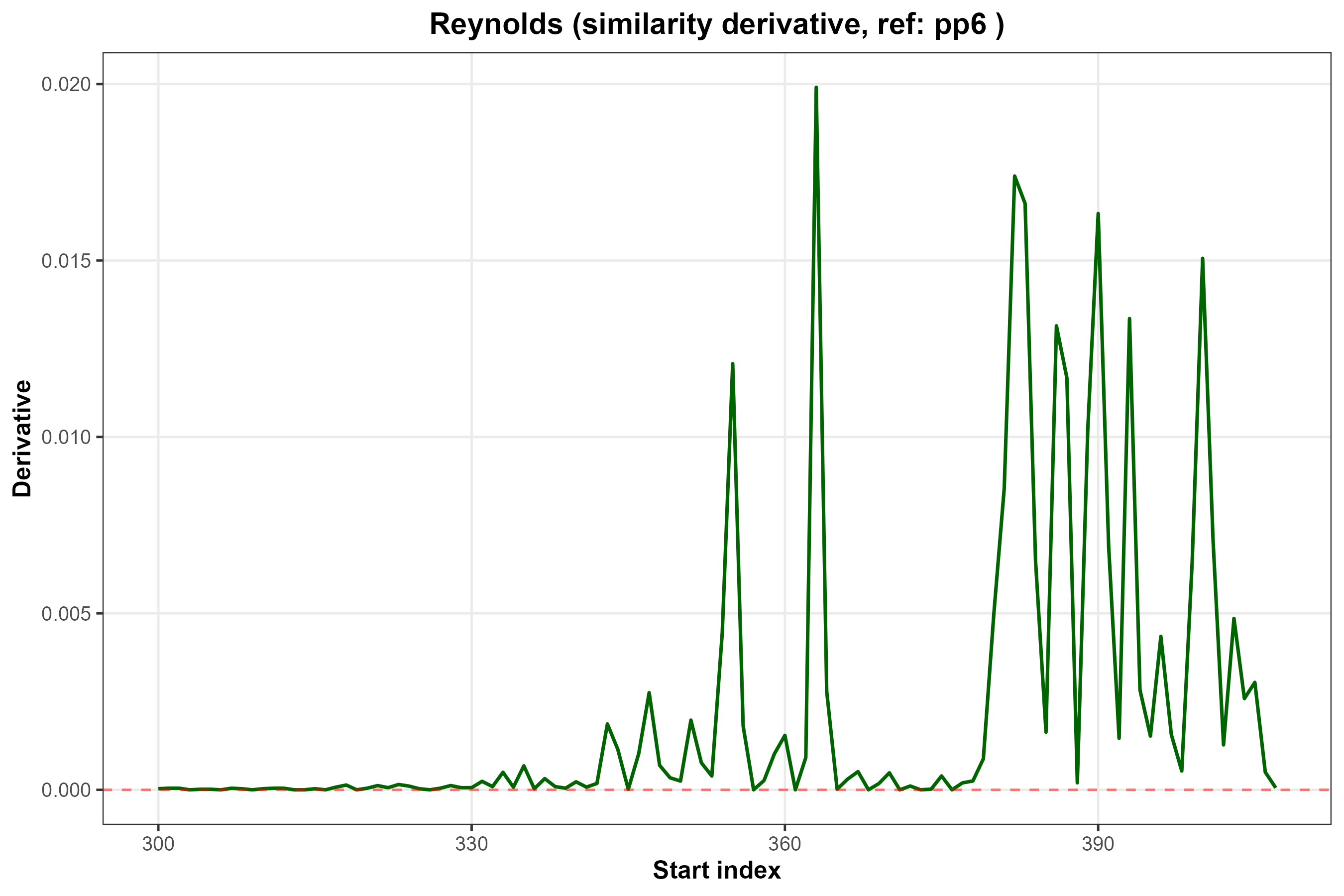

### Reynolds_combined.png

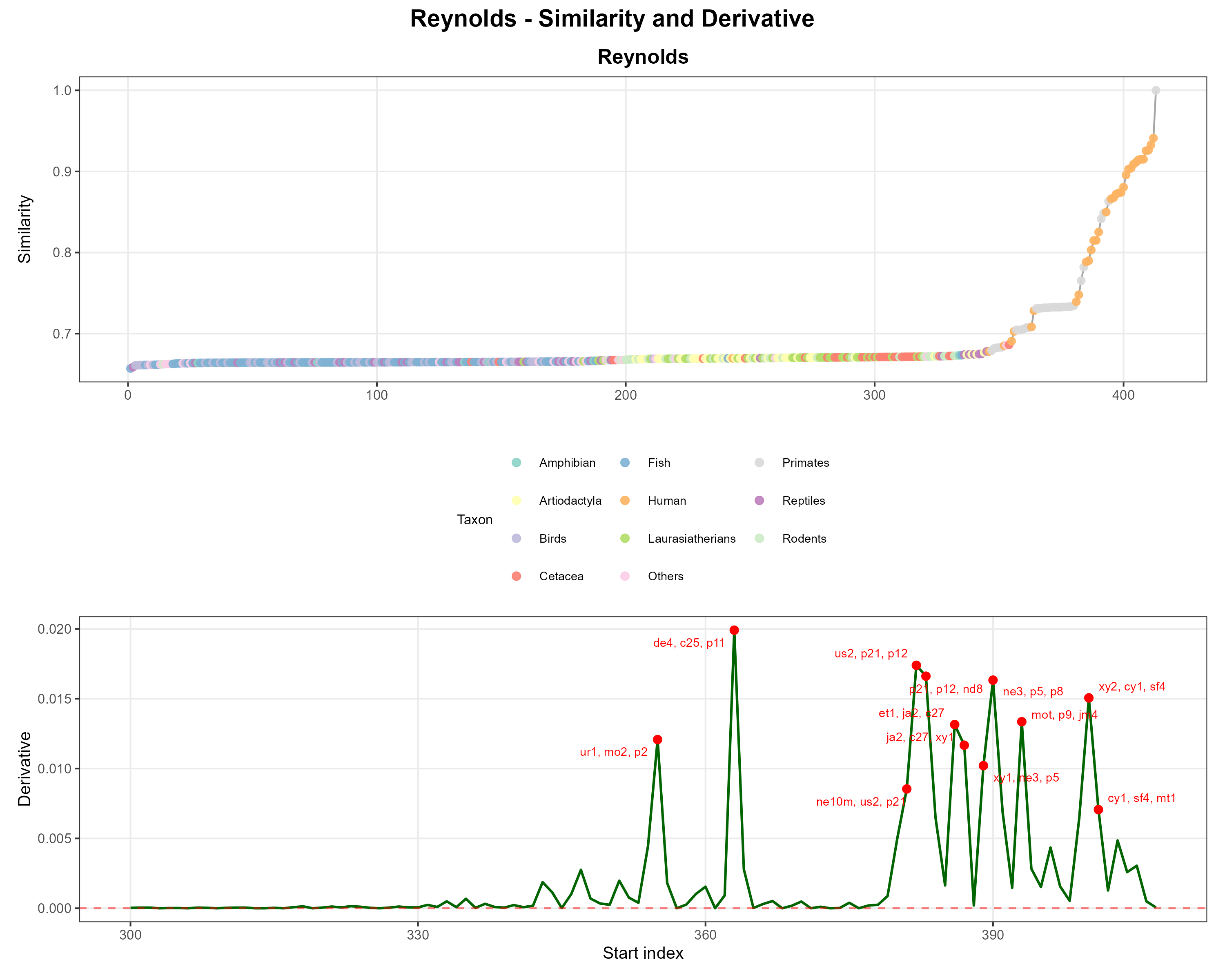

### Rogers.png

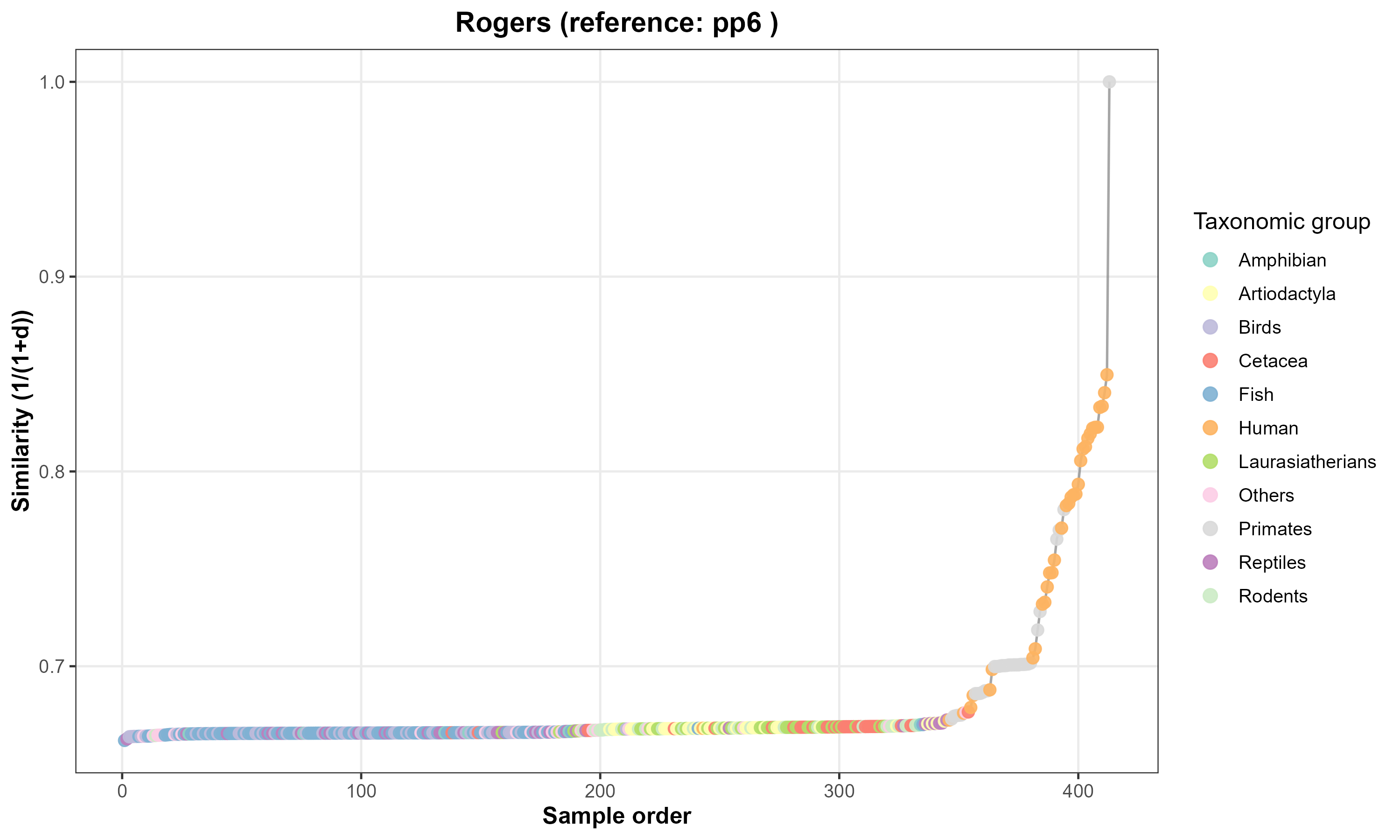

### Rogers.png

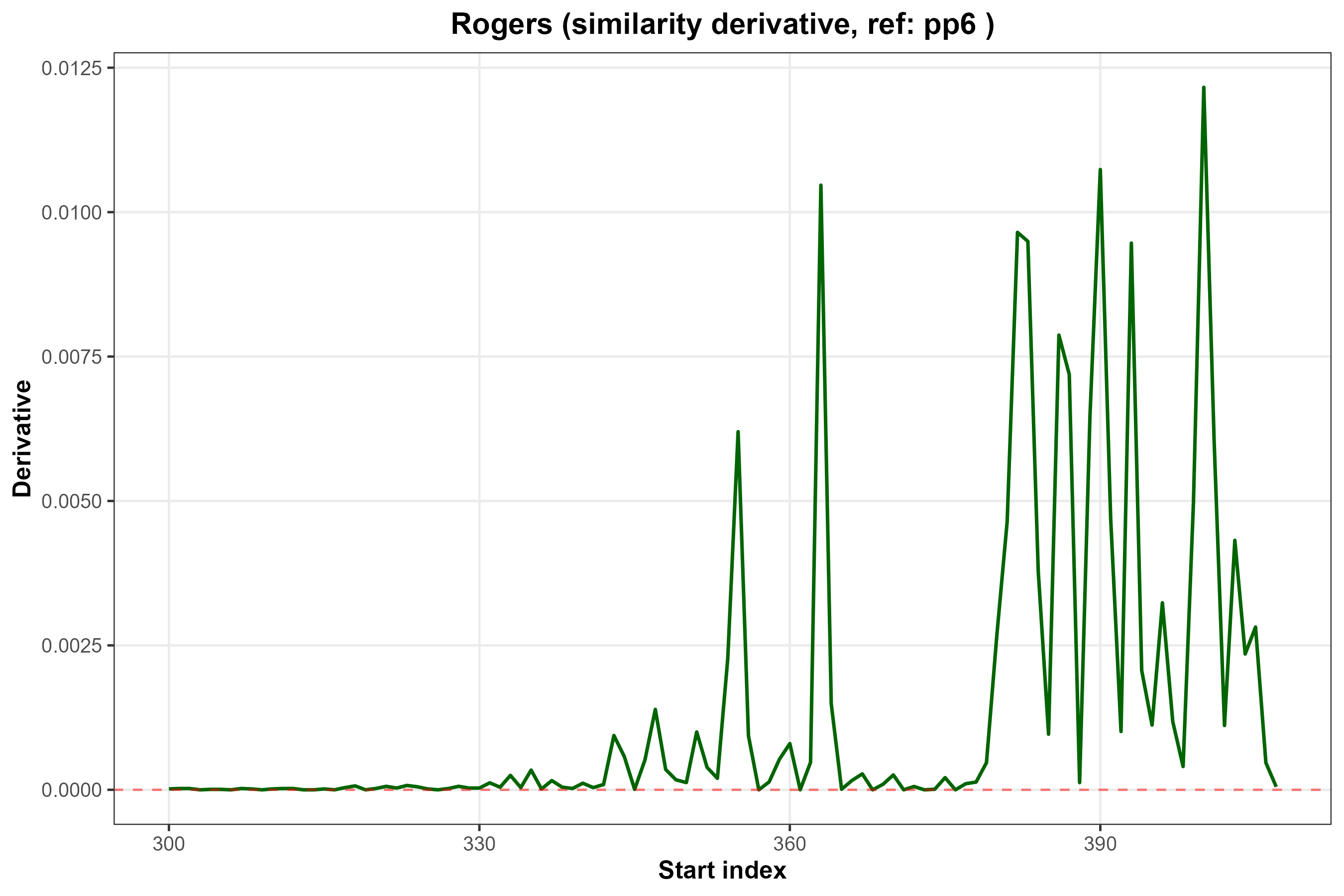

### Rogers_combined.png

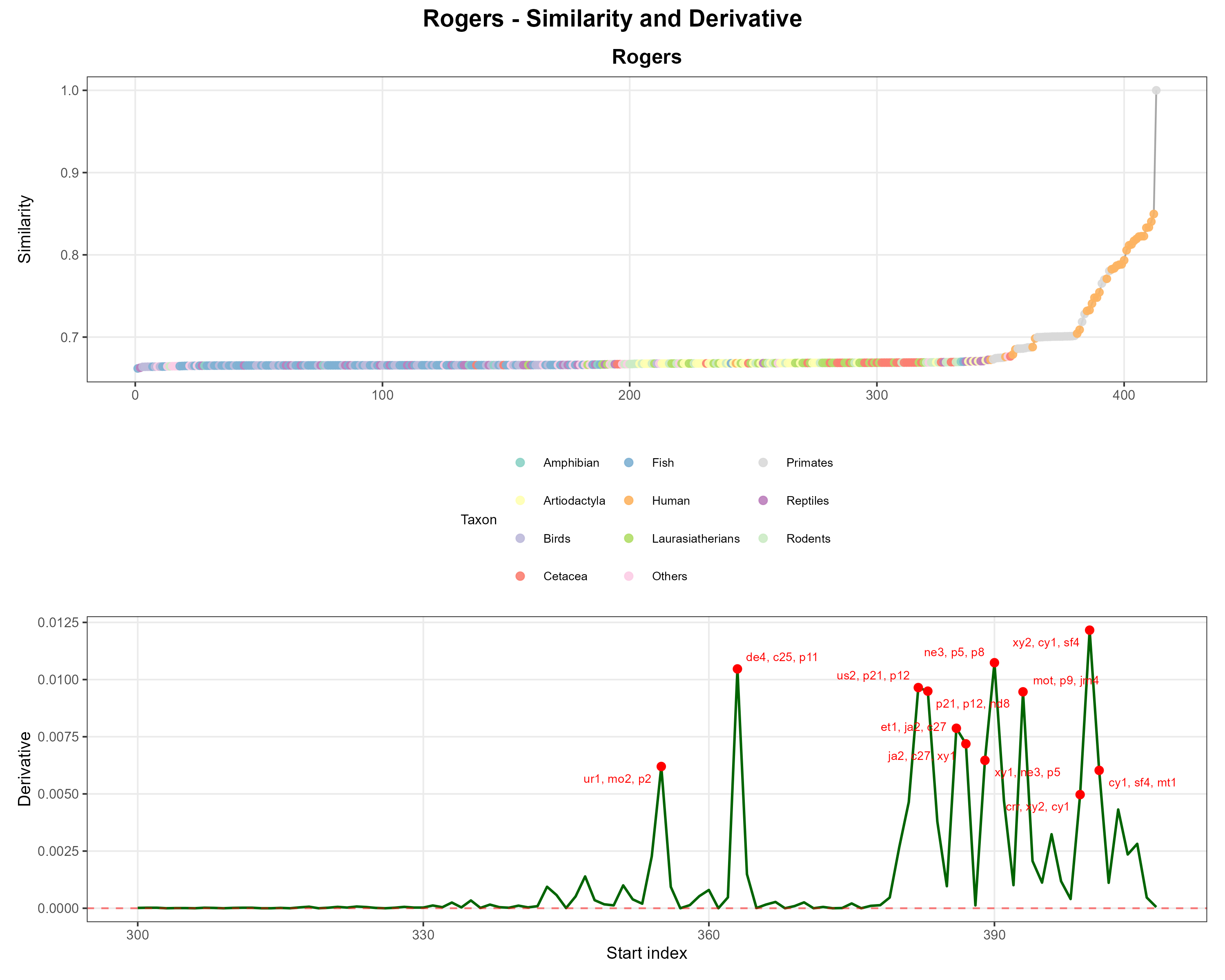

### Sørensen.png

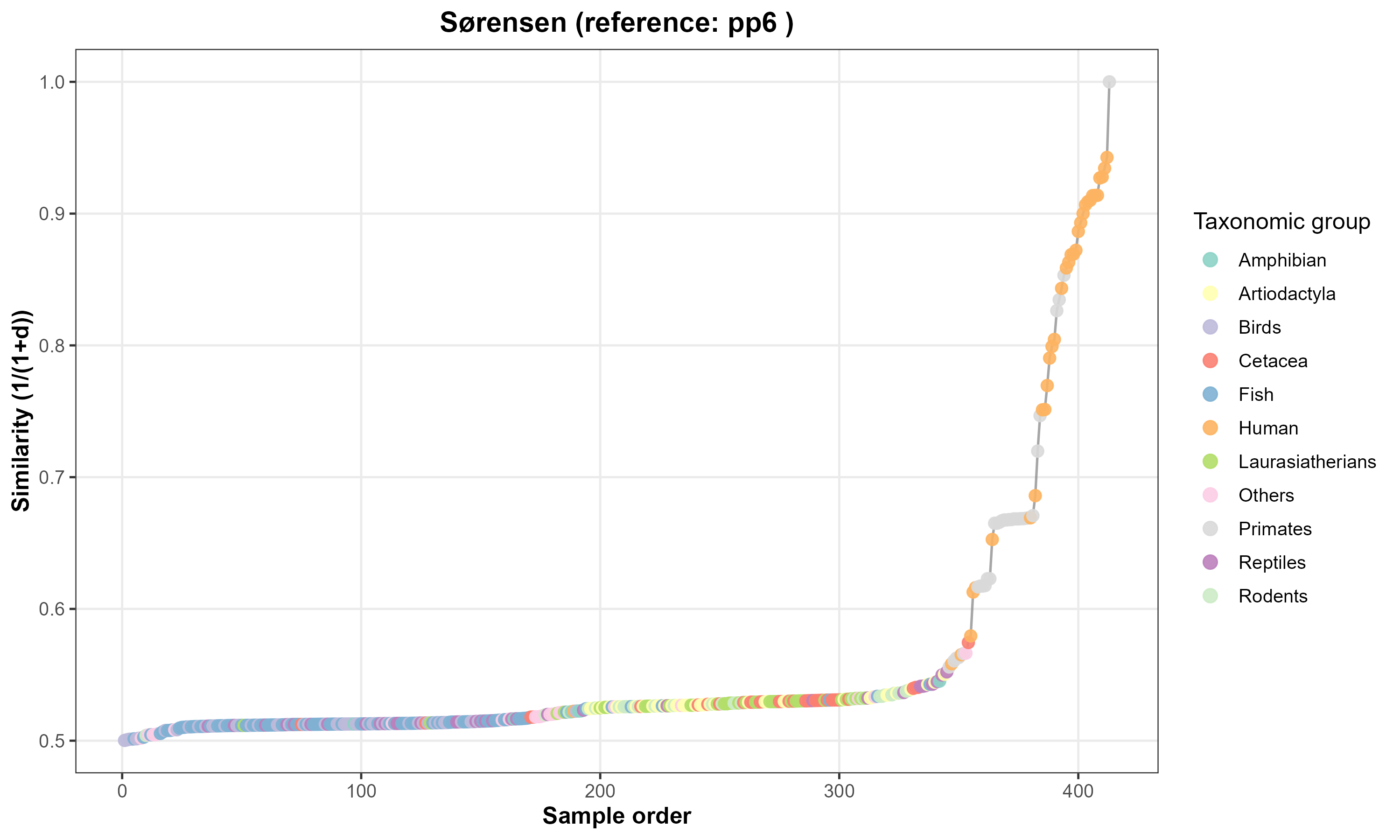

### Sørensen.png

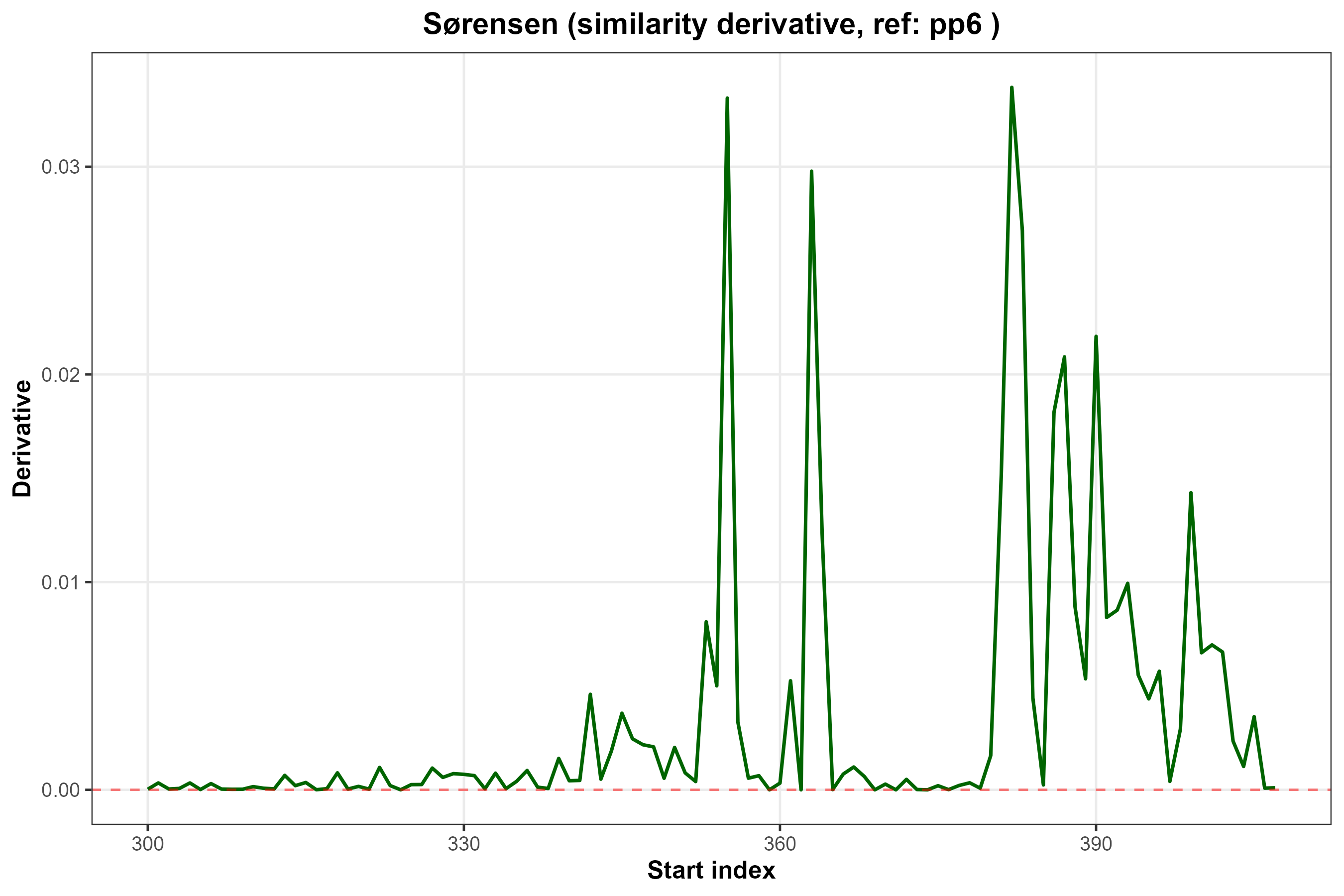

### Sørensen_combined.png

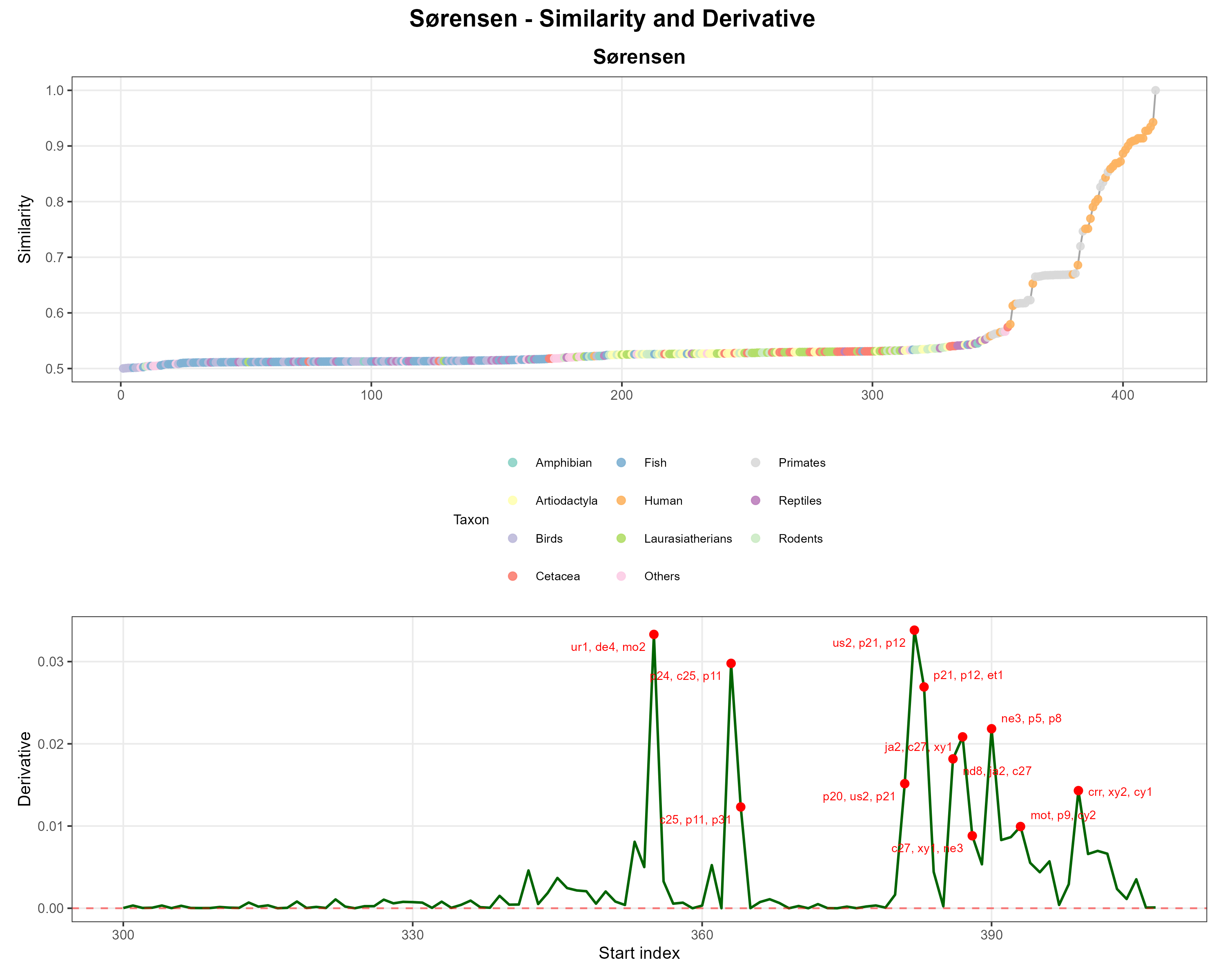
